# The relationship between sexual dimorphism and intersex correlation: do models support intuition?

**DOI:** 10.1101/2024.11.29.626061

**Authors:** Gemma Puixeu, Laura Katharine Hayward

## Abstract

That a high genetic correlation between the sexes (*r*_*fm*_) constrains the evolution of sexual dimorphism and that they should negatively correlate with one another, are assumptions commonly made in the field of sex-specific adaptation. While some empirical observations support a general negative relationship, the mechanisms underlying this pattern and the conditions under which it arises are poorly understood. Concretely, two primary hypotheses are often invoked: first, that traits with ancestrally low *r*_*fm*_ are less constrained in their ability to respond to sex-specific selection and thus evolve to be more dimorphic; second, that sex-specific selection acts to reduce *r*_*fm*_. However, no model to date has formalized these hypotheses and tested the conditions in which they hold. Here, we develop models of sex-specific stabilizing selection, mutation and drift to explore various scenarios potentially generating a negative correlation between intersex correlation and sexual dimorphism, with a focus on testing the common hypotheses. We recover the classical result that, with an infinite population size, *r*_*fm*_ and sexual dimorphism are independent at equilibrium. Further, we show that this independence is maintained with a finite population; and this is in spite of the fact that, as we demonstrate, genetic drift generates nonzero sexual dimorphism even when selection between the two sexes is identical. Moreover, we demonstrate that the two common hypotheses only imply a negative association if additional assumptions are made. Specifically, that 1) some traits are sex-specifically adapting under directional selection, and 2) this sex-specific adaptation favours increased dimorphism more often than decreased dimorphism. These results provide, to our knowledge, the first mechanistic framework for understanding the conditions under which a negative correlation between intersex correlation and sexual dimorphism may arise. They also offer a compelling explanation for the inconsistent empirical evidence observed in nature, highlighting the importance of context-specific factors in shaping this relationship.

## 1 Introduction

That a high correlation between the sexes (*r*_*fm*_) constrains the evolution of sexual dimorphism and that both should negatively correlate with one another are common assumptions in the field of sex-specific adaptation (Lande, 1980, 1987; Bonduriansky & Rowe, 2005; Fairbairn, 2007; Poissant et al., 2010; Stewart et al., 2010). These assumptions are supported by evidence that traits that are more sexually dimorphic have lower *r*_*fm*_ values. This trend, although far from universal (Cowley & Atchley, 1988; Preziosi & Roff, 1998; Chenoweth & Blows, 2003; Ashman & Majetic, 2006; Leinonen et al., 2011; Puixeu et al., 2019), has been described across traits and species (e.g. Delph et al., 2004, 2010; Bonduriansky & Rowe, 2005; McDaniel, 2005; Fairbairn, 2007; Poissant et al., 2010; Griffin et al., 2013; Cox et al., 2017). Two hypotheses are most commonly proposed as potential explanations for this pattern (stated in e.g. Bonduriansky & Rowe, 2005; Fairbairn, 2007; Griffin et al., 2013; Stewart & Rice, 2018; McGlothlin et al., 2019): first, that traits with ancestrally low *r*_*fm*_ are less constrained in their ability to respond to sex-specific selection and thus evolve to be more dimorphic (as discussed in, for example, Bolnick & Doebeli, 2003; Poissant et al., 2010; Stewart et al., 2010); second, that sex-specific selection acts to reduce the *r*_*fm*_ (Lande, 1980; Bonduriansky & Rowe, 2005; Bonduriansky & Chenoweth, 2009; McGlothlin et al., 2019).

In line with the first hypothesis is the idea that sexual dimorphism will easily (hardly) evolve for traits with a low (high) intersex correlation (Stewart et al., 2010; Stewart & Rice, 2018). The potential for a high intersex correlation to pose a long-term constraint on the evolution of sex differences has been illustrated by some artificial selection experiments (Harrison, 1953; Reeve & Fairbairn, 1996; Stewart & Rice, 2018). Most notably, Stewart and Rice (2018) observe a minimal change in sexual dimorphism in fly body size after as many as 250 generations of selection for sexual dimorphism. However, multiple studies have also provided evidence for fast, seemingly unconstrained, evolution of sexual dimorphism (Frankham, 1968a, 1968b; Bird & Schaffer, 1972; Eisen & Hanrahan, 1972; Zwaan et al., 2008; Delph et al., 2011; Kaufmann et al., 2021). For example, Bird and Schaffer (1972) selected fruit flies for sexual dimorphism on wing size and found a significant change in sex differences after only 15 generations. Such qualitative differences in outcomes are usually attributed to differences in genetic architecture underlying those traits. Specifically, that traits with a high (low) intersex correlation easily (hardly) decouple between the sexes (Stewart et al., 2010).

This prediction is supported by models of sex-specific adaptation of quantitative traits, first formulated by Lande (1980), who showed that intersex correlation determines the rate of sexually-dimorphic adaptation. Nevertheless, from the same models, it follows that as long as intersex correlation is imperfect (*r*_*fm*_ *<* 1) and given enough time, sexual conflict will be fully resolved. This suggests that, while *r*_*fm*_ poses a constraint on the *speed* of sex-specific adaptation, it is not predictive of the extent of sexual dimorphism eventually achieved. Most 2-sex models of this process (e.g. Lande, 1980; Cheverud et al., 1985) have assumed an infinitesimal genetic architecture (Lande, 1976; Barton et al., 2017), which ignores individual loci and assumes that genic (co)variances remain constant over time. However, we know that considering different genetic architectures can lead to qualitatively different results (as discussed in e.g. Rhen, 2000; Reeve & Fairbairn, 2001). For example, in single-locus (or, more generally, genetic variance-limited) models of sexual antagonism, sexual conflict is not resolved (Kidwell et al., 1977; Rice, 1984; Rhen, 2000; Morrow & Connallon, 2013), and more realistic models considering polygenic genetic architectures (Reeve & Fairbairn, 2001; Muralidhar & Coop, 2024) involve changes in genetic (co)variances over time, and thus display phenotypic dynamics that deviate from the infinitesimal predictions. In general, the relationship between sexual dimorphism and intersex correlation with a polygenic genetic architecture remains largely uncharacterized.

The second hypothesis states that a negative relationship between intersex correlation and sex differences arises because sex-specific selection favors genetic modifications that reduce the intersex covariance, which allows sex-specific adaptation (Lande, 1980, 1987; Bonduriansky & Rowe, 2005; Bonduriansky & Chenoweth, 2009; McGlothlin et al., 2019). Indeed, according to the standard picture of sexual dimorphism evolution (as discussed in e.g. Rice & Chippindale, 2001; Bonduriansky & Rowe, 2005; Cox & Calsbeek, 2009; Morrow, 2015), an initially monomorphic trait that becomes subject to sex-specific selection will decouple between sexes, allowing sex-specific means to approach their optima and resolve sexual conflict. The idea that this process involves a decrease in intersex correlation traces back to Fisher (1958, Chapter 6) and Lande (1980), who suggested that genes with sex-limited effects would accumulate over time leading to the prediction that *r*_*fm*_ will decrease as sexual dimorphism evolves. However, neither author presented a mathematical justification for this suggestion. Instead, it seems to be based on an intuition of how the intersex correlations should evolve, potentially implying the evolution of sex-specific modifiers, and generally an evolving genetic architecture (Bonduriansky & Rowe, 2005), allowing for a stable, long-term reduction in intersex correlation (Bonduriansky & Rowe, 2005; Williams & Carroll, 2009; Stewart et al., 2010). Nevertheless, the evolution of genetic architecture, in general and particularly for sexual dimorphism, is likely to be a very slow process (Williams & Carroll, 2009; Stewart et al., 2010), and is probably not occurring within the scope of shorter-term evolutionary processes, including most artificial selection experiments cited above, where phenotypes evolve without major changes in genetic architecture.

In spite of these predictions for scenarios leading to a negative correlation between intersex correlation and sexual dimorphism, the underlying mechanisms and the assumptions they require are not well understood—and addressing this gap in understanding is the main motivation of the current study.

We formulate a model of sex-specific stabilizing selection, mutation and drift (a 2-sex extension of Hayward & Sella, 2022), which is a common regime in sex-specific adaptation (Prasad et al., 2007; Abbott et al., 2010; Stulp et al., 2012; Sanjak et al., 2018), and analyze the dynamics of sexually-concordant and sexually-dimorphic evolution after a shift in sex-specific optima, while keeping track of intersex correlation over time. Given that the dynamics seem to strongly depend on the assumptions on the genetic architecture, we compare the predictions of the deterministic infinitesimal model with the evolutionary outcomes of simulations considering two types of highly polygenic architectures, which we consider non-evolving (i.e. we are not considering modifier loci leading to stable decreases in intersex covariances). The first is an approximately infinitesimal architecture, where all contributing alleles have small effect sizes and do not experience substantial changes in frequency under directional selection. The second is a less infinitesimal architecture with a significant proportion of large-effect mutations, which in humans seems to be the genetic architecture underlying most complex traits, as suggested by numerous GWAS (e.g. Wood et al., 2014; Locke et al., 2015; Simons et al., 2018).

We find that, consistent with Lande (1980)’s classical result, at equilibrium under stabilizing selection intersex correlation is independent of sexual dimorphism, when the population size is infinite and dynamics are deterministic. The classical predictions for sexual dimorphism are not entirely accurate with a finite population size, since genetic drift can generate sexual dimorphism even when selection between the two sexes is identical. We derive an expression for sexual dimorphism accounting for drift, showing that it nevertheless remains independent of intersex correlation.

By considering the transient phase of adaptation to new sex-specific optima (during which directional selection acts), we test the two hypotheses commonly used to explain the existence of a negative association between *r*_*fm*_ and sexual dimorphism. In line with the first hypothesis, that an initially low intersex correlation allows for greater evolution of dimorphism, is the previously-obtained result that intersex correlation determines the rate of sexually-dimorphic adaptation. We discuss how this might potentially lead to the expected pattern and derive an explicit expression for how sexual dimorphism changes with time depending on intersex correlation. Supporting the second hypothesis that evolution of sex differences drives a decrease in intersex correlation and agreeing with results by Reeve and Fairbairn (2001), we find that, when genetic architecture is not approximately infinitesimal, there is a transient increase in sex-specific variances leading to a temporary decrease in intersex correlation with sexually-dimorphic evolution (see Box 1 for some definitions) under directional selection. However, since the results for both hypotheses hold for divergent as well as convergent evolution (adaptation to a shift in sex-specific trait optima where they move farther apart and closer together, respectively), their contribution to the generation of a negative association between intersex correlation and sexual dimorphism requires the additional assumption that sex-specific adaptation is more commonly divergent than convergent.

Altogether, our results provide, to our knowledge, the first account of various mechanisms which can contribute to generating a negative correlation between intersex correlation and sexual dimorphism. They allow us to formalize and contextualize common intuitions in the field, as well as clearly state the assumptions and mechanisms that underlie common hypotheses, thereby providing a deeper understanding of the potential mechanisms underlying empirical observations.

Box 1. Terminology

- **Intersex correlation**: ratio between intersex covariance and geometric mean of sex-specific averages (Equation 3). It measures the correlation between the additive effects of genes as expressed in females and males.
- **Sexual dimorphism**: absolute value of the difference between female and male trait means (Equation 6). It reflects the magnitude of the difference between sex-specific averages.
- **Signed sexual dimorphism**: difference between female and male trait means (Equation 7). It reflects the magnitude and direction of sexual dimorphism.
- **Concordant adaptation**: dynamics of sex-specific trait means after a change in the average of sex-specific trait optima. Adaptation is purely concordant after a shift in optima of equal magnitude and direction between the sexes. When we refer to concordant adaptation we typically mean purely concordant.
- **Dimorphic adaptation**: dynamics of sex-specific trait means after a change in the difference between sex-specific trait optima. Adaptation is purely dimorphic after a shift in optima of equal magnitude and opposite direction between the sexes. When we refer to dimorphic adaptation we typically mean purely dimorphic. There are two types of dimorphic shifts:

**Divergent shift** brings sex-specific optima farther apart

**Convergent shift** brings sex-specific optima closer together

## 2 Methods

### 2.1 The model

We define a 2-sex extension of the standard model for the evolution of a highly polygenic, quantitative trait under stabilizing selection (S. Wright, 1935; Simons et al., 2018; Hayward & Sella, 2022). Assuming additivity, an individual’s phenotypic value follows from its genotype (Lynch & Walsh, 1998), and is given, for females (*z*_*f*_ ) and males (*z*_*m*_), by

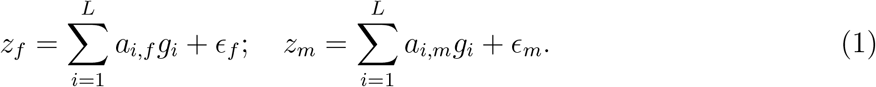

The first term is the genetic contribution, given by the sum of sex-specific phenotypic effects (*a*_*i*,*f*_ and *a*_*i*,*m*_), with *g*_*i*_ = 0, 1 or 2 indicating the number of copies of allele *i* inherited by the individual, and *L* being the target size of the trait. The second term is the sex-specific environmental contribution, which we take to be normally distributed and independent of the genetic contribution (*ϵ*_*α*_ ∼ *N* (0, *V*_*E*,*α*_) for *α* = *f, m*).

Stabilizing selection is modelled via sex-specific Gaussian fitness functions, where fitness declines with distance from sex-specific optima (*O*_*f*_, *O*_*m*_):

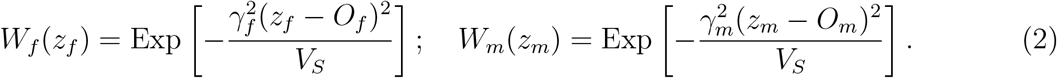

Here, 1*/V*_*S*_ determines the overall strength of stabilizing selection; *γ*_*f*_ and *γ*_*m*_ modulate the proportion of selection that acts on each sex, and satisfy 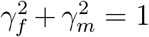. We assume that neither sex is evolving neutrally, so sex-specific selection strengths, 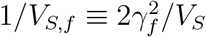 and 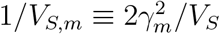, are nonzero (i.e., *γ*_*f*_, *γ*_*m*_ > 0). We choose to parametrize the problem in terms of *γ*_*f*_, *γ*_*m*_ and *V*_*S*_ instead of *V*_*S*,*f*_, *V*_*S*,*m*_ because it allows us to separate the overall strength of selection and the proportion that acts on each sex; however, replacing them with *V*_*S*,*f*_, *V*_*S*,*m*_ recovers the parametrization used in previous work (e.g. Lande, 1980). Since the sex-specific additive environmental contributions to phenotypic variation can be absorbed into *V*_*S*,*f*_, *V*_*S*,*m*_ (by replacing them with 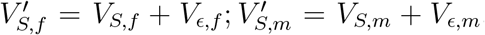, Turelli, 1984), we consider only the genetic contributions.

The population evolves according to the standard model of a diploid, panmictic population of constant size *N*, with non-overlapping generations. Exactly half of individuals are female and the other half male and, each generation, mothers and fathers are randomly chosen to reproduce with probabilities proportional to their fitness (via Wright-Fisher sampling with fertility selection). This is followed by mutation, free recombination and Mendelian segregation. We use the infinite sites approximation, which is accurate provided that the per site mutation rate, *µ*, is sufficiently low so that very few sites are hit by mutation more than once over relevant timescales (4*Nµ* ≪1). Consequently, we sample the number of new mutations per gamete per generation from a Poisson distribution with mean *U* = *Lµ*.

The sex-specific effect sizes of incoming mutations, *a*_*f*_ and *a*_*m*_, are obtained as follows: we draw the overall scaled strength of stabilizing selection on the allele (2*Ns*_*e*_) from an exponential distribution with a specific average (see Section 2.3), and we determine the fraction of stabilizing selection that acts on the allele via females (and males) from a second distribution (more details provided in Section 3.1.1.1). Sex-specific effect sizes follow from these two quantities (using Equation 17 in Section 3.1.1.1). For each mutation, we assume there is an equal probability of it being positive or negative (increasing or decreasing the trait value). In Table 1 we provide a summary of all notation used.

**Table 1:**
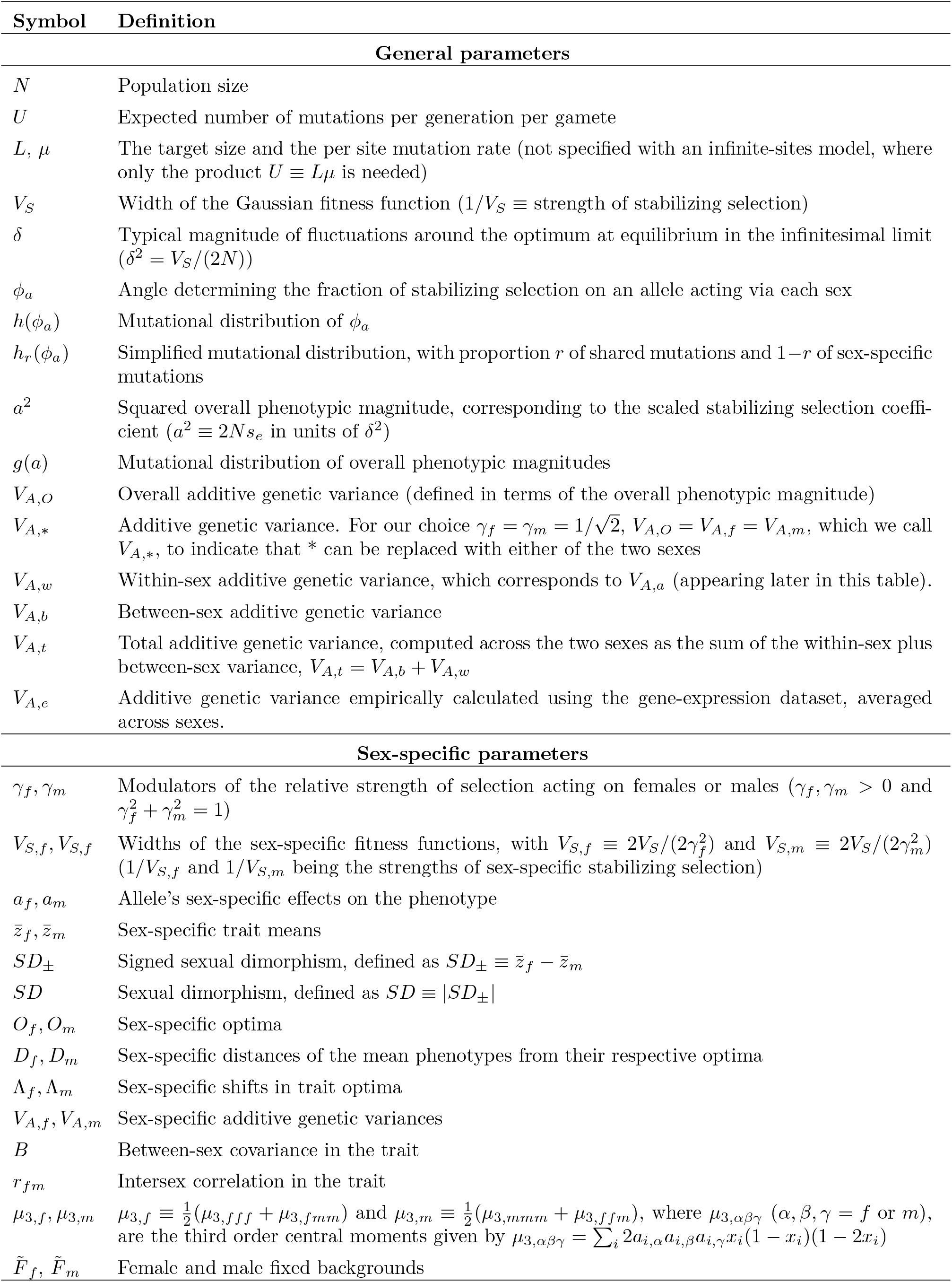
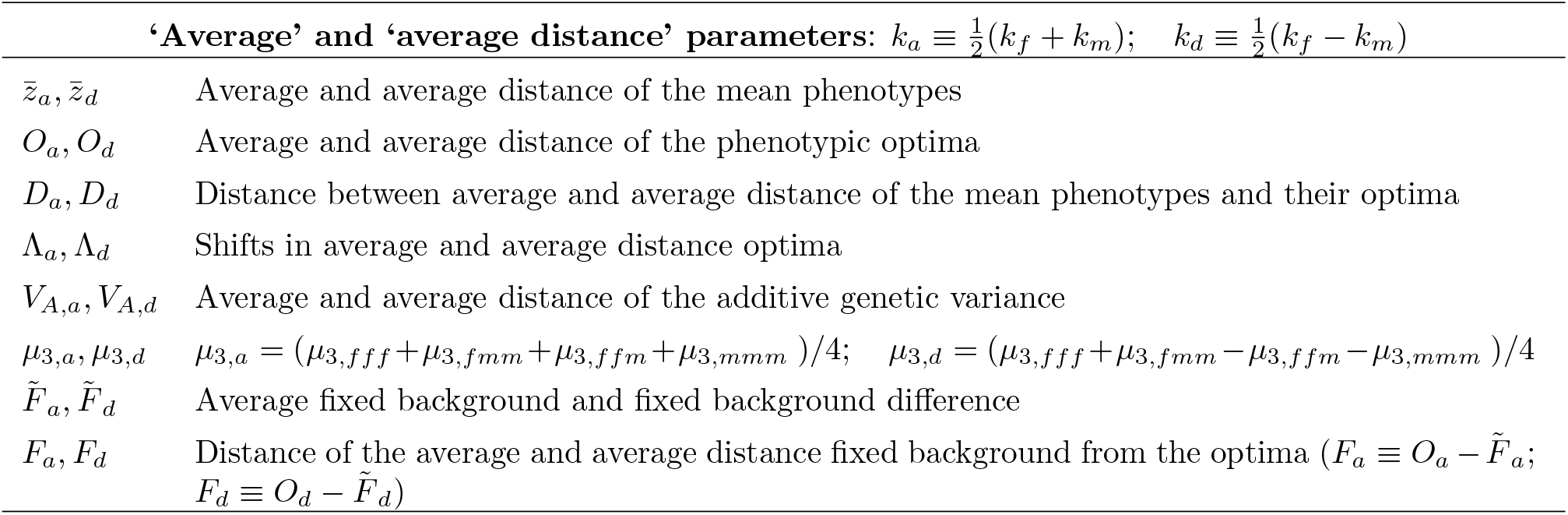
Summary of notation.

### 2.2 Parameter ranges and choice of units

We examine the genetic and phenotypic dynamics of a 2-sex population adapting to changes in sex-specific optima. We follow previous studies (Simons et al., 2018; Hayward & Sella, 2022) in defining the working parameter ranges to ensure that the conditions assumed by the analytic framework hold.

In particular, we assume that the trait is highly polygenic (2*NU* ⪢ 1) and subject to substantial but not catastrophically strong stabilizing selection. We further assume that the distance between the optimum phenotype in females (*O*_*f*_ ) and that in males (*O*_*m*_) is not massive relative to the width of the fitness function, i.e., 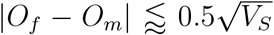 (where the symbol ⪅ denotes less than or on the same order as); see Supplementary Section 3 for details. Under these assumptions, the phenotypic distribution at stabilizing selection-mutation-drift balance is symmetric, and the sex-specific mean phenotypes exhibit small, rapid fluctuations around the respective optima, with the variance of those fluctuations given by *δ*^2^ = *V*_*S*_*/*(2*N* ) in the infinitesimal limit (Bürger & Lande, 1994). The phenotypic variance is greater than these fluctuations *V*_*A*_ *> δ*^2^, but substantially smaller than the width of the fitness function *V*_*A*_ ≪ *V*_*S*_.

After ensuring that the population is at equilibrium under mutation-selection-drift balance, we apply a shift in sex-specific optima Λ_*f*_, Λ_*m*_. We assume that the magnitude of the shift is larger than the random fluctuations of the sex-specific trait means (|Λ_*f*_ |, |Λ_*m*_| > *δ*), but smaller than, or on the order of, half the width of the fitness function 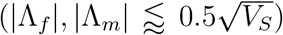. The lower bound on shift sizes was motivated by a desire to consider only non-negligible shifts, and the upper bound was motivated by the fact that our analytic predictions for (asymptotic) phenotypic variation after the shift in optimum remain accurate in the range 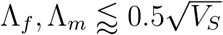 (even for tests run in the extreme case of symmetric sex-specific selection and completely shared genetic architecture between the sexes; see Supplementary Section 3).

We work in units of *δ*, the typical deviation of the population mean from the optimum at equilibrium in the infinitesimal limit. Working in these units (by setting *V*_*S*_ = 2*N* so that *δ*^2^ = *V*_*S*_*/*(2*N* ) = 1) makes our results invariant with respect to changing the population size, *N*, stabilizing selection parameter, *V*_*S*_, mutational input per generation, 2*NU*, and distributions of incoming effect magnitudes, *g*(*a*).

### 2.3 Simulations

For reasons of efficiency, our simulations are based on two additional simplifying assumptions. First, that alleles are at linkage equilibrium, allowing us to simulate the evolution of the population by tracking only the list of segregating alleles in the population, and their frequencies, rather than individuals. We refer to simulations in which we make this simplification as *Wright-Fisher* simulations because in each generation allele frequencies are updated according to a Wright-Fisher process. Second, we assume that allele frequency differences between sexes after selection are negligible (i.e., *x*_*f*_ = *x*_*m*_ = *x* so alleles are at Hardy-Weinberg equilibrium). This assumption allows us to track only average frequencies of alleles, rather than sex-specific frequencies; and we refer to simulations which make this simplification as *Hardy-Weinberg* simulations. In Supplementary Section 4, we provide more details about the assumptions behind each simulation type and test the robustness of our simulations to these two simplifying assumptions. We test the assumption of Hardy-Weinberg equilibrium by comparing the results of our *Wright-Fisher Hardy-Weinberg* simulations with *Wright-Fisher* simulations that track sex-specific allele frequencies; and we test the assumption of linkage equilibrium by comparing the results of *Wright-Fisher* simulations that track sex-specific allele frequencies with individual based simulations.

Note that the robustness of our simulation results to these tests also provides justification for the fact that our analytic framework is robust, as it relies on the same two simplifying assumptions. In addition, the assumption of Hardy-Weinberg equilibrium is plausible a priori because we consider fairly weak selection. Previous studies have shown that sexually-antagonistic selection can lead to considerable differences in allele frequencies between the sexes, where balancing selection contributes to the maintenance of substantial genetic variation (Kidwell et al., 1977; Rice, 1984; Morrow & Connallon, 2013; Connallon & Clark, 2014a). However, this requires very strong selection, beyond the range we consider in this study, and also beyond what is likely to apply to most traits.

In simulations, we let populations burn in for a period of 10*N*, 100*N* or 500*N* generations (depending on the time each parameter combination takes to reach equilibrium, stated in the respective figure captions) to ensure they attain mutation-selection-drift balance, before applying the shift in optima or taking measurements when no shift in optima applies. In figures we display averages and 95% confidence intervals (CIs) across replicates. Throughout, we simulate highly polygenic traits (2*NU* ⪢ 1) in two different parameter regimes, with genetic architectures that differ in such a way as to affect simulation results qualitatively. In the first parameter regime, simulation results are well-approximated by the infinitesimal model, which assumes that the trait is underlain by an infinite number of alleles, each with an infinitesimal effect size (Barton et al., 2017). For our modest shifts in optima, this will be the case when most mutations have fairly small effect sizes (2*Ns*_*e*_ *<* 4; corresponding to the *Lande case* in Hayward & Sella, 2022). The second parameter regime, while still highly polygenic, has a significant contribution to trait variation from larger effect alleles (with 2*Ns*_*e*_ *>* 4) and displays deviations from infinitesimal behaviour when subject to directional selection (the *Non-Lande case* in Hayward & Sella, 2022). We henceforth refer to these two types of genetic architecture as ‘approximately infinitesimal’ and ‘multigenic’, respectively.

To simulate traits with different degrees of intersex correlation, we relied on previous studies, which typically reduce the very complex regulatory genetic architecture of sex-specific trait expression into the consideration of shared and sex-specific mutations (Rhen, 2000; Reeve & Fairbairn, 2001; Bolnick & Doebeli, 2003). In this case, we assume there is a proportion *r* of shared mutations, with equal effect sizes in females and males (*a*_*f*_ = *a*_*m*_), and the remaining 1 − *r* are sex-specific, out of which half are female-specific (*a*_*m*_ = 0) and half are male-specific (*a*_*f*_ = 0). For each mutation, there is an equal probability of it increasing or decreasing the trait value. This choice of trait architecture is extremely convenient because it gives us direct control over *r*_*fm*_, as the expected intersex correlation exactly corresponds to the proportion of shared mutations (*E*[*r*_*fm*_] = *r*; see Section 3.1.1.2 for details). It is worth noting, however, that our analytic results do not rely on this simplification.

Here is a summary of the parameter values used in the simulations:

- In all simulations the population size is *N* = 1, 000 and we take 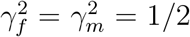, so that the strength of stabilizing selection is the same in both sexes and equal to the overall strength (*V*_*S*,*f*_ = *V*_*S*,*m*_ = *V*_*S*_).
- In all simulations (except for Figure 1) we consider an overall genetic variance of *V*_*A*_ = 40 (in units of *δ*^2^).
- In order to illustrate the approximately infinitesimal and multigenic architectures, we consider different combinations of mutation rate *U* and average squared effect size *E*(*a*^2^) (in units of *δ*^2^), sampled from an exponential distribution, yielding the same overall variance at equilibrium before the shift:
  – Approximately infinitesimal architecture: *E*(*a*^2^) = 1 (and *U* = 0.0134 for *V*_*A*_ = 40)
  – Multigenic architecture: *E*(*a*^2^) = 16 (and *U* = 0.0047 for *V*_*A*_ = 40)
- We run simulations with various *E*[*r*_*fm*_] (= *r*) values, to illustrate the evolutionary outcomes with various genetic correlations between sexes. These correspond to choices of the proportion of shared mutations of *r* = 0.5, 0.8 and 0.95 (except for Figure 1, where we cover the whole *r*_*fm*_ range).
- We typically implement shifts in sex-specific means of three concrete sizes. These correspond to 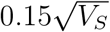 (small), 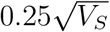 (medium) and 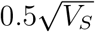 (large). These magnitudes are within the limits of the shift size for our analytical approximations to work (tested in Supplementary Section 3). Relative to the equilibrium standard deviation of the phenotypic distribution (considering *V*_*A*_ = 40), the three shift sizes correspond to: 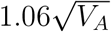 (small), 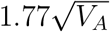 (medium) and 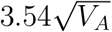 (large).
- In Figure 1, where we show simulation results for the dynamics at equilibrium, we explore a wide range of optimum differences (|*O*_*f*_ − *O*_*m*_|). The large optimum differences correspond to our three shift sizes, the small optimum differences are less than or equal to 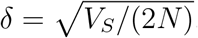, and the intermediate optimum differences are between 2 and 4 (*δ*).

**Figure 1.**
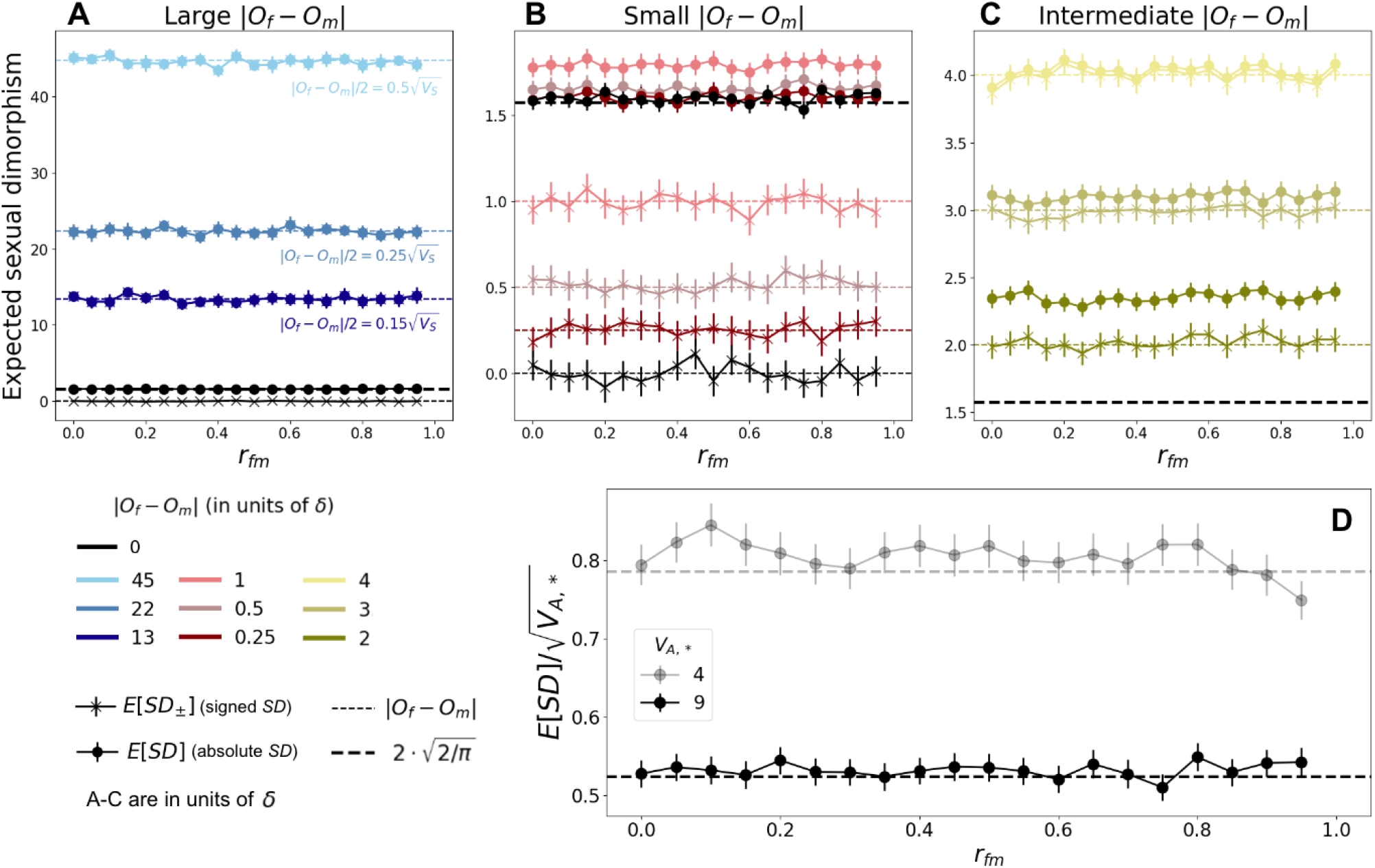
Relationship between expected intersex correlation (*r*_*fm*_) and sexual dimorphism at equilibrium with an approximately infinitesimal genetic architecture. A-C: Expected sexual dimorphism, signed (as the difference between sex-specific trait means, Equation 7; crosses) and absolute (as the absolute difference between sex-specific trait means, Equation 6; circles) across *r*_*fm*_ ∈ [0, 1), with *V*_*A*,*_ = 9 and for various |*O*_*f*_ − *O*_*m*_| ranges: large, with |*O*_*f*_ − *O*_*m*_| *>* 10 (A); small, with |*O*_*f*_ − *O*_*m*_| ∈ [0, 1] (B); intermediate, |*O*_*f*_ − *O*_*m*_| ∈ [2, 4] (C). The thick black dashed line corresponds to *E*[*SD*] predicted by Equation 33. D: Expected (absolute) sexual dimorphism, scaled by the standard deviation in sex-specific trait distributions, 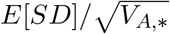, for *O*_*f*_ = *O*_*m*_ = 0 and genetic variances *V*_*A*,*_ = 4 (semi-transparent) and 9 (opaque). Simulations with *V*_*A*,*_ = 9 were run for 100*N* generations, and simulations with *V*_*A*,*_ = 4 were run for 500*N* generations. Markers and error bars indicate estimates and 95% CIs calculated as 1.96·SEM across 2,000 replicates.

Documented code for simulations can be found at

https://github.com/gemmapuixeu/Puixeu_Hayward_2025.

## 3 Results

In the present study, we examine the relationship between intersex correlation (*r*_*fm*_) and sexual dimorphism (*SD*). The intersex correlation is defined as the ratio between the between-sex covariance, *B* and the geometric mean of sex-specific variances, *V*_*A*,*f*_ and *V*_*A*,*m*_:

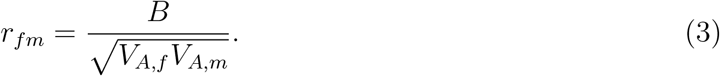

Under our assumptions of linkage equilibrium and an additive trait with no environmental contribution, *V*_*A*,*f*_ and *V*_*A*,*m*_ correspond to the sex-specific genic variances, which are the sum of the contributions to variance of all segregating alleles in each sex:

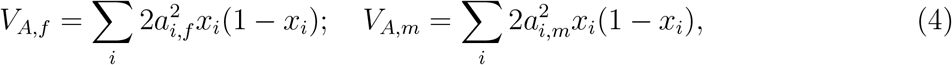

where *x*_*i*_ is the frequency and *a*_*i*,*j*_ the effect size of allele *i* in sex *j*, for *j* = *f, m*. Similarly, under our assumptions, the intersex covariance, *B*, is given by the contributions to covariance of all segregating alleles:

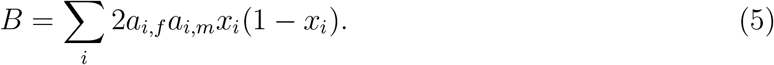

It is important to note that such calculations for *r*_*fm*_, *V*_*A*,*f*_, *V*_*A*,*m*_ and *B* are only possible in simulations where sex-specific effects and allele frequencies are known. In empirical studies, other, ‘empirical’ measures of sex-specific variances, intersex covariance and intersex correlation are needed (see Supplementary Section 5 for more details).

The definition of sexual dimorphism is less universal than that of *r*_*fm*_, as there are many ways to measure a dissimilarity between sex-specific trait means. In this study, we define sexual dimorphism to be the absolute value of the difference between sex-specific trait means:

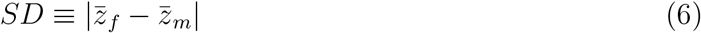

(where sex-specific trait means can be calculated by summing the allelic contributions to the mean 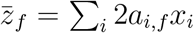 and 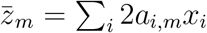). It is worth noting that some classical theoretical work (e.g. Lande, 1980; Reeve & Fairbairn, 2001) uses a signed difference in trait means to characterize sexual dimorphism:

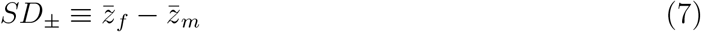

(actually, Lande (1980) and Reeve and Fairbairn (2001) consider 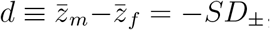, but since sexes are interchangeable in the model this sign difference has no conceptual consequences).

Nevertheless, most studies characterizing the relationship between intersex correlation and sexual dimorphism consider absolute measures. Most commonly, they consider the (sometimes error or average-normalized) absolute value of difference in trait means (McDaniel, 2005; Ashman & Majetic, 2006; Griffin et al., 2013) or absolute values of variations of the size dimorphism index (defined by Lovich & Gibbons, 1992), obtained by subtracting one from the ratio of the trait mean of the larger sex to the trait mean of the smaller sex (Bonduriansky & Rowe, 2005; Poissant et al., 2010; Leinonen et al., 2011). We choose to define *SD* as the absolute value of the difference in sex-specific averages because, of the commonly used measures, it is the simplest, and also the most similar to the signed characterization (*SD*_*±*_) used in classical theoretical work—allowing us to make comparisons in a straightforward way. In addition, in order to easily evaluate the *significance* of deviations in *SD* from zero, we sometimes scale it by the standard deviation of the phenotypic distribution.

We obtain predictions for the circumstances under which one should expect a negative correlation between *r*_*fm*_ and *SD*, a pattern which, although sometimes observed in empirical studies and often expected, we currently lack a mechanistic understanding of. To this end, we characterize the phenotypic and allele dynamics of a population at equilibrium under sex-specific stabilizing selection, mutation and drift. In Section 3.1, we describe the implications for the equilibrium relationship between intersex correlation and sexual dimorphism. Then, in Section 3.2, we examine two common hypotheses for the relationship between intersex correlation and sexual dimorphism. In order to do this, we explore the allelic and phenotypic response of a population (initially at equilibrium) to a change in sex-specific optima. We consider how these two common hypotheses are affected by assumptions made regarding 1) the genetic architecture of the trait (i.e. if the trait is approximately infinitesimal or multigenic) and 2) whether adaptation is sexually-concordant (i.e, the mean trait optimum across both sexes changes) or sexually-dimorphic (i.e, the distance between sex-specific optima changes).

Throughout our analysis we rely on the fact that (under the continuous time approximation) allele dynamics, both in and out of equilibrium, can be described in terms of the first two moments of change in allele frequency in a single generation. The first moment of change, for an allele segregating at frequency *x* with effect sizes *a*_*f*_ and *a*_*m*_ in females and males, respectively, is calculated by averaging the fitness of the three genotypes over genetic backgrounds, and is given by

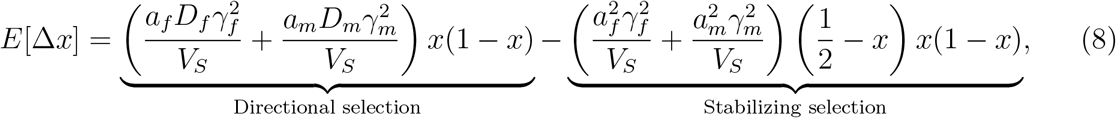

where 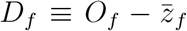 and 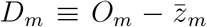 are the distances of sex-specific trait means from their respective optima (Equation 8 is derived in Section 1 of the Supplementary Material). The second moment is the standard drift term

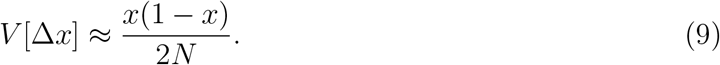

The two terms in Equation 8 reflect two selection modes. The first corresponds to directional selection, which, within each sex, acts to increase (decrease) the frequency of those alleles which move sex-specific mean phenotypes closer to (further away from) sex-specific optima; its effect becomes weaker as the sex-specific distances to the optima, *D*_*f*_, *D*_*m*_, decrease. The second term corresponds to stabilizing selection, which acts to decrease alleles’ contributions to phenotypic variance by reducing minor allele frequencies (MAFs); it weakens as the MAF approaches 1*/*2. As a reminder, 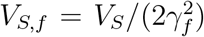 and 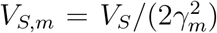correspond to sex-specific strengths of stabilizing selection. The relative importance of the two selection modes changes as *D*_*f*_, *D*_*m*_ decrease, which allows us to define two phases in the allele dynamics. An initial, rapid phase, where directional selection acts to bring sex-specific means close to the new optima via allele frequency changes, and a later, equilibration phase, in which stabilizing selection drives alleles to loss/fixation at a slower pace. More details of these processes are provided when we examine the out-of-equilibrium dynamics in Section 3.2.

### 3.1 The relationship between *r*_*fm*_ and *SD* at equilibrium

We begin by recovering, in the context of a finite population, the classical result (previously derived in the deterministic limit of an infinite population size; Lande, 1980) that, at equilibrium, expected intersex correlation and *signed* sexual dimorphism (defined in Equations 3 and 7, respectively) are independent of each other. It is important to note that, in a population of infinite size, it follows immediately from the result for signed sexual dimorphism that expected (absolute) sexual dimorphism and intersex correlation are also independent of each other. This is because population dynamics are deterministic so

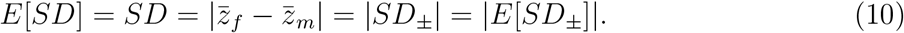

In a finite population, the relationship between absolute sexual dimorphism and signed sexual dimorphism is less straightforward; in Section 3.1.2, we explore that relationship. We show that, although genetic drift generates deviations between *E*[*SD*] and |*E*[*SD*_*±*_]| when sex specific optima are close, they are nevertheless both independent of the intersex correlation at equilibrium.

#### 3.1.1 Equilibrium *E*[*SD*_*±*_] and *r*_*fm*_ are independent

Under our assumption of an infinite sites model, and provided that at least some incoming mutations have different effects in the two sexes (i.e. *a*_*f*_ ≠ *a*_*m*_ for some alleles), directional selection will eventually drive the expected sex-specific means to their respective optima (Figure 1A-C). Thus, at equilibrium,

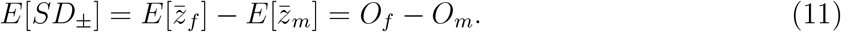

Clearly, the expression for *E*[*SD*_*±*_] does not depend on intersex correlation. To establish that expected equilibrium intersex correlation and expected signed sexual dimorphism are independent, it remains to derive an expression for expected *r*_*fm*_ at equilibrium and show that it does not depend on trait optima or trait means. In order to do this, we introduce a useful way to parameterize sex-specific allele effects.

##### 3.1.1.1 Parametrization of allele effects

At equilibrium, *D*_*f*_ = *D*_*m*_ = 0 in expectation and only the stabilizing selection term in Equation 8 is relevant:

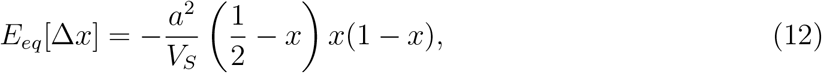

where we define *a* > 0 to be the *overall* phenotypic magnitude of an allele, with

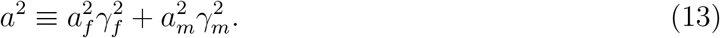

Dynamics at equilibrium for a particular allele depend only on its equilibrium scaled selection coefficient which, it follows from Equation 12 (and given our choice to measure the trait in units of *δ*, i.e. set *V*_*S*_ = 2*N* ), equals its overall phenotypic manitude:

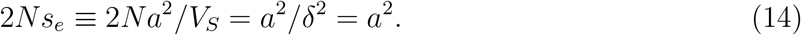

Consequently, dynamics at equilibrium are independent of mean trait values and therefore of the level of sexual dimorphism.

Although allele frequency distributions at equilibrium depend only on the overall strength of selection on alleles (captured by *a*^2^), the intersex correlation depends on whether stabilizing selection is stronger when the allele is present in a female or when it is present in a male; which we parametrize in terms of an angle, *ϕ*_*a*_. This angle directly determines the fraction of stabilizing selection on an allele that acts via females (cos^2^(*ϕ*_*a*_)) and via males (sin^2^(*ϕ*_*a*_)) and corresponds to:

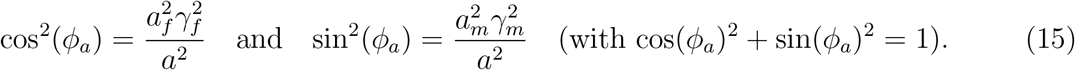

Parameterizing allele effects in terms of the allele magnitude *a*, and the angle, *ϕ*_*a*_ (rather than the sex specific effects *a*_*f*_ and *a*_*m*_), we can re-write the expected change in frequency at equilibrium under stabilizing selection (Equation 12) as

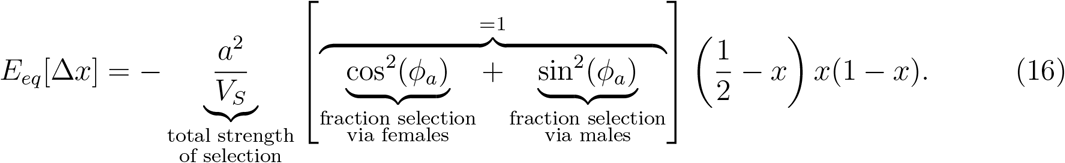

We have chosen this parameterization because the distribution of allele magnitudes, *g*(*a*), directly determines whether the genetic architecture is approximately infinitesimal or multigenic and, as we will soon demonstrate, the distribution of angles, *h*(*ϕ*_*a*_), determines the intersex correlation. However, (using *γ*_*f*_ and *γ*_*m*_) it is easy to recover the sex-specific effects from *a* and *ϕ*_*a*_:

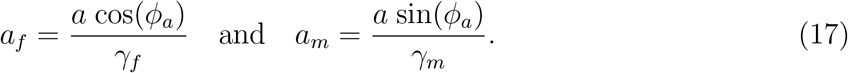

It should be noted that our analysis makes the assumption that *a* and *ϕ*_*a*_ are independent, meaning that large-effect mutations are as likely to be female- or male-biased as small-effect mutations.

##### 3.1.1.2 The intersex correlation at equilibrium

In order to characterize the inter-sex correlation we need to calculate the 2^nd^ central moments of the phenotypic distribution (*V*_*A*,*f*_, *V*_*A*,*m*_ and *B* defined in Equations 4 and 5). To do so, it is useful to define an overall genetic variance which depends on alleles’ overall phenotypic magnitudes (as defined in Equation 13):

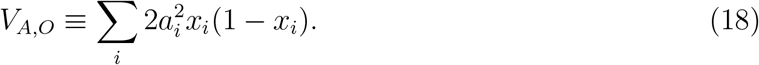

Since Equation 12 for the expected change in frequency is identical to the single-sex case for an allele with magnitude *a*, the overall variance is equal to the genic variance in the single-sex case and is given by

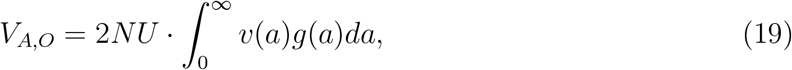

where *g*(*a*) is the distribution of incoming overall effect magnitudes and *v*(*a*) = 4*a* · *D*_+_ (*a/*2), where *D*_+_ is the Dawson function (Hayward & Sella, 2022).

In Supplementary Section 2, we show that one can compute the expressions for sex-specific variances and covariance (relative to *V*_*A*,*O*_) at equilibrium under stabilizing selection-mutation-drift balance as integrals over the distribution of angles, *h*(*ϕ*_*a*_),

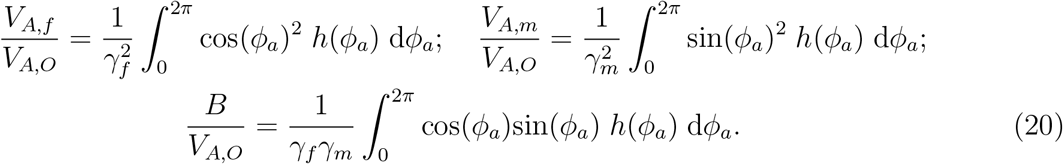

The expressions in Equation 20 can be combined to obtain the intersex correlation, yielding

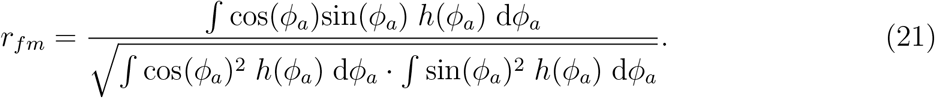

It is immediate from Equation 21, that the intersex correlation at equilibrium is independent of trait means and trait optima and therefore does not depend on the expected level of (signed) sexual dimorphism. In addition, Equation 21 shows that *r*_*fm*_ at equilibrium depends only on the fraction of stabilizing selection acting on alleles via females (or males), which is determined by the distribution of angles *h*(*ϕ*_*a*_). Since

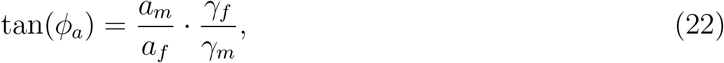

it is apparent that the parameter *ϕ*_*a*_ depends both on the ratio of alleles’ sex-specific mutational effects (i.e., *a*_*f*_ */a*_*m*_) and on the ratio of the strength of stabilizing selection in the two sexes (i.e. *γ*_*f*_ */γ*_*m*_). Thus Equation 21 demonstrates that the presence of sex-specific variation (i.e., *r*_*fm*_ *<* 1) can arise from both sex-specific mutation (*a*_*f*_ ≠ *a*_*m*_) and sex-specific stabilizing selection (*γ*_*f*_ ≠*γ*_*m*_), confirming the findings of other studies (e.g. Connallon & Clark, 2014b).

As mentioned in Section 2.3, in simulations we use a specific, highly simplified distribution *h*(*ϕ*_*a*_). In particular, we assume a proportion *r* of mutations are shared, with equal effect sizes in the two sexes (*a*_*f*_ = *a*_*m*_ and *ϕ*_*a*_ = *π/*4 or 5*π/*4), and a proportion 1 − *r* of mutations are sex-specific, out of which half are female-specific (*a*_*m*_ = 0 and *ϕ*_*a*_ = 0 or *π*) and half are male-specific (*a*_*f*_ = 0 and *ϕ*_*a*_ = *π/*2 or 3*π/*2). For each mutation, there is an equal probability of its increasing the trait (i.e, *ϕ*_*a*_ = 0, *π/*4 or *π/*2) or decreasing the trait (i.e., *ϕ*_*a*_ = *π*, 5*π/*4 or 3*π/*2). Substituting this simplified distribution of angles into Equation 21 and performing the integrals (see Supplementary Section 2.2.1) yields *E*[*r*_*fm*_] = *r*. This provides a simple way to control the expected *r*_*fm*_: we choose 0 ≤ *r* ≤ 1 and define *h*_*r*_(*ϕ*_*a*_) to be the simplified distribution described above with proportion *r* of shared mutations. Note that, although we use this simplified distribution in simulations, our analytical results are derived for general distributions *h*, provided alleles are equally likely to be positive or negative (i.e, *h*(*ϕ*_*a*_) = *h*(*ϕ*_*a*_ + *π*), e.g. Equation 21).

In simulations, in addition to using *h*_*r*_(*ϕ*_*a*_), we also typically assume that the overall strength of stabilizing selection is the same in both sexes 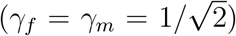 In this case, sex-specific variances are equal and in referencing them we can replace the subscripts *f* and *m* with a general *, i.e.

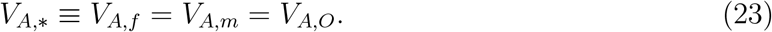

In addition, the intersex covariance is given by *B* = *rV*_*A*,*_ and the variance from shared as well as sex-specific mutations equals to:

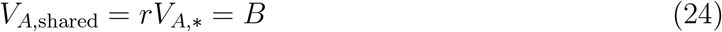

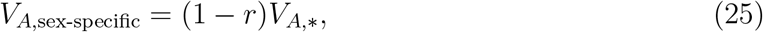

It is important to note that our expressions for *V*_*A*,*O*_, *V*_*A*,*f*_, *V*_*A*,*m*_, *B, r*_*fm*_, *V*_*A*,*_, *V*_*A*,shared_ and *V*_*A*,sex-specific_ (Equations 19, 20, 21, 23, 24 and 25) are actually expressions for the expected values of these quantities. Since, in this study, we consider only the expected values of the phenotypic variances, covariance and correlations, we suppress the *E*[…] when referring to them, for ease of reading.

It is also worth noting that neither *V*_*A*,*O*_ nor *V*_*A*,*_ capture the *total* variance in the population, as would be empirically obtained across all the individuals of both sexes. This ‘total variance’, *V*_*A*,*t*_, can be computed from allele frequency data as the sum of the the within-sex and betweensex variance:

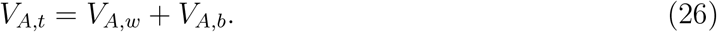

The concrete expressions for *V*_*A*,*w*_, *V*_*A*,*b*_ and *V*_*A*,*t*_ can be found in Section 6 of the Supplementary Material.

#### 3.1.2 Drift generates nonzero *E*[*SD*] even when sex-specific optima coincide

In the previous section we saw that, in expectation, between-sex correlation, *r*_*fm*_, and signed sexual dimorphism, 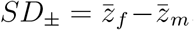, are independent of each other at equilibrium. In particular, we saw that (provided *r*_*fm*_ *<* 1) *E*[*SD*_*±*_] = *O*_*f*_ − *O*_*m*_ and that, consequently, when sex-specific optima coincide there will be no signed sexual dimorphism on average, irrespective of intersex correlation. Here, we show that, in finite populations, genetic drift can generate a nonzero average sexual dimorphism even when sex-specific optima are equal (*O*_*f*_ = *O*_*m*_). However, the amount of dimorphism generated does not depend on the intersex correlation.

The nonzero dimorphism arises from the fact that—although, in expectation, at equilibrium trait means are equal to trait optima—genetic drift leads them to undergo rapid fluctuations around their expected values (Bürger & Lande, 1994). This, in turn means that the difference in trait means, 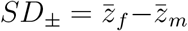, will also typically undergo fluctuations. The only exception is when the intersex correlation is 1, with all incoming mutations having identical effect in both sexes (*a*_*f*_ = *a*_*m*_). In this case, mean trait values in females and males must always coincide, and both signed sexual dimorphism and sexual dimorphism will be zero at all times (Figure S5; although *SD* displays some increase due to new mutations, which arise sex-specifically, as discussed in Supplementary Section 7.2). Indeed, whenever the intersex correlation is high, short-term fluctuations in the two trait means are highly correlated since most segregating variation has identical effects in both sexes (Figure S6C,D). However, provided *r*_*fm*_ *<* 1, mutations with effects that differ between the sexes will occasionally arise and fix, causing the two trait means to drift apart (over sufficiently long time periods). Consequently, at equilibrium sex-specific trait means will typically not be equal, *SD*_*±*_≠ 0 (Figure S4A), implying that *SD* = |*SD*_*±*_| *>* 0 and hence that *E*[*SD*] *>* 0 (Figure 1). It is easy to see that when trait values in the two sexes are uncorrelated (*r*_*fm*_ = 0), female and male trait means will fluctuate independently over both short and long time-scales (Figure S6A,B).

The fact that sexual dimorphism is nonzero for *r*_*fm*_ *<* 1 is a simple consequence of the fact that the variance in the difference in trait means is nonzero. Indeed, if the distribution of *SD*_*±*_ were Gaussian, sexual dimorphism and the variance in the difference in trait means would follow a very simple relationship:

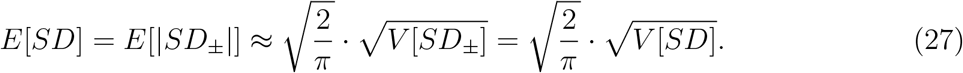

It turns out that the distribution of *SD*_*±*_ *is* well-approximated by a Gaussian distribution for both an approximately infinitesimal and a multigenic genetic architecture (QQ plot in Figure S7), and Equation 27 performs remarkably well. Consequently, we can calculate expected sexual dimorphism by calculating its variance, which is more mathematically tractable.

We begin by finding an expression for *V* [*SD*] in terms of the variance in population trait mean and the covariance in sex-specific trait means at equilibrium under stabilizing selection. From the definition of *SD* it follows that:

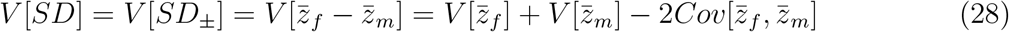

We can re-write the expression above by considering the population mean phenotype, 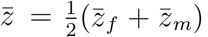 It has variance 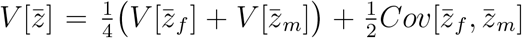, implying that 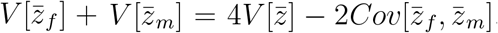. Assuming that the magnitude of fluctuations in trait mean is equal between sexes, 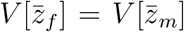, this gives us the size of sex-specific fluctuations around the optima:

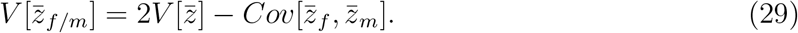

In Figure S8 we demonstrate, using simulation results for a wide range of *r*_*fm*_ *<* 1 and both an approximately infinitesimal and a multigenic genetic architecture, that 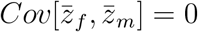 (by showing that 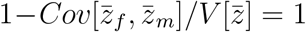; Figure S8B). Putting this result together with Equations 28 and 29 reveals that both the magnitude of sex-specific fluctuations around the optima and variance in sexual dimorphism can be expressed in terms of the variance in population mean:

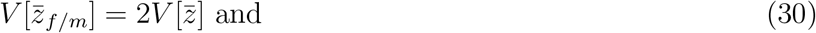

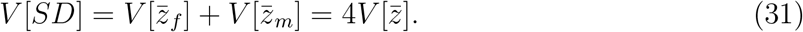

Fortunately, 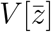 is theoretically predicted: the size of the fluctuations of the population mean around the optimum at equilibrium under stabilizing selection is 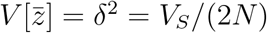 (Bürger & Lande, 1994) which, for our choice of units, equals 1 (see also Figure S8A). Consequently, the magnitude of sex-specific fluctuations around the optima is given by 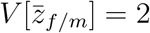 (Equation 30). Some intuition for this result can be gleaned by considering that it might arise from the fact that the population size of females and males is 0.5*N*, so that:

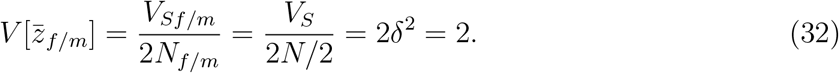

Also, following Equation 31, the expected variance in sexual dimorphism, *V* [*SD*], when *O*_*f*_ = *O*_*m*_ = 0 is equal to 4, which we recover in Figure S8C. Finally, it follows from Equations 27 and 31 that when sex-specific optima coincide, the expected sexual dimorphism is given by

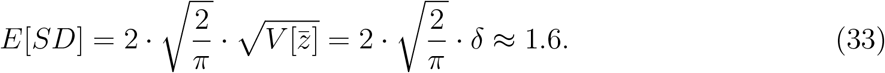

This implies that, even when selection on the two sexes is identical, the typical value of *SD* at equilibrium under stabilizing selection is nonzero and larger than than the typical deviation of the population mean phenotype from the optimum, both with an approximately infinitesimal (Figure 1B) as well as with a multigenic (Figure S3B) genetic architecture.

##### 3.1.2.1 The significance of drift-inflated

*SD* As an interesting aside to exploring the relationship between sexual dimorphism and intersex correlation, we have established that even when trait optima coincide, genetic drift is likely to induce a nonzero sexual dimorphism. However, whether or not these deviations from zero in *E*[*SD*] for *r*_*fm*_ *<* 1 are *of significance* depends on how their magnitude compares to the standard deviation in the sex-specific phenotypic distributions. Consequently, we can evaluate their significance by considering the unitless quantity 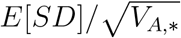 This scaling also provides a natural way to compare sexual dimorphism across different traits. Since *E*[*SD*] is on the order of *δ*, the effect of drift will be negligible for traits with genetic variance ⪢ *δ*^2^ (which is true for most traits that we simulate in this article, which have *V*_*A*,*_ = 40; see Section 2.3). However, it may well be highly relevant for traits with genetic variance on the order of *δ*^2^.

In Figure 1D we show one such example displaying 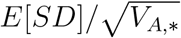 for a trait with a fairly low genetic variance of *V*_*A*,*_ = 4 or 9, which corresponds to fluctuations of the trait mean around the (shared) optimum with a typical magnitude of about half or a third of the standard deviation in the trait distribution. For these particular (low phenotypic variance) examples, the effect of drift can be highly significant. Indeed, 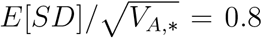 when *V* _*A*,*_ = 4 and 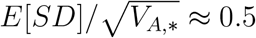 when *V*_*A*,*_ = 9, implying that, just by chance, trait means in the two sexes could frequently differ by about a full or half of a phenotypic standard deviation, respectively.

This is important because it suggests that special care should be taken before attributing even fairly large differences in female and male trait means to natural selection. The same results hold with a multigenic genetic architecture (Figure S3B).

It is worth noting that our model and most of our simulations assume linkage equilibrium which provides a good approximation for the dynamics with free recombination (see Supplementary Section 4). However, although a proper investigation into the effect of linkage disequilibrium is beyond the scope of this work, we speculate that more significant linkage disequilibrium might be expected to increase the importance of drift. This is because, in a finite population subject to stabilizing selection, linkage disequilibrium has the effect of decreasing the effective population size, and decreasing genetic variance in the trait (Santiago, 1998). In addition, the decrease in effective population size might be expected to increase the size of the random fluctuations in the sex-specific optima (since genetic drift will be stronger).

#### 3.1.3 Equilibrium *E*[*SD*] and *r*_*fm*_ are independent

When sex-specific optima coincide, the prediction of *E*[*SD*] ≈ 1.6 is supported by both theory (Equation 33) and simulations for all values of *r*_*fm*_ *<* 1 (Figure 1B). It follows immediately that sexual dimorphism and intersex correlation are independent of each other when *O*_*f*_ = *O*_*m*_. Simulation results reveal that this independence holds also for *O*_*f*_≠ *O*_*m*_. When the difference in sex-specific optima is nonzero and small (|*O*_*f*_ − *O*_*m*_| ⪅ 1), the prediction of *E*[*SD*] ≈ 1.6 for coinciding optima remains surprisingly accurate (Figure 1B). This prediction holds across genetic variances (*V*_*A*,*_) and for both approximately infinitesimal and multigenic genetic architectures (Figure S4B). For larger differences in optima (|*O*_*f*_ − *O*_*m*_| ⪆ 4), drift can be neglected and the absolute value of signed sexual dimorphism provides a good proxy for sexual dimorphism, i.e.,

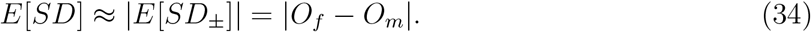

For differences in optima between these two ranges, *E*[*SD*] ≥ |*O*_*f*_ − *O*_*m*_|. Importantly, in all cases, expected sexual dimorphism and *r*_*fm*_ are independent of each other.

### 3.2 A negative relationship between *r*_*fm*_ and *SD* – exploring common hypotheses

In the previous section we describe how expected intersex correlation and sexual dimorphism at equilibrium are independent of each other at equilibrium. In this section, we examine the out-of-equilibrium dynamics of sex-specific adaptation in order to explore the two hypotheses most commonly discussed to explain the sometimes observed and often expected negative correlation between *r*_*fm*_ and *SD* (Bonduriansky & Rowe, 2005; Fairbairn, 2007; Griffin et al., 2013; Stewart & Rice, 2018; McGlothlin et al., 2019): first, that traits with ancestrally low *r*_*fm*_ are less constrained to respond to sex-specific selection and therefore evolve to be more dimorphic (H_1_: low *r*_*fm*_ precedes); second, that sex-specific selection acts to reduce the intersex correlation (H_2_: low *r*_*fm*_ follows).

We explore the applicability of these two hypotheses in the context of a population, initially at equilibrium under sex-specific stabilizing selection, mutation and drift, that is subject to a sudden environmental change leading to a shift in sex-specific optima. In our analysis, we rely on the following equation describing how the per generation change in distances between sex-specific means and their optima (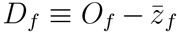 and 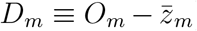) depend on the second and third order central moments of the joint female and male phenotype distribution:

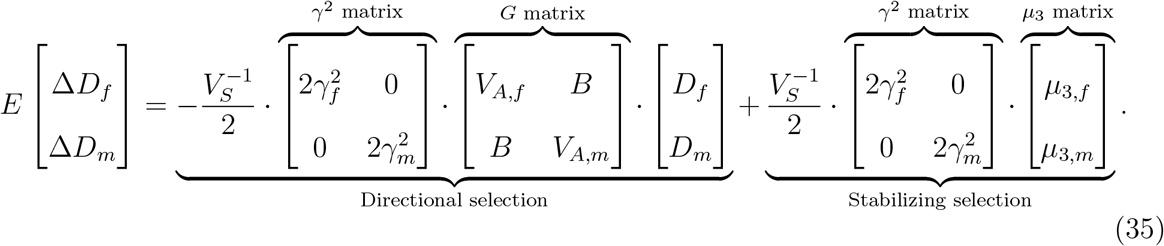

Here, 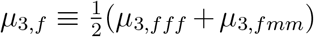 and 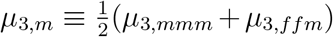, where *µ*_3,*αβγ*_ (*α, β, γ* = *f* or *m*) equal 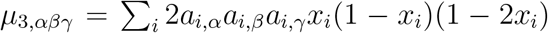and are the third order central moments of the joint female and male phenotype distribution. Equation 35 is derived by adding up the contributions to the change in mean phenotype coming from all segregating variants. Just like in the equation for alleles’ expected change in frequency (Equation 8), the two terms correspond to the two modes of selection underlying the dynamics: the first describes directional selection acting to reduce distances between means and respective optima at a rate that depends on sex-specific variances and covariance, while the second reflects the effect of stabilizing selection on an asymmetric (skewed) phenotypic distribution.

#### 3.2.1 Exploring H_1_ (low *r*_*fm*_ precedes)

This hypothesis relies on the idea that traits that initially have a low intersex correlation respond faster to novel sex-specific selection, eventually achieving higher levels of sexual dimorphism. As we saw in Section 3.1.1 and in agreement with previous results assuming a polygenic or infinitesimal genetic architecture (Lande, 1980), so long as there is variation for sexual dimorphism (in other words, if *r*_*fm*_ *<* 1), the two sexes will eventually evolve to diverge until sexual conflict is resolved – regardless of the intersex correlation (Figure 1). However, while at equilibrium (signed and absolute) sexual dimorphism is independent of *r*_*fm*_, the *rate* at which it evolves, and therefore the time frame for sexually-dimorphic evolution, is not. Consequently, in this section we characterize the time frame of adaptation to new sex-specific optima, its dependence on *r*_*fm*_ and the conditions under which this dependence can generate a negative correlation between intersex correlation and sexual dimorphism.

As in the single-sex case, this time frame can roughly be split into two phases. An initial, rapid phase dominated by directional selection (first term in Equation 35), where small changes in allele frequencies at many loci move the sex-specific means close to the new optima (which we refer to as the ‘rapid phase’); and a longer, stabilizing selection-dominated equilibration phase (second term in Equation 35), during which the small frequency differences translate into a slight increase in the fixation probability of alleles with effects that align with the shifts in optima, relative to those with effects that oppose the shifts in optima (which we refer to as the ‘equilibration phase’). We examine the impact of intersex correlation on the time frame of both phases for sexually-concordant and sexually-dimorphic adaptation of traits with approximately infinitesimal and multigenic architectures, and discuss the implications of our findings for the hypothesis H_1_ that lower intersex correlation leads to increased sexual dimorphism. We find that, in agreement with H_1_, because a high intersex correlation delays sexually-dimorphic evolution, intersex correlation might be correlated with the degree of sexual dimorphism at a given time during sex-specific adaptation. However, we also conclude that in order to show that this correlation is negative (as expected from empirical observations), an additional assumption is required.

##### 3.2.1.1 Adaptation in the infinitesimal limit: *r*_*fm*_ determines the relative rate of sexually-concordant vs sexually-dimorphic evolution

We first explore the rate of response to a change in sex-specific optima assuming an approximately infinitesimal genetic architecture. We also make the simplifying assumption that the strength of stabilizing selection is equal in the two sexes (i.e., *V*_*S*,*f*_ = *V*_*S*,*m*_ = *V*_*S*_) so that the *γ*^2^ matrix in Equation 35 is equal to the identity matrix. When the genetic architecture is approximately infinitesimal, phenotypic variances and covariance remain almost unchanged after the shift in optima, and the trait distribution remains approximately symmetric (*µ*_3,*αβγ*_ = 0 for *α, β, γ* = *f* or *m*). Consequently, Equation 35 for the expected change in the distances of the sex-specific means from the optima reduces to

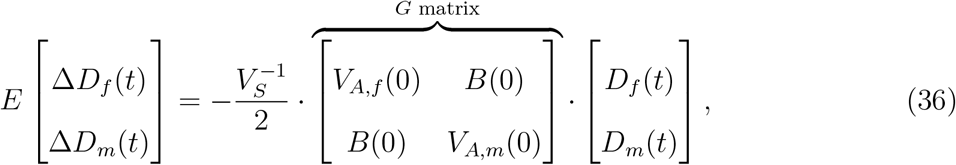

which is the 2-sex extension of the breeder’s equation, as formulated by Lande (1980). Assuming that (co)variances remain constant along time (*V*_*A*,*f*_ (0), *V*_*A*,*m*_(0), *B*(0)) this equation provides an accurate description of phenotypic evolution in the infinitesimal limit, where individual alleles do not change in frequency due to directional selection and the moments of the phenotypic distribution remain unchanged. From Equation 36, we see that after the shift in optima, directional selection acts directly on each sex to decrease the distance between the sex-specific trait mean and its optimum (*D*_*f*_ (*t*) or *D*_*m*_(*t*)) at a rate proportional to the distance itself, as well as to the initial phenotypic variance within that sex (*V*_*A*,*f*_ (0) or *V*_*A*,*m*_(0)). Directional selection within the opposite sex, however, can act to either increase or decrease the rate of adaptation to the new optimum at a rate proportional to the distance of the opposite sex from its new optimum, and to the intersex covariance, *B*(0).

To better understand the role played by intersex covariance, we follow Lande (1980) in proposing a change of variables: instead of tracking sex-specific means (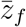 and 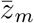), we track the ‘average’ and ‘average distance’ of their means, given by

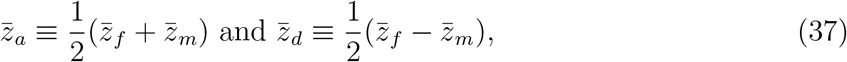

respectively. Notice that changes in 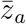 capture the evolution of the population as a whole (in fact,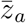 is the population mean for the trait) and changes in 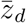 over time capture the evolution of signed sexual dimorphism, as

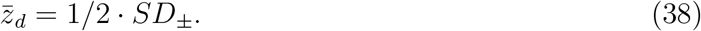

Similarly, we define an ‘average’ and ‘average distance’ version of every variable *k* that has both a female and male counterpart, as

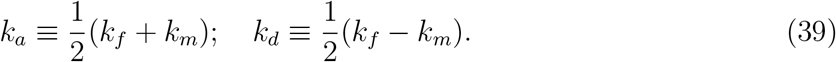

So, for example, *O*_*a*_ ≡ (*O*_*f*_ +*O*_*m*_)*/*2 and *O*_*d*_ ≡ (*O*_*f*_ −*O*_*m*_)*/*2 are the average and average distance optima. With this change of variables, we can use Equation 36 to obtain an expression for the expected per generation change in 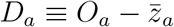 and 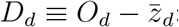:

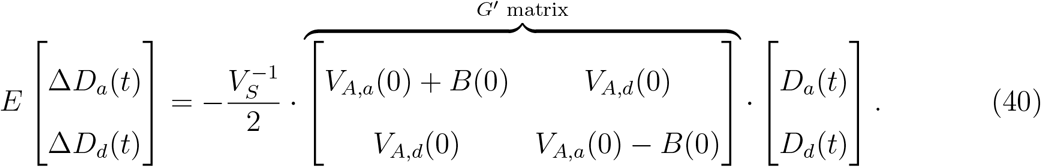

From Equation 40 it follows that: 1) a high average phenotypic variance, *V*_*A*,*a*_(0), favours the evolution of both the overall trait mean (to the new mean optimum) and sexual dimorphism (to the new difference in optima); 2) a large, positive intersex covariance, *B*(0), speeds up the evolution of the population mean to the new mean optimum, but delays the evolution of sexual dimorphism; 3) differences in phenotypic variance between the two sexes, *V*_*A*,*d*_(0) *>* 0, generate interactions in the evolution of the overall trait mean, and sexual dimorphism.

If the initial phenotypic variance is the same in the two sexes, so that *V*_*A*,*d*_(0) = 0, then the population mean and sexual dimorphism evolve independently and Equation 40 above reduces to

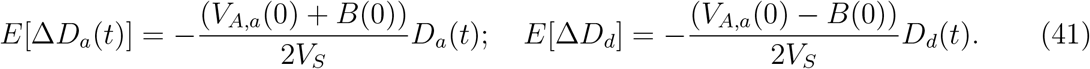

In continuous time this is solved by

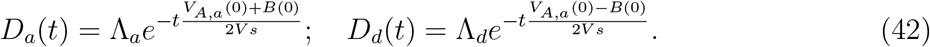

where Λ_*a*_ and Λ_*d*_ are the sizes of the shifts in *O*_*a*_ and *O*_*d*_.

Defining the length of the initial rapid phase of sexually-concordant (*t*_*a*_) and sexually-dimorphic (*t*_*d*_) adaptation to be the time that that it takes for *D*_*a*_ and *D*_*d*_ to equal the typical deviation of the population mean from the optima at equilibrium, 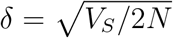 respectively, it follows that

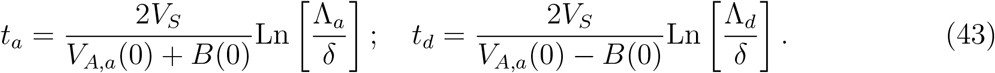

Thus the length of the initial phase of sexually-dimorphic adaptation relative to sexually-concordant adaptation is

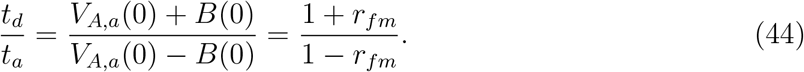

This result, initially obtained by Lande (1980), illustrates the quantitative constraint that intersex correlation places on the evolution of sex differences. In particular, when intersex correlation is close to 1, the denominator in Equation 44, 1 − *r*_*fm*_, will be very small, and sexually-dimorphic adaptation in the directional-selection dominated rapid phase could take orders of magnitude longer than sexually-concordant adaptation (*t*_*d*_ ⪢ *t*_*a*_).

These dynamics are illustrated in the top panels of Figure 2. Concretely, we implement sexually-concordant selection by applying sex-specific shifts in optima of the same magnitude and direction (Λ_*a*_ *>* 0, Λ_*d*_ = 0, shown in Figure 2A), and sexually-dimorphic selection by applying sex-specific shifts in optima of the same magnitude but in opposite directions (Λ_*a*_ = 0, Λ_*d*_ *>* 0, shown in Figure 2B), for low, intermediate and high values of intersex correlation. We see that concordant adaptation happens at a much faster rate than dimorphic adaptation, and a higher *r*_*fm*_ speeds up (slows down) concordant (dimorphic) adaptation, illustrated by a faster (slower) the reduction in *D*_*a*_ (*D*_*d*_) in Figure 2C(D). This result holds qualitatively for both the approximately infinitesimal and the multigenic genetic architectures. However, the latter shows some important quantitative differences, as we outline in the next section.

**Figure 2.**
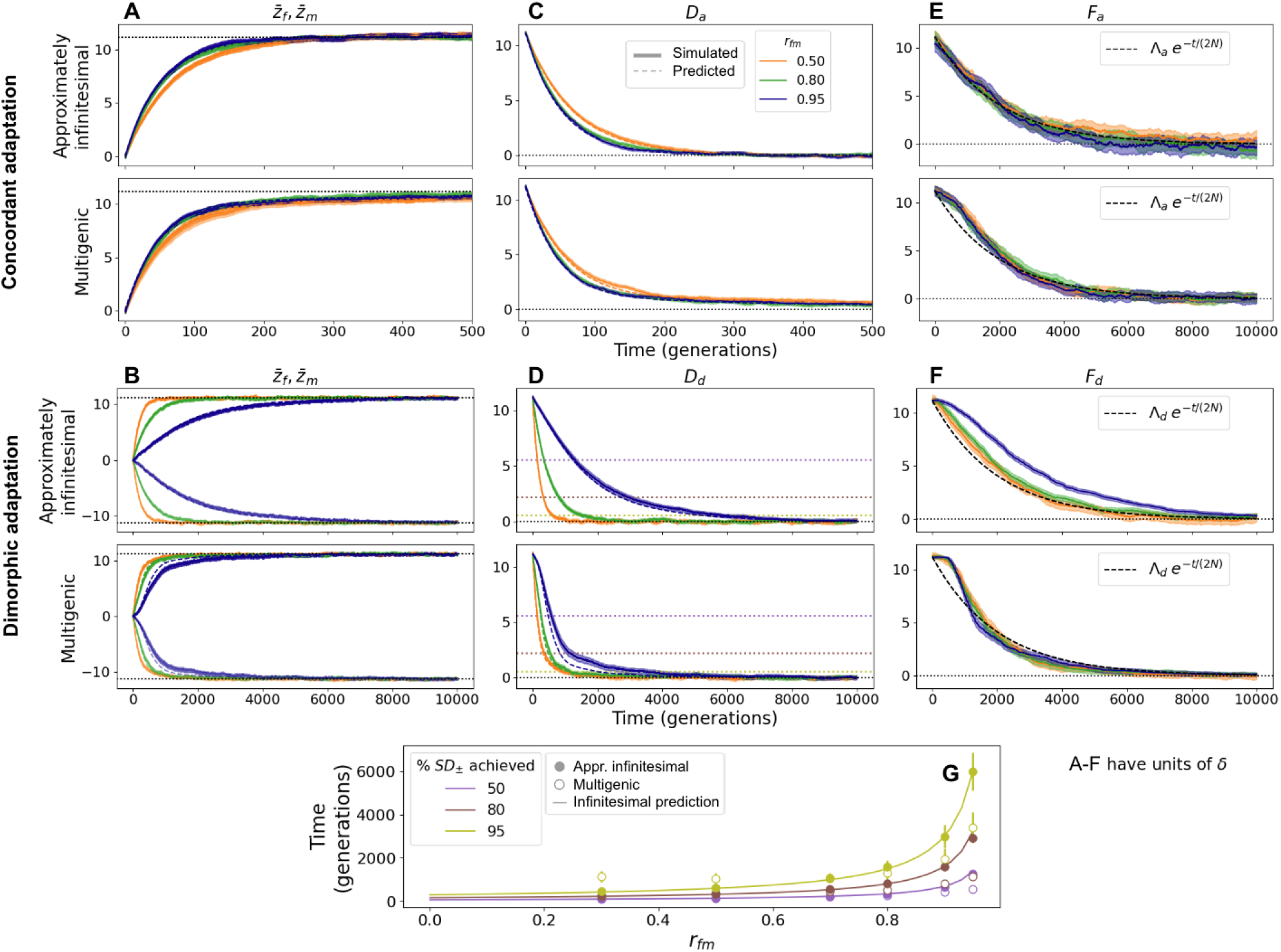
Phenotypic evolution with an approximately infinitesimal (*E*(*a*^2^) = 1, top panels) and multigenic (*E*(*a*^2^) = 16, bottom panels) genetic architecture. A: Sex-specific trait means adapting to a shift in sex-specific optima of equal magnitude and direction, which implies only sexually-concordant adaptation (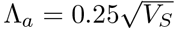 and Λ_*d*_ = 0). B: Sex-specific trait means adapting to a shift in sex-specific optima of equal magnitude and opposite direction, which implies only sexually-dimorphic adaptation (Λ_*a*_ = 0 and 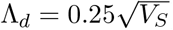). Sex-specific optima before the shift are both at zero, and after the shift are indicated as dotted lines. Thicker solid lines are simulations, and thin dashed lines are predictions using Equations 36 (approximately infinitesimal) and 46 (multigenic). C (D): *D*_*a*_ (*D*_*d*_) along time in simulations (thick solid lines) and predicted (thin dashed lines) using Equation 40 (approximately infinitesimal) and 47 (multigenic) for the sex-specific shifts in means in A (B). E (F): *F*_*a*_ (*F*_*d*_) along time for the optima shifts in A (B). Coloured lines correspond to simulations and the dashed black line corresponds to the prediction according to Equation 49 (50). G: Time to reach a given percentage of *SD*_*±*_ (50, 80 and 90%, as purple, dark red and olive circles; indicated as dotted horizontal lines in D) in simulations with approximately infinitesimal (solid) and multigenic (empty) genetic architectures and various levels of *r*_*fm*_. Lines correspond to the infinitesimal prediction using Equation 43. All simulations have been run for 10*N* generations before the shift in optima, and for three levels of *r*_*fm*_: 0.5 (orange), 0.8 (green) and 0.95 (blue; only G has more *r*_*fm*_ data points). Results display averages and 95% CIs computed as 1.96·SEM across 200 replicates. The *x*-axis in A and C spans a far shorter time period reflecting the fact that the initial phase of concordant adaptation tends to occur far more rapidly than dimorphic adaptation. All quantities displayed in A-F are in units of *δ*.

Considering the simple relationship between 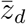 and *SD*_*±*_ (Equation 38), we obtain an expression for the signed sexual dimorphism over time:

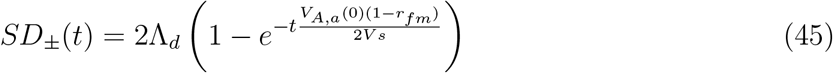

Equation 45 shows that the amount of sexual dimorphism at a given time after a shift in sex-specific optima, depends on the shift in the difference between sex-specific optima, the strength of selection, the average genetic variance of the trait considered and the intersex correlation.

When differences in trait means are large or moderate 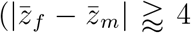 Section 3.1.2) we expect sexual dimorphism and signed sexual dimorphism to be similar; and in order to explore out-of-equilibrium dynamics we follow previous theoretical work (Lande, 1980; Reeve & Fairbairn, 2001) and frequently consider 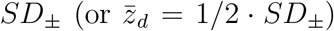 as proxies for *SD*, especially when deriving analytic expressions. This removes the need to deal with (likely complicated) deviations between *SD*_*±*_ and *SD* introduced by genetic drift: deviations that are unlikely to be illuminating for our purpose of exploring the common hypotheses. Importantly, in figures explicitly related to testing the common hypotheses (3C, D and 4F) we show results for the absolute *SD* and these confirm intuitions gleaned from considering signed *SD*.

##### 3.2.1.2 Adaptation with a multigenic genetic architecture: transient changes in the 2^nd^ and 3^rd^ order moments of the phenotype distribution alter the dynamics of phenotypic adaptation

The accuracy of the predictions for the evolution of phenotypic means in Equations 36 and 40 relies on the assumption that the respective *G* and *G*^*′*^ matrices remain constant over time. This will be approximately true when the genetic architecture is approximately infinitesimal. However, when considering a less infinitesimal trait architecture, with a significant proportion of mutations with larger effect sizes (*a*^2^ *>* 4) as exemplified by our multigenic trait architecture, the approximations in Equations 36 and 40 are no longer accurate. This is because directional selection on alleles with larger effects can generate a significant increase in the 2^nd^ central moments of the joint phenotype distribution, as well as the establishment of nonzero third central moments (Figure S9). To accurately predict phenotypic evolution with a multigenic genetic architecture we therefore need the full expression for the expected change in the distances of the sex-specific means from their respective optima (Equation 35), with time-dependent 2^nd^ and 3^rd^ central moments (i.e. *V*_*A*,*f*_ (*t*), *V*_*A*,*m*_(*t*), *B*(*t*), *µ*_3,*f*_ (*t*), *µ*_3,*m*_(*t*)). Assuming, as we did for the approximately infinitesimal architecture, that the strength of stabilizing selection is equal in the two sexes (i.e., *V*_*S*,*f*_ = *V*_*S*,*m*_ = *V*_*S*_) Equation 35 simplifies to:

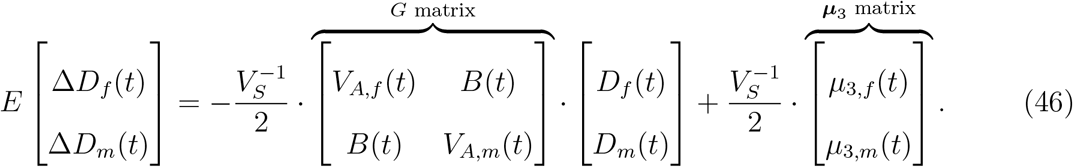

As before, a simple change of variables (Equation 39) yields an expression for the evolution of the overall trait mean (captured by *D*_*a*_) and the level of sexual dimorphism (captured by *D*_*d*_):

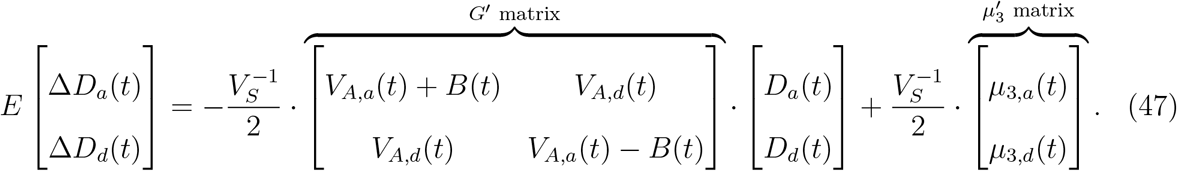

By updating (co)variances and 3^rd^ central moments according to simulation results, we can use Equation 47 to accurately predict the mean trajectories of *D*_*a*_ and *D*_*d*_.

In cases where the trait has a multigenic genetic architecture, changes in the 2^nd^ and 3^rd^ central moments of the phenotypic distribution *do* affect the trajectories of mean phenotypes. However, they affect *D*_*a*_ and *D*_*d*_ in qualitatively different ways (see Figure S9 for changes in 2^nd^ and 3^rd^ central moments in both concordant and dimorphic adaptation, and Figure S10 for their separate effects on phenotypic evolution).

For concordant adaptation after a change in sex-specific means of equal magnitude and direction (captured by the decay of *D*_*a*_), the dynamics are highly analogous to those observed in the single sex case (Hayward & Sella, 2022). Adaptation during the rapid phase occurs similarly as in the infinitesimal case and is well approximated by Equations 42, 43 and 44 (Figure 2A,C; Figure S10, top). However, after the rapid phase, the average trajectories of *D*_*a*_ can deviate significantly from the exponential decrease predicted by Equation 42. Once the mean phenotype nears the new optimum, the system enters the equilibration phase when the decreasing distance and increasing 3^rd^ central moments reach the point at which the two terms on the right-hand side of Equation 47 approximately cancel out, and the changes in *D*_*a*_ come almost to a stop (Figure 2C bottom, S10 top). The rates of approaching the new optima are then largely determined by the rate at which the 3^rd^ central moments decay. This roughly corresponds to the rate at which the allele frequency distribution equilibrates (changes in frequency generated by directional selection translate into fixed differences, as we discuss in Section 3.2.1.3). At this point, the system attains the original mutation-selection-drift balance around the new optima, and 2^nd^ and 3^rd^ central moments of the phenotypic distribution are restored to their equilibrium values. Indeed, the phenotypic dynamics for a trait with multigenic genetic architecture at the beginning of the equilibration phase are well captured by a quasistatic approximation (derived in Supplementary Section 8.1 and illustrated in Figure S11). We find that, while intersex correlation determines the time it takes to reach the equilibration phase (given approximately by Equation 43), it does not seem to make a qualitative difference in the trajectories of the means during the initial part of the equilibration phase. However, as we demonstrate in the next section, a higher intersex correlation does imply a longer equilibration phase.

In contrast to concordant adaptation, dimorphic adaptation after a shift in sex-specific optima of equal magnitude and opposite directions (captured by decay in *D*_*d*_), shows qualitatively different dynamics with a multigenic genetic architecture: concretely, changes in 2^nd^ and 3^rd^ central moments of the phenotypic distribution appear to significantly speed up phenotypic adaptation with respect to the infinitesimal predictions (Figure 2B,D,G; Figure S10, bottom). Indeed, we find that the time it takes for *SD* to evolve is much less in the multigenic case than in the infinitesimal case, especially for higher levels of *r*_*fm*_ (Figure 2D,G). For example, as Figure 2G illustrates graphically, to achieve an 80% of the total difference in *SD* after a shift in optima (in dark red), it takes 50% and 220% longer with *r*_*fm*_ of 0.8 and 0.95, respectively, with an infinitesimal genetic architecture relative to a trait with a multigenic architecture. These results suggest that the quantitative constraint that *r*_*fm*_ poses on dimorphic adaptation derived for the infinitesimal case (Equation 44) is not as strict for traits with a less infinitesimal genetic architecture; an observation of potential empirical significance.

##### 3.2.1.3 Higher intersex correlation delays equilibration for sex differences

In Section 3.2.1.1 we described how the time required for the average and average distance of the sex-specific trait means to approach their new optima depends on *r*_*fm*_ (Equations 43 and 44). These timepoints correspond to the length of the inital, directional selection-dominated phases of sexually-concordant and sexually-dimorphic adaptation, which are driven by small changes in allele frequencies at many loci. In this section, we analyze the timeframe associated with equilibration, during which stabilizing selection translates the allele frequency differences (generated by directional selection) between alleles with phenotypic effects that are aligned and opposed to the phenotypic shift into differences in fixation probabilities. This process restores the equilibrium phenotypic distributions with means at the new optima.

To examine the dynamics of equilibration we track the female and male fixed backgrounds (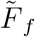 and 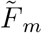), defined as the trait values that females or males in the population would have if every segregating derived allele went extinct; and can be thought of as the component of the mean phenotypes maintained by fixations (as opposed to segregating variation). As before, we distinguish between sexually-concordant and sexually-dimorphic adaptation by performing a change of variables. Using Equation 39, we define the average fixed background and the fixed background difference 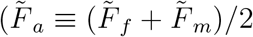 and 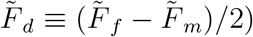 Their distances from the new optima are:

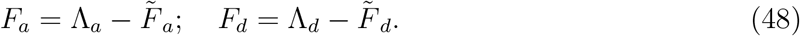

At equilibrium, we expect the fixed distances, *F*_*a*_ and *F*_*d*_, to be 0; the rate at which *F*_*a*_ (*F*_*d*_) approaches 0 gives the timescale over which sexually-concordant (sexually-dimorphic) equilibration occurs.

Not unexpectedly, we find that, in the approximately infinitesimal regime, sexually-concordant equilibration takes place at much the same rate as when there is just a single sex, and thus the trajectory of *F*_*a*_ is well-approximated by

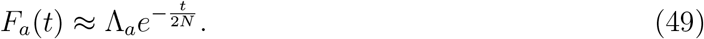

(Hayward & Sella, 2022; Figure 2E). Sexually-concordant equilibration thus occurs over a time period on the order of 2*N* generations. Somewhat surprisingly, we find that when the intersex correlation is fairly low, *F*_*d*_ also decays approximately exponentially at a rate 1*/*(2*N* ):

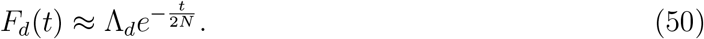

(Figure 2F). When intersex correlation is high, however, the approximation in Equation 50 becomes quite inaccurate since the decay of *F*_*d*_ can be quite delayed (Figure 2F). Thus high intersex correlation increases the time period over which sexually-dimorphic equilibration occurs in the approximately infinitesimal case.

With a multigenic trait architecture, however, we observe slight deviations from exponential decay in *F*_*a*_ and *F*_*d*_ (even when intersex correlation is low). This is analogous to the deviations observed with a multigenic architecture in the single-sex case (Hayward & Sella, 2022). In particular, the decay is initially slower and later faster then predicted by the approximations in Equations 49 and 50 (Figure 2E,F bottom). However, the time taken for the fixed backgrounds to reach the new optima, and therefore for the various moments of the phenotypic distribution to be restored to equilibrium values, is nevertheless on the order of 2*N* generations.

##### 3.2.1.4 H_1_ holds – given an additional assumption

We have shown that, while intersex correlation does not predict the overall realized sexual dimorphism, it does determine the rate at which it evolves. First, it directly determines the rate of sexually-concordant vs dimorphic phenotypic adaptation in the rapid phase; second, a high intersex correlation can delay sexually-dimorphic equilibration. When considering non-equilibrium dynamics of adaptation, these aspects might contribute to generate an overall, negative relationship between *r*_*fm*_ and sexual dimorphism, consistent with the first common hypothesis that initially lower intersex correlation allows for faster decoupling between sexes and more sexual dimorphism evolution. However, this only holds given the extra assumption that traits are more likely to be selected to diverge (i.e., trait optima move further apart) than to converge (i.e., trait optima move closer together) between sexes.

This extra assumption is required because a lower intersex correlation allows for a faster sexually-dimorphic evolution, both after a divergent as well as *convergent* shift in sex-specific optima (Figure 3A,B). Concretely, after a divergent shift in sex-specific optima (i.e. keeping *O*_*a*_ constant and increasing the absolute value of *O*_*d*_), traits with a higher intersex correlation will take longer to diverge between sexes, leading to the commonly expected pattern of a negative relationship between intersex correlation and sex differences at a given time during divergent adaptation (black dashed line in Figure 2C,D, corresponding to the timepoint of the black vertical dashed line in Figure 2A,B). However, this is also true for adaptation after a convergent shift in optima (i.e. keeping *O*_*a*_ constant and decreasing the absolute value of *O*_*d*_): traits with a higher intersex correlation will take longer to adapt to a convergent shift than traits with an initially lower *r*_*fm*_, potentially leading to the opposite pattern, i.e. to a *positive* relationship between intersex correlation and sex differences at a given time during convergent adaptation (grey solid line in Figure 2C,D, corresponding to the timepoint of the grey vertical solid line in Figure 2A,B). Importantly, because *r*_*fm*_ imposes a stronger constraint on dimorphic adaptation in the infinitesimal case, this effect is markedly weaker when considering multigenic genetic architectures (Figure 2B,D).

**Figure 3.**
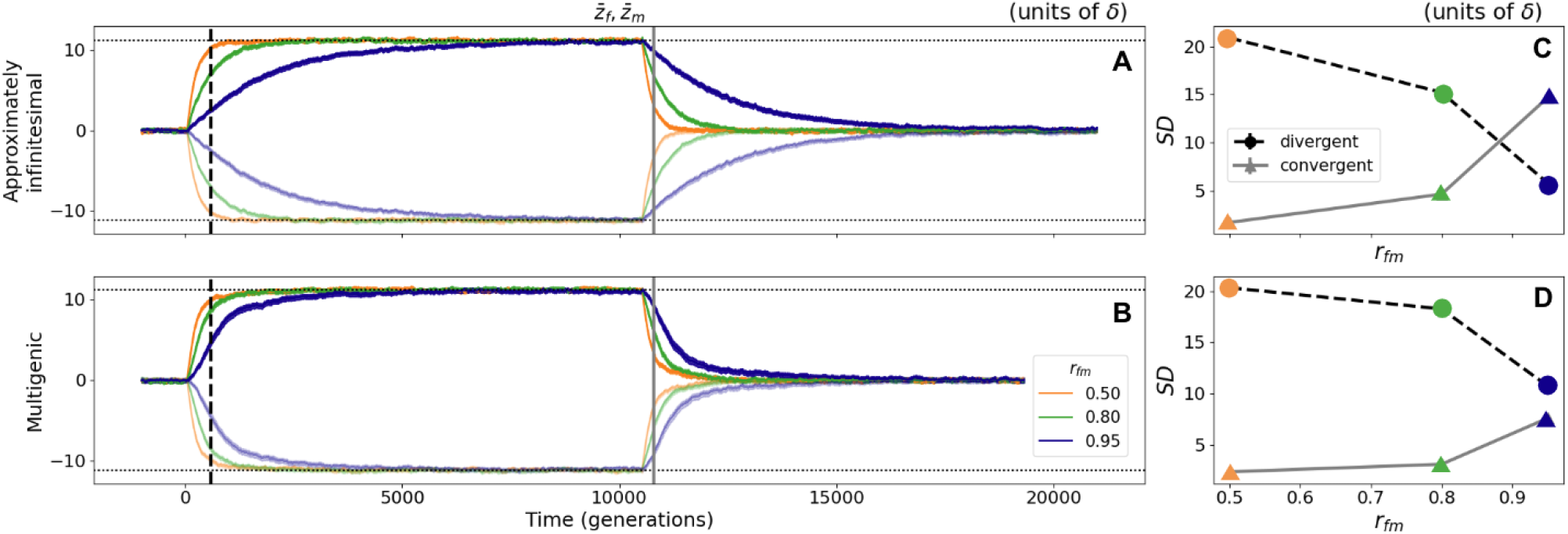
Negative (positive) correlation between *r*_*fm*_ and *SD* with divergent (convergent) adaptation. A (B): Sex-specific trait means adapting first to divergent and then to convergent shifts in optima of magnitude 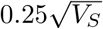 for approximately infinitesimal (multigenic) genetic architectures and three levels of *r*_*fm*_: 0.5 (orange), 0.8 (green) and 0.95 (blue). C (D): Sexual dimporhism (given by the absolute value of the difference between sex-specific means, Equation 6) for the three different levels of *r*_*fm*_ at a given point of sexually-dimorphic divergent—black dashed, corresponding to the timepoint of the black dashed vertical line in A (B)—and convergent —grey solid, corresponding to the timepoint of the grey solid vertical line in A (B)—adaptation, with an approximately infinitesimal (multigenic) genetic architecture. Results display averages and 95% CIs across 200 replicates.

#### 3.2.2 Exploring H_2_ (low *r*_*fm*_ follows): sex-specific directional selection *transiently* reduces *r*_*fm*_

In this section we explore the hypothesis, often stated as an alternative to H_1_ (discussed in Section 3.2.1), that a negative correlation between *r*_*fm*_ and sexual dimorphism arises as a consequence of sex-specific adaptation driving a reduction in intersex correlation. To do so, we examine how intersex correlation evolves with sexually-dimorphic adaptation. Intersex correlation depends both on the variances within a single sex, *V*_*A*,*f*_ and *V*_*A*,*m*_, and on the covariance, *B* (Equation 3). In Section 3.2.1.1, we established that for traits with approximately infinitesimal genetic architectures, the 2^nd^ order central moments remain approximately unchanged by directional selection (Figure 4A,C,D; see Supplementary Section 8 and Figure S9 for a more detailed discussion on the evolution of these moments). Consequently, when the trait has an approximately infinitesimal architecture, intersex correlation does not evolve at all (Figure 4A,E). In contrast, as we discussed in Section 3.2.1.2, for traits with multigenic architectures directional selection generates transient changes in 2^nd^ central moments of the phenotypic distributions (Figure S9 and Figure 4B,C,D). These changes *can* result in a temporary decrease in intersex correlation (Figure 4B,E).

**Figure 4.**
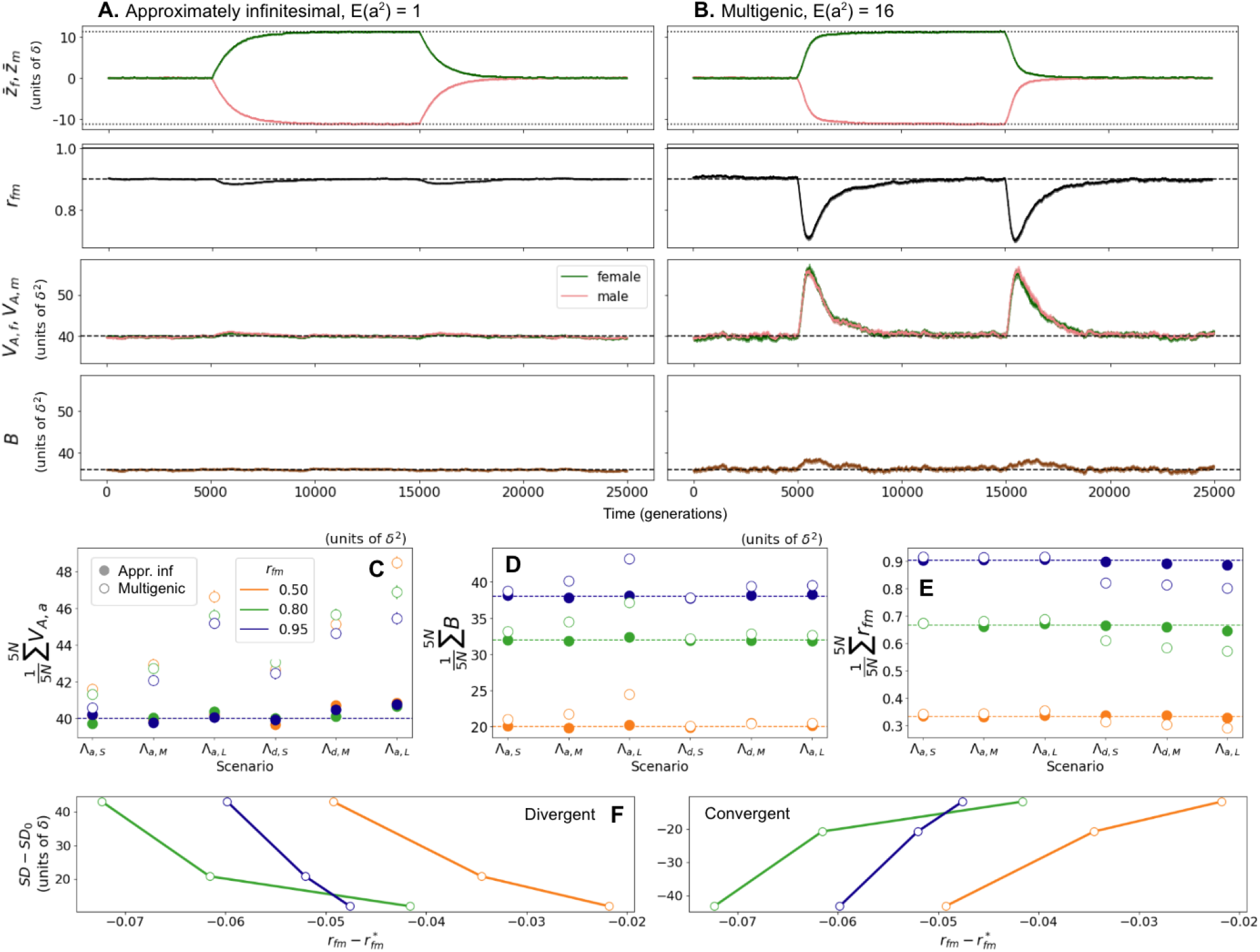
Transient decrease in *r*_*fm*_ during sexually-dimorphic (divergent and convergent) evolution. A, B: Evolution of sex-specific trait means 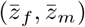, intersex correlation (*r*_*fm*_), sex-specific variances (*V*_*A*,*f*_, *V*_*A*,*m*_) and covariance (*B*) along time with an approximately infinitesimal (A, *E*(*a*^2^) = 1) and multigenic (B, *E*(*a*^2^) = 16) genetic architecture. We let the population evolve for 10*N* generations before and after applying a shift in sex-specific optima of magnitude 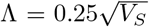 inducing divergent (optima move apart), and then convergent (optima move together) evolution between the sexes. C, D, E: Means of average genetic variance (*V*_*A*,*a*_; C), covariance (*B*; D) and intersex correlations (*r*_*fm*_; E) across 5*N* generations after the shift in optima ( empirical integrals during the rapid phase of adaptation), for approximately infinitesimal (solid circles) and multigenic (open circles) genetic architecture and across different scenarios indicating different types of shifts: Λ_*a*,_ are shifts of same magnitude and direction in both sexes, leading to sexually-concordant adaptation (similar to scenario depicted in Figure 2A, in which Λ_*d*,_ = 0); Λ_*d*,_ are shifts of same magnitude and opposite direction in both sexes, leading to sexually-dimorphic adaptation (similar to scenario in Figure 2B, in which Λ_*a*,-_0). Λ_-,*S*_, Λ _-,*M*_ and Λ _-,*L*_ indicate small, medium and large shifts, with magnitudes 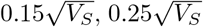 and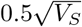, respectively. F: Negative (positive) relationship between intersex correlation and sexual dimorphism with divergent—left— (convergent, right) sexually-dimorphic selection. The *y*-axis corresponds to the difference between (theoretical predictions of) sexual dimorphism before and (long) after the shift, for the three shift magnitudes (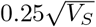 and 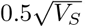); on the *x*-axis is the difference between the average *r*_*fm*_ across 5*N* generations after the shift with a multigenic genetic architecture (corresponding to the open circles in Λ_*d*,*S*_, Λ_*d*,*M*_ and Λ_*d*,*L*_ in E), and the equilibrium *r*_*fm*_ values (dashed horizontal lines in E), for the three *r*_*fm*_ (0.5, 0.8 and 0.95 in orange, green and blue). A-E display averages and 95% CIs across 200 replicates.

This decrease in intersex correlation (for traits with a multigenic architecture) is expected for sexually-dimorphic adaptation (i.e., when the distance between sex-specific trait optima changes), but not for sexually-concordant adaptation (i.e., when the mean optimum trait value changes for the two sexes equally). With sexually-concordant adaptation there is selection for phenotypic change along the main diagonal of the *G* matrix (under our assumption that *V*_*A*,*f*_ = *V*_*A*,*m*_), so there is an increase in sex-specific variance contributed by *shared* (but not sex-specific) mutations, which is equal to the increase in between-sex covariance (Figure S9B,C,E; Figure S12). Consequently, intersex correlation, which is a ratio of the two, remains constant over time regardless of the magnitude of the shift (scenarios Λ_*a*,*S*_, Λ_*a*,*M*_ and Λ_*a*,*L*_ in Figure 4C,D,E).

However, with sexually-dimorphic adaptation, directional selection drives an increase in frequency of those sex-specific mutations which drive phenotypic change in the direction of the shift, leading to an increase in sex-specific variances. Nevertheless, it does not on average increase the frequency of shared mutations, so covariance remains at equilibrium values (Figure S9H,I,K; Figure S12), which leads to a decrease in *r*_*fm*_ (scenarios Λ_*d*,*S*_, Λ_*d*,*M*_ and Λ_*d*,*L*_ in Figure 4C,D,E). This is only a transient phenomenon; as described in Section 3.2.1.3, (co)variances, as well as *r*_*fm*_ are restored to their equilibrium values during the equilibration phase, over a time periond on the order 2*N* (Figure 2E,F).

The potential transient decrease in sexual dimorphism described above could generate an association between intersex correlation and sexual dimorphism. However, the direction of this association depends on whether sexually-dimorphic adaptation is divergent (i.e., sex-specific optima move further apart) or convergent (i.e., sex-specific optima move closer together). For some intuition, let us consider a set of monomorphic (dimorphic) traits with similar *r*_*fm*_ values at equilibrium, a subset of which becomes sex-specifically selected after a divergent (convergent) shift in sex-specific optima. Those traits in the process of diverging (converging) will experience a temporary decrease in intersex correlation, which would generate a negative (positive) correlation between *r*_*fm*_ and sexual dimorphism. The negative (positive) association between intersex correlation and sexual dimorphism that might arise as a consequence of divergent (convergent) sexually-dimorphic adaptation is illustrated in Figure 4F.

These results indicate that, in accordance with H_2_, a negative correlation between intersex correlation and sexual dimorphism could arise from sex-specific adaptation leading to a reduction in *r*_*fm*_. However, this phenomenon is only transient. In addition, it only applies when some additional conditions are met. First, at least some traits must have a non-infinitesimal genetic architecture, where (co)variances change under directional selection; second, traits must be adapting to (partially) non-concordant directional selection between sexes, where (a subset of) sex-specific mutations are more beneficial than shared mutations; third, this sexually-dimorphic adaptation must be divergent more frequently than it is convergent.

## 4 Discussion

Based on the quantitative constraint that a high intersex correlation poses on the evolution of sexual dimorphism (Lande, 1980, 1987; Stewart & Rice, 2018) is the general idea that they should negatively correlate with one another, either because traits will evolve to be more dimorphic if they are less correlated between the sexes (which we discuss as hypothesis H_1_: ‘low *r*_*fm*_ precedes’; Bolnick & Doebeli, 2003; Poissant et al., 2010; Stewart & Rice, 2018) or because sexually dimorphic evolution leads to a decrease in intersex correlation, which should allow independent adaptation of both sexes (which we discuss as hypothesis H_2_: ‘low *r*_*fm*_ follows’; Lande, 1980; Bonduriansky & Rowe, 2005; Bonduriansky & Chenoweth, 2009; McGlothlin et al., 2019). Although these are common assumptions in the sexual dimorphism literature, partially based on the empirical observation of a general negative correlation between *r*_*fm*_ and sexual dimorphism (Ashman, 2003; Delph et al., 2004; Bonduriansky & Rowe, 2005; McDaniel, 2005; Poissant et al., 2010; Griffin et al., 2013), we lack a mechanistic understanding for them, which poses the main motivation of the present study: we use models of sex-specific stabilizing selection, mutation and drift to explore the relationship between intersex correlation and sexual dimorphism, and outline the conditions under which a negative correlation between the two is expected.

First, we reproduce the well-known result (first obtained by Lande, 1980) that, for a highly polygenic or quantitative trait with enough sex-specific genetic variation (either because there is enough standing variation or we have substantial sex-specific mutational input) sexual conflict will be resolved, in the sense that, given enough time and as long as *r*_*fm*_ *<* 1, sex-specific means will eventually align with their optima (Figure 1A). We derive explicit expressions to illustrate that allele dynamics at equilibrium under stabilizing selection are independent of trait optima and, consequently, trait means (Equation 12); instead, they depend on the overall strength of stabilizing selection (Equations 14). We show that the *G* matrix at equilibrium depends only on the overall and sex-specific mutational input and selection strength, which has also been shown for correlated traits in the 1-sex literature (Lande & Arnold, 1983; Turelli, 1985; Jones et al., 2003; Chantepie & Chevin, 2020). This implies that, at equilibrium, the expected difference in trait means (signed sexual dimorphism) and expected intersex correlation are independent of each other (Figure 1A).

With a finite population, genetic drift generates random fluctuations in the sex-specific mean phenotypes. When sex-specific optima are far apart, these fluctuations can be neglected and sexual dimorphism is well-approximated by the difference in trait optima (for *r*_*fm*_ *<* 1 and, consequently, independent of intersex correlation. When trait optima coincide (or are close), however, random fluctuations can cause the expected *absolute value* of the difference in trait means (sexual dimorphism) to differ noticeably from their expected difference (signed sexual dimorphism) (Figure 1A-C, Figure S3A-C). When intersex correlation is high, drift-induced fluctuations in *SD* are *slow* (on the time-scale of molecular evolution as they are generated largely by the rare fixation of mutations with sex-specific effects) and when intersex correlation is low they are rapid (generated largely by small fluctuations in frequency of standing variation with sex-specific effects). Nevertheless, we show that in both cases *E*[*SD*] ≈ 1.6 · *δ* where *δ* is the typical deviation of the population mean from the (shared) optimum. Since this result holds for all *r*_*fm*_ *<* 1, we find that, consistent with classical work, the magnitude of difference in trait means (sexual dimorphism) and intersex correlation are independent at equilibrium. Nevertheless, the result is of interest because it suggests that nonzero sexual dimorphism is actually *expected* —even in the absence of selection for such.

The significance of this (or any) nonzero sexual dimorphism depends on how it compares to the scale of genetic variation in the trait. Consequently, whenever we make a point about the *magnitude* of sexual dimorphism, we choose to scale *SD* by the standard deviation of the phenotypic distribution(Figure 1D, Figure S3D). This standardization is often not done in empirical work, where sexual dimorphism is computed based on averages and ignoring variances (e.g. Lovich & Gibbons, 1992; Poissant et al., 2010). However, it is very hard to get a glimpse of the magnitude of sexual dimorphism, as well as compare across traits if their variances are not considered, since a given difference in sex-specific mean phenotypes is far more relevant for traits with lower standard deviations. We therefore encourage future work on sexual dimorphism to report variance-standardized measures of sex differences.

With respect to drift-induced *SD*, we show that when fluctuations of the population mean are relatively large (of magnitude 1*/*3 or 1*/*2 of the genetic standard deviation of the trait distribution) then, just by chance, trait means in the two sexes could differ by almost a full and a half phenotypic standard deviation, respectively. The effect is expected to be smaller for traits with smaller fluctuation in means relative to phenotypic variance (i.e. higher 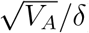 We do not know the empirical values of this quantity, so it is hard to predict the empirical relevance of the effect of drift on *SD* at equilibrium. However, we expect it to be of particular significance for traits that are more drift-sensitive, and where sex-specific optima are close to one another (Figure 1B), such as gene expression. We encourage future studies on sexual dimorphism in drift-sensitive traits to consider the possibility that, even fairly large, nonzero *SD* might not reflect the action of natural selection.

The two hypotheses most commonly considered in the literature with the potential to explain the negative correlation between intersex correlation and sexual dimorphism involve dynamic properties of the system, so we explored them by looking at the out-of-equilibrium dynamics of sex-specific adaptation under directional selection. The first hypothesis, discussed in Section 3.2.1, predicts higher levels of sexual dimorphism if intersex correlation is initially lower. We find that this can hold –– transiently and, importantly, given some additional assumptions. This is because, while intersex correlation does not determine the ultimate realized sexual dimorphism, it does determine the rate at which it evolves. Concretely, as Lande (1980) described, the rates of sexually-concordant vs sexually-dimorphic evolution are proportional to 1 + *r*_*fm*_ and 1 − *r*_*fm*_ (Equations 43 and 44, Figure 2C,D), and therefore evolve over two very different timescales for high intersex correlation. This result illustrates the quantitative constraint that *r*_*fm*_ imposes on the evolution of sex differences, and supports the idea that, after a limited time, the expected realized sexual dimorphism negatively correlates with intersex correlation (Bolnick & Doebeli, 2003), partially validating this first hypothesis.

However, this result requires two additional considerations: first, the fact that the constraint that *r*_*fm*_ poses on the evolution of *SD* is much weaker for a multigenic genetic architecture than the analytical results derived for the infinitesimal case suggest (Figure 2D,G; Figure 3). This is important because it offers an alternative explanation for the disparities in *SD* evolution timescales across experimental designs. Concretely, our results suggest that a more rapid evolution of *SD* for some traits (as in e.g. Bird & Schaffer, 1972) compared to others (as in e.g. Stewart & Rice, 2018) might not arise solely due to a higher *r*_*fm*_ in the less rapidly evolving traits, as is often discussed. It could also come about if the more rapidly evolving traits have a less infinitesimal genetic architecture—even when both sets of traits have similar similar *r*_*fm*_ levels. Second, as we show, a higher intersex correlation not only constrains *divergent* but also *convergent* evolution between the sexes, and the latter has the potential to generate the opposite pattern of a positive relationship between intersex correlation and sexual dimorphism (Figure 3). So the hypothesis that a negative correlation between intersex correlation and sexual dimorphism arises because sexually-dimorphic evolution is less constrained with lower *r*_*fm*_ requires some additional assumptions: i) that traits are sexually-dimorphically adapting under directional selection and ii) that sexually-dimorphic adaptation is more commonly divergent than convergent. In addition, its effect is weaker for traits with a less infinitesimal genetic architecture. The plausibility of the assumptions as well as our expectations surrounding genetic architecture are discussed below.

The second hypothesis, discussed in Section 3.2.2, supports the idea that a negative correlation between intersex correlation and extent of sex differences arises as a consequence of sexually-dimorphic adaptation involving an accumulation of sex-specific mutations leading to a decrease in *r*_*fm*_ over time. This idea traces back to D. B. Wright (1993) and Lande (1980, 1987) and, since neither author provides a mathematical justification, it seems rather based on an intuition of how such a process should evolve. Indeed, we find that (for a trait with a non-infinitesimal genetic architecture) intersex correlation decreases due to an increase in sex-specific variances, but not covariance, during sexually-dimorphic adaptation (Figure 4).

These changes in the (co)variance matrix are only transient; stabilizing selection translates the allele frequency changes between alleles with effects that are aligned and opposed to the phenotypic shift generated by directional selection into differences in fixation probabilities. In time, the transient increase in (co)variances ceases, and their equilibrium values are restored. In addition, the same transient decrease in intersex correlation is expected for divergent as well as convergent evolution (Figure 4).

This suggests that sexual dimorphism can evolve without long-term changes in *r*_*fm*_, as already discussed by Reeve and Fairbairn (2001). They too illustrate how this transient increase in second order moments speeds up adaptation with a non-infinitesimal genetic architecture. We recapitulate their result, and additionally show that transient increases in third central moments have the opposite effect, in that they act to slow down phenotypic adaptation (Figure S11), as was previously shown for the 1-sex case (Hayward & Sella, 2022).

We also obtain predictions for the equilibration timescale. Hayward and Sella (2022) showed that equilibrium is restablished over a time frame of the order of 2*N* generations, which we find to hold for equilibration under sexually-concordant adaptation regardless of the *r*_*fm*_ (Figure 2E). However, we find that higher intersex correlation delays equilibration when the population is sexually-dimorphically adapting for traits with an approximately infinitesimal genetic architecture (Figure 2F). Nevertheless, we find the effect of *r*_*fm*_ on equilibration time to be surprisingly small, given the constraint it poses on phenotypic evolution. This suggests some compensatory process: while, with high *r*_*fm*_, the amount of mutations contributing to sexually-dimorphic adaptation is less than those contributing to sexually-concordant adaptation, they might individually be subject to stronger directional selection and fix faster, leading to a similar rate of equilibration for the two types of adaptation.

Our model predicts a transient reduction in *r*_*fm*_ due to a temporary increase in sex-specific variances, and not a permanent reduction due to a decrease in between-sex covariance, as the common intuition seems to suggest (verbal arguments tracing back to Fisher, 1958 and Lande, 1980). These two results seem easy to reconcile by noting that, while we assumed that genetic architecture remains stable, as do most models of sex-specific adaptation (Reeve & Fairbairn, 2001; Bolnick & Doebeli, 2003; Connallon & Clark, 2014a, 2014b; Muralidhar & Coop, 2024), the general intuition seems to suggest an evolving genetic architecture (Lande, 1980; D. B. Wright, 1993; Bonduriansky & Rowe, 2005). Indeed, there are many different mechanisms that can lead to sexual conflict resolution: sex-specific expression of autosomal loci, via sex-linked modifiers or alternative splicing mechanisms (McIntyre et al., 2006; Stewart et al., 2010; Pennell & Morrow, 2013; Singh & Agrawal, 2023), gene duplication followed by sex-specific regulation of the paralogues (Rice & Chippindale, 2002; Proulx & Phillips, 2006; Sison-Mangus et al., 2006; Connallon & Clark, 2011), genomic imprinting (Day & Bonduriansky, 2004) and sex-dependent dominance of antagonistic alleles (Kidwell et al., 1977; Barson et al., 2015). The evolution of some of these mechanisms (e.g. a target for sex-hormone regulation or moving to a sex chromosome) would involve changes in the genetic architecture, i.e. leading to a higher proportion of sex-specific mutations underlying its expression – which we assume to be constant through time in our model. Some of those sex-specific changes in genetic architecture are expected to accelerate the rate of sexual dimorphism evolution and would likely drive more permanent reduction in *r*_*fm*_ as sexual dimorphism evolves (D. B. Wright, 1993; Bonduriansky & Rowe, 2005), contributing to a more stable negative correlation between sexual dimorphism and *r*_*fm*_ (Williams & Carroll, 2009; Stewart et al., 2010). However, they are likely to be slow (Williams & Carroll, 2009; Stewart et al., 2010; Bonduriansky & Chenoweth, 2009), probably occurring at an extra phase to the two described by Lande (1980, 1987) (and that we reproduce here) for sexually-concordant and -dimorphic adaptation with a constant genetic architecture, as suggested by D. B. Wright (1993). Looking at the dynamics with a non-changing genetic architecture is a useful first step that likely reflects the most likely genetic changes over the timescale of most experimental studies (e.g. Bird & Schaffer, 1972; Reeve & Fairbairn, 1996; Stewart & Rice, 2018). However, incorporating the option for an evolving genetic architecture in our model, involving changes in *h*(*ϕ*_*α*_) leading to a higher or lower proportion of sex-specific vs shared mutations, would be an easy and logical next step which would allow us to explore questions such as the conditions in which a slower pace in sexual dimorphism evolution could somehow ‘incentivize’ a more permanent reduction in *r*_*fm*_ involving changes in the genetic architecture, and whether these changes are expected to be partially restored after sexual conflict has been resolved.

Data can help shed light into whether sexually-dimorphic evolution typically implies changes in the genetic architecture leading to a permanent reduction in *r*_*fm*_, or whether it often relies on the current genetic architecture. Empirical studies report a general negative association between both (e.g. Poissant et al., 2010; Griffin et al., 2013), but this is far from universal, with many studies finding only weak or even absent associations (e.g. Cowley & Atchley, 1988; Ashman & Majetic, 2006). Thus, although we have evidence of sexual conflict resolution having relied on various types of changes in the genetic architecture (Delph et al., 2011; A. E. Wright et al., 2018), the fact that this pattern is inconsistent suggests that it might in many cases be more likely to reflect the transient dynamics under constant genetic architecture that we illustrate in this study. Dynamics which, importantly, only lead to the predicted negative correlation *given certain assumptions*.

The main assumptions to consider with respect to the two common hypotheses are three. First, that a good fraction of traits are out of equilibrium, adapting under (at least partially) sex-specific selection. This is because, as we have shown, a negative association between *r*_*fm*_ and sexual dimorphism due to directional selection arises only from transient dynamics of sexually-dimorphic adaptation to sex-specific selection. Given the high prevalence of sex-specific selection (Cox & Calsbeek, 2009) and the long time frame for sexual dimorphism evolution, specially for traits with high *r*_*fm*_ and an approximately infinitesimal genetic architecture (Equations 43, Figure 2C,D), it seems likely that many traits are subject to (sex-specific) directional selection.

The second assumption is that this sex-specific directional selection is more often divergent than convergent. This is required because under both hypotheses we predict that the correlation between *r*_*fm*_ and sex differences is negative with divergent but positive with convergent sex-specific adaptation (Figure 3; Figure 4F). However, selection for convergent evolution has also been reported for some traits and species (Owens & Hartley, 1998; Bonduriansky, 2006; Chursina, 2019; Lassek & Gaulin, 2022), indicating that this assumption might not generally hold. In general, there is no good reason to expect that one would occur more often than the other.

The third assumption concerns the genetic architecture of the trait considered, with the two hypotheses having strongest effects with opposite genetic architectures. On the one hand, although not strictly required, we expect H_1_ to be a more significant factor for traits with an approximately infinitesimal architecture. This is because the constraint that *r*_*fm*_ imposes on sexually-dimorphic adaptation, and therefore the potential to generate the expected negative correlation, is stronger for approximately infinitesimal traits. On the other hand, we show that the transient reduction in *r*_*fm*_ with sexually-dimorphic adaptation illustrated in H_2_ only occurs with non-infinitesimal traits since, with an infinitesimal genetic architecture, the phenotypic distribution remains unchanged under directional selection. The presence of large-effect mutations seems to be the rule for most complex traits, as suggested by GWAS (e.g. Wood et al., 2014; Locke et al., 2015; Simons et al., 2018), so most traits seem to deviate from the infinitesimal regime.

Other aspects of the genetic architecture are important to consider in the dynamics of (sex-specific) adaptation. The specific choice of sex-specificity of individual mutations can impact the evolutionary outcome (Rhen, 2000). In this case, we are sampling overall squared effect sizes from an exponential distribution, which seems to be a popular option (e.g. Connallon & Clark, 2014b), and define them as shared or sex-specific, with equal probabilities of being female- or male-specific. This choice is common in similar studies (Rhen, 2000; Reeve & Fairbairn, 2001; Bolnick & Doebeli, 2003) and is partially based on empirical evidence that sex-biased and sexually-antagonistic mutations, with phenotypic effects of different magnitudes and of different signs across sexes, respectively, should be rare (Dimas et al., 2012; Oliva et al., 2020; Puixeu et al., unpublished). However, running simulations that incorporate such mutations is likely to be a relevant extension to our work, since theoretical studies suggest that they can have a substantial contribution to phenotypic adaptation (Connallon & Clark, 2014a; Muralidhar & Coop, 2024). Also, we assume that the effect size distribution of new mutations is symmetric across sexes, and that this is independent of the effect size, meaning that large-effect mutations are equally likely to be female- or male-biased. However, there is empirical evidence of a male bias in fitness effects of spontaneous mutations in *Drosophila* (Mallet et al., 2011; Sharp & Agrawal, 2013), which might be worth considering in further extensions of this work.

Also importantly, previous work has shown that even with perfect intersex correlation and sexually concordant selection, sexual dimorphism can evolve if sex-specific genetic variances are unequal (Lynch & Walsh, 1998; Connallon & Clark, 2014b; Houle & Cheng, 2021). This suggests that interpreting *r*_*fm*_ as a constraint, which is the narrative employed in this manuscript, as well as many others cited throughout, relies on the assumptions that all genetic variance is additive, and that variances do not differ between the sexes (Lynch & Walsh, 1998; Bonduriansky & Chenoweth, 2009). These aspects illustrate the importance of clearly stating the assumptions underlying the chosen models, as they may lead to qualitatively different results, and support the idea that differences in genetic architecture are likely to account for a big part of the differences in the evolutionary dynamics of sexual dimorphism that have been observed across species and traits.

In summary, our work provides an in-depth examination of the relationship between intersex correlation and sex differences as well as their joint evolutionary dynamics in a population adapting to a sex-specific shift in optima under sex-specific stabilizing selection, mutation and drift, assuming non-evolving genetic architecture. It represents, to our knowledge, the first comprehensive account of various mechanisms that can generate a negative association between intersex correlation and sexual dimorphism, formalizing common intuition in the field. Also, it stresses the importance of revisiting commonly-used verbal arguments and illustrates how contextualizing their underlying assumptions can provide insightful information on the evolutionary forces shaping empirical patterns.

## Acknowledgments

We thank Tim Connallon for useful discussions and correspondence, Himani Sachdeva and Nick Barton for comments on the manuscript and the Scientific Computing unit at ISTA for technical support.

GP is the recipient of a DOC Fellowship of the Austrian Academy of Sciences at the Institute of Science and Technology Austria (DOC 25817) and received funding from the European Union’s Horizon 2020 research and innovation program under the Marie Sk?lodowska-Curie Grant (agreement no. 665385). LH received funding from the European Research Council, under the HaplotypeStructure Grant (grant no. 101055327) to Nick Barton.

## Author Contributions

GP and LH conceived the study, performed the analyses and wrote the manuscript.

## References

Abbott, J. K., Bedhomme, S., & Chippindale, A. K. (2010, September). Sexual conflict in wing size and shape in drosophila melanogaster. Journal of Evolutionary Biology, 23 (9), 1989–1997. doi: 10.1111/j.1420-9101.2010.02064.x

Ashman, T.-L. (2003, September). Constraints on the evolution of males and sexual dimor-phism: field estimates of genetic architecture of reproductive traits in three populations of gynodioecious fragaria virginiana. Evolution; International Journal of Organic Evolution, 57 (9), 2012–2025. doi: 10.1111/j.0014-3820.2003.tb00381.x

Ashman, T.-L., & Majetic, C. J. (2006, May). Genetic constraints on floral evolution: a review and evaluation of patterns. Heredity, 96 (5), 343–352. doi: 10.1038/sj.hdy.6800815

Barson, N. J., Aykanat, T., Hindar, K., Baranski, M., Bolstad, G. H., Fiske, P., Primmer, C. R. (2015, December). Sex-dependent dominance at a single locus maintains variation in age at maturity in salmon. Nature, 528 (7582), 405–408. Retrieved 2023-05-10, from https://www.nature.com/articles/nature16062 (Number: 7582 Publisher: Nature Publishing Group) doi: 10.1038/nature16062

Barton, N. H., Etheridge, A. M., & Véber, A. (2017, December). The infinitesimal model: definition, derivation, and implications. Theoretical Population Biology, 118, 50–73. doi: 10.1016/j.tpb.2017.06.001

Bird, M. A., & Schaffer, H. E. (1972, November). A study of the genetic basis of the sexual dimorphism for wing length in drosophila melanogaster. Genetics, 72 (3), 475–487. doi: 10.1093/genetics/72.3.475

Bolnick, D. I., & Doebeli, M. (2003, November). Sexual dimorphism and adaptive speciation: two sides of the same ecological coin. Evolution; International Journal of Organic Evolution, 57 (11), 2433–2449. doi: 10.1111/j.0014-3820.2003.tb01489.x

Bonduriansky, R. (2006, May). Convergent evolution of sexual shape dimorphism in diptera. Journal of Morphology, 267 (5), 602–611. doi: 10.1002/jmor.10426

Bonduriansky, R., & Chenoweth, S. F. (2009, May). Intralocus sexual conflict. Trends in Ecology & Evolution, 24 (5), 280–288. doi: 10.1016/j.tree.2008.12.005

Bonduriansky, R., & Rowe, L. (2005, September). Intralocus sexual conflict and the genetic architecture of sexually dimorphic traits in prochyliza xanthostoma (diptera: Piophilidae). Evolution; International Journal of Organic Evolution, 59 (9), 1965–1975.

Bürger, R., & Lande, R. (1994, November). On the distribution of the mean and variance of a quantitative trait under mutation-selection-drift balance. Genetics, 138 (3), 901–912. Retrieved 2023-05-28, from https://www.ncbi.nlm.nih.gov/pmc/articles/PMC1206237/

Chantepie, S., & Chevin, L.-M. (2020, December). How does the strength of selection influence genetic correlations? Evolution Letters, 4 (6), 468–478. doi: 10.1002/evl3.201

Chenoweth, S. F., & Blows, M. W. (2003). Signal trait sexual dimorphism and mutual sexual selection in drosophila serrata. Evolution, 57 (10), 2326–2334. Retrieved 2023-04-27, from https://onlinelibrary.wiley.com/doi/abs/10.1111/j.0014-3820.2003.tb00244.x ( eprint: https://onlinelibrary.wiley.com/doi/pdf/10.1111/j.0014-3820.2003.tb00244.x) doi: 10.1111/j.0014-3820.2003.tb00244.x

Cheverud, J. M., Dow, M. M., & Leutenegger, W. (1985). The quantitative assessment of phylogenetic constraints in comparative analyses: sexual dimorphism in body weight among primates. Evolution, 39 (6), 1335–1351. Retrieved 2023-05-05, from https://onlinelibrary.wiley.com/doi/abs/10.1111/j.1558-5646.1985.tb05699.x ( eprint: https://onlinelibrary.wiley.com/doi/pdf/10.1111/j.1558-5646.1985.tb05699.x) doi: 10.1111/j.1558-5646.1985.tb05699.x

Chursina, M. A. (2019, August). Convergent evolution of sexual dimorphism in species of the family dolichopodidae (diptera). Biodiversitas Journal of Biological Diversity, 20 (9). Retrieved 2023-05-10, from https://smujo.id/biodiv/article/view/4066 (Number: 9) doi: 10.13057/biodiv/d200908

Connallon, T., & Clark, A. G. (2011, March). The resolution of sexual antagonism by gene duplication. Genetics, 187 (3), 919–937. Retrieved 2023-05-10, from https://www.ncbi.nlm.nih.gov/pmc/articles/PMC3063682/ doi: 10.1534/genetics.110.123729

Connallon, T., & Clark, A. G. (2014a, July). Balancing selection in species with separate sexes: insights from fisher’s geometric model. Genetics, 197 (3), 991–1006. doi: 10.1534/genetics.114.165605

Connallon, T., & Clark, A. G. (2014b, February). Evolutionary inevitability of sexual antagonism. Proceedings. Biological Sciences, 281 (1776), 20132123. doi: 10.1098/rspb.2013.2123

Cowley, D. E., & Atchley, W. R. (1988, June). Quantitative genetics of drosophila melanogaster. ii. heritabilities and genetic correlations between sexes for head and thorax traits. Genetics, 119 (2), 421–433. doi: 10.1093/genetics/119.2.421

Cox, R. M., & Calsbeek, R. (2009, February). Sexually antagonistic selection, sexual dimor-phism, and the resolution of intralocus sexual conflict. The American Naturalist, 173 (2), 176–187. doi: 10.1086/595841

Cox, R. M., Cox, C. L., McGlothlin, J. W., Card, D. C., Andrew, A. L., & Castoe, T. A. (2017, March). Hormonally mediated increases in sex-biased gene expression accompany the breakdown of between-sex genetic correlations in a sexually dimorphic lizard. The American Naturalist, 189 (3), 315–332. doi: 10.1086/690105

Day, T., & Bonduriansky, R. (2004, August). Intralocus sexual conflict can drive the evolution of genomic imprinting. Genetics, 167 (4), 1537–1546. doi: 10.1534/genetics.103.026211

Delph, L. F., Arntz, A. M., Scotti-Saintagne, C., & Scotti, I. (2010, October). The genomic architecture of sexual dimorphism in the dioecious plant silene latifolia. Evolution; International Journal of Organic Evolution, 64 (10), 2873–2886. doi: 10.1111/j.1558-5646.2010.01048.x

Delph, L. F., Frey, F. M., Steven, J. C., & Gehring, J. L. (2004). Investigating the independent evolution of the size of floral organs via g-matrix estimation and artificial selection. Evolution & Development, 6 (6), 438–448. doi: 10.1111/j.1525-142X.2004.04052.x

Delph, L. F., Steven, J. C., Anderson, I. A., Herlihy, C. R., & Brodie, E. D. (2011, October). Elimination of a genetic correlation between the sexes via artificial correlational selection. Evolution; International Journal of Organic Evolution, 65 (10), 2872–2880. doi: 10.1111/j.1558-5646.2011.01350.x

Dimas, A. S., Nica, A. C., Montgomery, S. B., Stranger, B. E., Raj, T., Buil, A., … Dermitzakis, E. T. (2012, December). Sex-biased genetic effects on gene regulation in humans. Genome Research, 22 (12), 2368–2375. doi: 10.1101/gr.134981.111

Eisen, E. J., & Hanrahan, J. P. (1972, October). Selection for sexual dimorphism in body weight of mice. Australian Journal of Biological Sciences, 25 (5), 1015–1024. doi: 10.1071/bi9721015

Fairbairn, D. J. (2007, July). Sexual dimorphism in the water strider, aquarius remigis: a case study of adaptation in response to sexually antagonistic selection. In D. J. Fairbairn, W. U. Blanckenhorn, & T. Székely (Eds.), Sex, Size and Gender Roles: Evolutionary Studies of Sexual Size Dimorphism (p. 0). Oxford University Press. Retrieved 2024-04-30, from https://doi.org/10.1093/acprof:oso/9780199208784.003.0011 doi: 10.1093/acprof:oso/9780199208784.003.0011

Fisher, R. (1958). The genetical theory of natural selection (Second edition ed.). New York: dover.

Frankham, R. (1968a, December). Sex and selection for a quantitative character in drosophila. ii. the sex dimorphism. Australian Journal of Biological Sciences, 21 (6), 1225–1237. doi: 10.1071/bi9681225

Frankham, R. (1968b, December). Sex and selection for a quantitative character in drosophila. i. single-sex selection. Australian Journal of Biological Sciences, 21 (6), 1215–1223. doi: 10.1071/bi9681215

Griffin, R. M., Dean, R., Grace, J. L., Rydén, P., & Friberg, U. (2013, September). The shared genome is a pervasive constraint on the evolution of sex-biased gene expression. Molecular Biology and Evolution, 30 (9), 2168–2176. doi: 10.1093/molbev/mst121

Harrison, B. J. (1953, August). Reversal of a secondary sex character by selection. Heredity, 7 (2), 153–164. Retrieved 2023-04-27, from https://www.nature.com/articles/hdy195324 (Number: 2 Publisher: Nature Publishing Group) doi: 10.1038/hdy.1953.24

Hayward, L. K., & Sella, G. (2022, September). Polygenic adaptation after a sudden change in environment. eLife, 11, e66697. doi: 10.7554/eLife.66697

Houle, D., & Cheng, C. (2021, May). Predicting the evolution of sexual dimorphism in gene expression. Molecular Biology and Evolution, 38 (5), 1847–1859. doi: 10.1093/molbev/msaa329

Jones, A. G., Arnold, S. J., & Bürger, R. (2003). Stability of the g-matrix in a population experiencing pleiotropic mutation, stabilizing selection, and genetic drift. Evolution, 57 (8), 1747–1760. Retrieved 2023-05-10, from https://www.jstor.org/stable/3448700 (Publisher: [Society for the Study of Evolution, Wiley])

Kaufmann, P., Wolak, M. E., Husby, A., & Immonen, E. (2021, October). Rapid evolution of sexual size dimorphism facilitated by y-linked genetic variance. Nature Ecology & Evolution, 5 (10), 1394–1402. doi: 10.1038/s41559-021-01530-z

Kidwell, J. F., Clegg, M. T., Stewart, F. M., & Prout, T. (1977, January). Regions of stable equilibria for models of differential selection in the two sexes under random mating. Genetics, 85 (1), 171–183. doi: 10.1093/genetics/85.1.171

Lande, R. (1976, June). Natural selection and random genetic drift in phenotypic evolution. Evolution; International Journal of Organic Evolution, 30 (2), 314–334. doi: 10.1111/j.1558-5646.1976.tb00911.x

Lande, R. (1980, March). Sexual dimorphism, sexual selection, and adaptation in polygenic characters. Evolution; International Journal of Organic Evolution, 34 (2), 292–305. doi: 10.1111/j.1558-5646.1980.tb04817.x

Lande, R. (1987). Genetic correlations between the sexes in the evolution of sexual dimorphism and mating preferences. In J. W. Bardbury & M. B. Andersson (Eds.), Sexual Selection: Testing the Alternatives : Report of the Dahlem Workshop on Sexual Selection: Testing the Alternatives, Berlin 1986, August 31-September 5. Wiley. (Google-Books-ID: Z9lqAAAAMAAJ)

Lande, R., & Arnold, S. J. (1983). The measurement of selection on correlated characters. Evolution, 37 (6), 1210–1226. Retrieved 2023-05-10, from https://www.jstor.org/stable/2408842 (Publisher: [Society for the Study of Evolution, Wiley]) doi: 10.2307/2408842

Lassek, W. D., & Gaulin, S. J. C. (2022). Substantial but misunderstood human sexual dimorphism results mainly from sexual selection on males and natural selection on females. Frontiers in Psychology, 13. Retrieved 2023-05-10, from https://www.frontiersin.org/articles/10.3389/fpsyg.2022.859931

Leinonen, T., Cano, J. M., & Merilä, J. (2011, February). Genetic basis of sexual dimorphism in the threespine stickleback gasterosteus aculeatus. Heredity, 106 (2), 218–227. doi: 10.1038/hdy.2010.104

Locke, A. E., Kahali, B., Berndt, S. I., Justice, A. E., Pers, T. H., Day, F. R., … Speliotes, E. K. (2015, February). Genetic studies of body mass index yield new insights for obesity biology. Nature, 518 (7538), 197–206. Retrieved 2023-05-10, from https://www.nature.com/articles/nature14177 (Number: 7538 Publisher: Nature Publishing Group) doi: 10.1038/nature14177

Lovich, J. E., & Gibbons, J. W. (1992). A review of techniques for quantifying sexual size dimorphism. Growth, development, and aging: GDA, 56 (4), 269–281.

Lynch, M., & Walsh, B. (1998). Genetics and analysis of quantitative traits (1st edition ed.). Sunderland, Mass: Sinauer Associates is an imprint of Oxford University Press.

Mallet, M. A., Bouchard, J. M., Kimber, C. M., & Chippindale, A. K. (2011, June). Experimental mutation-accumulation on the x chromosome of drosophila melanogaster reveals stronger selection on males than females. BMC Evolutionary Biology, 11 (1), 156. Retrieved 2023-05-10, from https://doi.org/10.1186/1471-2148-11-156 doi: 10.1186/1471-2148-11-156

McDaniel, S. F. (2005, November). Genetic correlations do not constrain the evolution of sexual dimorphism in the moss ceratodon purpureus. Evolution; International Journal of Organic Evolution, 59 (11), 2353–2361.

McGlothlin, J. W., Cox, R. M., & Brodie, E. D. (2019, July). Sex-specific selection and the evolution of between-sex genetic covariance. The Journal of Heredity, 110 (4), 422–432. doi: 10.1093/jhered/esz031

McIntyre, L. M., Bono, L. M., Genissel, A., Westerman, R., Junk, D., Telonis-Scott, M., … Nuzhdin, S. V. (2006, August). Sex-specific expression of alternative transcripts in drosophila. Genome Biology, 7 (8), R79. Retrieved 2023-05-10, from https://doi.org/10.1186/gb-2006-7-8-r79 doi: 10.1186/gb-2006-7-8-r79

Morrow, E. H. (2015). The evolution of sex differences in disease. Biology of Sex Differences, 6, 5. doi: 10.1186/s13293-015-0023-0

Morrow, E. H., & Connallon, T. (2013, December). Implications of sex-specific selection for the genetic basis of disease. Evolutionary Applications, 6 (8), 1208–1217. doi: 10.1111/eva.12097

Muralidhar, P., & Coop, G. (2024, February). Polygenic response of sex chromosomes to sexual antagonism. Evolution; International Journal of Organic Evolution, 78 (3), 539–554. doi: 10.1093/evolut/qpad231

Oliva, M., Mun∼oz-Aguirre, M., Kim-Hellmuth, S., Wucher, V., Gewirtz, A. D. H., Cotter, D. J., … Stranger, B. E. (2020, September). The impact of sex on gene expression across human tissues. Science (New York, N.Y.), 369 (6509), eaba3066. doi: 10.1126/science.aba3066

Owens, I. P. F., & Hartley, I. R. (1998, March). Sexual dimorphism in birds: why are there so many different forms of dimorphism? Proceedings of the Royal Society of London. Series B: Biological Sciences, 265 (1394), 397–407. Retrieved 2023-05-10, from https://royalsocietypublishing.org/doi/10.1098/rspb.1998.0308 doi: 10.1098/rspb.1998.0308

Pennell, T. M., & Morrow, E. H. (2013). Two sexes, one genome: the evolutionary dynamics of intralocus sexual conflict. Ecology and Evolution, 3 (6), 1819–1834. Retrieved 2023-05-10, from https://onlinelibrary.wiley.com/doi/abs/10.1002/ece3.540 ( eprint: https://onlinelibrary.wiley.com/doi/pdf/10.1002/ece3.540) doi: 10.1002/ece3.540

Poissant, J., Wilson, A. J., & Coltman, D. W. (2010, January). Sex-specific genetic variance and the evolution of sexual dimorphism: a systematic review of cross-sex genetic correlations. Evolution; International Journal of Organic Evolution, 64 (1), 97–107. doi: 10.1111/j.1558-5646.2009.00793.x

Prasad, N. G., Bedhomme, S., Day, T., & Chippindale, A. K. (2007, January). An evolutionary cost of separate genders revealed by male-limited evolution. The American Naturalist, 169 (1), 29–37. doi: 10.1086/509941

Preziosi, R. F., & Roff, D. A. (1998, July). Evidence of genetic isolation between sexually monomorphic and sexually dimorphic traits in the water strider aquarius remigis. Heredity, 81 (1), 92–99. Retrieved 2023-04-27, from https://www.nature.com/articles/6883800 (Number: 1 Publisher: Nature Publishing Group) doi: 10.1046/j.1365-2540.1998.00380.x

Proulx, S. R., & Phillips, P. C. (2006). Allelic divergence precedes and promotes gene duplication. Evolution, 60 (5), 881–892. Retrieved 2023-05-10, from https://www.jstor.org/stable/4095392 (Publisher: [Society for the Study of Evolution, Wiley])

Puixeu, G., Pickup, M., Field, D. L., & Barrett, S. C. H. (2019, November). Variation in sexual dimorphism in a wind-pollinated plant: the influence of geographical context and life-cycle dynamics. The New Phytologist, 224 (3), 1108–1120. doi: 10.1111/nph.16050

Reeve, J. P., & Fairbairn, D. J. (1996, October). Sexual size dimorphism as a correlated response to selection on body size: an empirical test of the quantitative genetic model. Evolution; International Journal of Organic Evolution, 50 (5), 1927–1938. doi: 10.1111/j.1558-5646.1996.tb03580.x

Reeve, J. P., & Fairbairn, D. J. (2001). Predicting the evolution of sexual size dimorphism. Journal of Evolutionary Biology, 14 (2), 244–254. Retrieved 2023-04-27, from https://onlinelibrary.wiley.com/doi/abs/10.1046/j.1420-9101.2001.00276.x ( eprint: https://onlinelibrary.wiley.com/doi/pdf/10.1046/j.1420-9101.2001.00276.x) doi: 10.1046/j.1420-9101.2001.00276.x

Rhen, T. (2000, February). Sex-limited mutations and the evolution of sexual dimorphism. Evolution; International Journal of Organic Evolution, 54 (1), 37–43. doi: 10.1111/j.0014-3820.2000.tb00005.x

Rice, W. R. (1984, July). Sex chromosomes and the evolution of sexual dimorphism. Evolution; International Journal of Organic Evolution, 38 (4), 735–742. doi: 10.1111/j.1558-5646.1984.tb00346.x

Rice, W. R., & Chippindale, A. K. (2001). Intersexual ontogenetic conflict. Journal of Evolutionary Biology, 14 (5), 685–693. Retrieved 2023-04-27, from https://onlinelibrary.wiley.com/doi/abs/10.1046/j.1420-9101.2001.00319.x ( eprint: https://onlinelibrary.wiley.com/doi/pdf/10.1046/j.1420-9101.2001.00319.x) doi: 10.1046/j.1420-9101.2001.00319.x

Rice, W. R., & Chippindale, A. K. (2002, November). The evolution of hybrid infertility: perpetual coevolution between gender-specific and sexually antagonistic genes. Genetica, 116 (2), 179–188. Retrieved 2023-05-10, from https://doi.org/10.1023/A:1021205130926 doi: 10.1023/A:1021205130926

Sanjak, J. S., Sidorenko, J., Robinson, M. R., Thornton, K. R., & Visscher, P. M. (2018, January). Evidence of directional and stabilizing selection in contemporary humans. Proceedings of the National Academy of Sciences of the United States of America, 115 (1), 151–156. doi: 10.1073/pnas.1707227114

Santiago, E. (1998, April). Linkage and the maintenance of variation for quantitative traits by mutation–selection balance: an infinitesimal model. Genetics Research, 71 (2), 161–170. Retrieved 2024-05-07, from https://www.cambridge.org/core/journals/genetics-research/article/linkage-and-the-maintenance-of-variation-for-quantitative-traits-by-mutationselection-balance-an-infinitesimal-model/2085C53021580D05FCA785172FEE0D5D doi: 10.1017/S0016672398003231

Sharp, N. P., & Agrawal, A. F. (2013, April). Male-biased fitness effects of spontaneous mutations in drosophila melanogaster. Evolution; International Journal of Organic Evolution, 67 (4), 1189–1195. doi: 10.1111/j.1558-5646.2012.01834.x

Simons, Y. B., Bullaughey, K., Hudson, R. R., & Sella, G. (2018, March). A population genetic interpretation of gwas findings for human quantitative traits. PLoS biology, 16 (3), e2002985. doi: 10.1371/journal.pbio.2002985

Singh, A., & Agrawal, A. F. (2023, May). Two forms of sexual dimorphism in gene expression in drosophila melanogaster: their coincidence and evolutionary genetics. Molecular Biology and Evolution, 40 (5), msad091. Retrieved 2023-05-10, from https://doi.org/10.1093/molbev/msad091 doi: 10.1093/molbev/msad091

Sison-Mangus, M. P., Bernard, G. D., Lampel, J., & Briscoe, A. D. (2006, August). Beauty in the eye of the beholder: the two blue opsins of lycaenid butterflies and the opsin gene-driven evolution of sexually dimorphic eyes. The Journal of Experimental Biology, 209 (Pt 16), 3079–3090. doi: 10.1242/jeb.02360

Stewart, A. D., Pischedda, A., & Rice, W. R. (2010). Resolving intralocus sexual conflict: genetic mechanisms and time frame. The Journal of Heredity, 101 Suppl 1 (Suppl 1), S94–99. doi: 10.1093/jhered/esq011

Stewart, A. D., & Rice, W. R. (2018, September). Arrest of sex-specific adaptation during the evolution of sexual dimorphism in drosophila. Nature Ecology & Evolution, 2 (9), 1507–1513. doi: 10.1038/s41559-018-0613-4

Stulp, G., Kuijper, B., Buunk, A. P., Pollet, T. V., & Verhulst, S. (2012, December). Intralocus sexual conflict over human height. Biology Letters, 8 (6), 976–978. doi: 10.1098/rsbl.2012.0590

Turelli, M. (1984, April). Heritable genetic variation via mutation-selection balance: Lerch’s zeta meets the abdominal bristle. Theoretical Population Biology, 25 (2), 138–193. doi: 10.1016/0040-5809(84)90017-0

Turelli, M. (1985, September). Effects of pleiotropy on predictions concerning mutationselection balance for polygenic traits. Genetics, 111 (1), 165–195. doi: 10.1093/genetics/111.1.165

Williams, T. M., & Carroll, S. B. (2009, November). Genetic and molecular insights into the development and evolution of sexual dimorphism. Nature Reviews. Genetics, 10 (11), 797–804. doi: 10.1038/nrg2687

Wood, A. R., Esko, T., Yang, J., Vedantam, S., Pers, T. H., Gustafsson, S., … Frayling, T. M. (2014, November). Defining the role of common variation in the genomic and biological architecture of adult human height. Nature Genetics, 46 (11), 1173–1186. Retrieved 2023-05-10, from https://www.nature.com/articles/ng.3097 (Number: 11 Publisher: Nature Publishing Group) doi: 10.1038/ng.3097

Wright, A. E., Fumagalli, M., Cooney, C. R., Bloch, N. I., Vieira, F. G., Buechel, S. D., … Mank, J. E. (2018, April). Male-biased gene expression resolves sexual conflict through the evolution of sex-specific genetic architecture. Evolution Letters, 2 (2), 52–61. doi: 10.1002/evl3.39

Wright, D. B. (1993). Evolution of sexually dimorphic characters in peccaries (mammalia, tayassuidae). Paleobiology, 19 (1), 52–70. Retrieved 2023-05-10, from https://www.cambridge.org/core/product/identifier/S0094837300012318/type/journal_article doi: 10.1017/S0094837300012318

Wright, S. (1935, March). The analysis of variance and the correlations between relatives with respect to deviations from an optimum. Journal of Genetics, 30 (2), 243–256. Retrieved 2023-05-10, from https://doi.org/10.1007/BF02982239 doi: 10.1007/BF02982239

Zwaan, B. J., Zijlstra, W. G., Keller, M., Pijpe, J., & Brakefield, P. M. (2008, December). Potential constraints on evolution: sexual dimorphism and the problem of protandry in the butterfly bicyclus anynana. Journal of Genetics, 87 (4), 395–405. doi: 10.1007/s12041-008-0062-y

