## Supplementary Material for "The relationship between sexual dimorphism and intersex correlation: do models support intuition?"

Gemma Puixeu[1*] and Laura Katharine Hayward[1‡*]

**1** Institute of Science and Technology Austria, Klosterneuburg, Lower Austria, Austria

‡Senior author

* (G Puixeu), (LK Hayward)

### Contents

**1 Derivation of the allelic equation** **3**

**2 Genetic variances and covariance at equilibrium** **5**
- 2.1 The number of segregating sites at equilibrium . . . . . 5
- 2.2 Computing the variances and covariance . . . . . 5
  - 2.2.1 The variances and covariance using our distribution of angles . 7

**3 Exploring the limits to shift size** **8**

**4 Types of simulations** **10**

**5 Empirical calculation of sex-specific variances, intersex covariance and intersex correlation** **13**

**6 Calculation of the total genetic variance, $V_{A,t}$** **15**

**7 The relationship between various measures of sexual dimorphism and $r_{fm}$ at equilibrium** **17**

**References** **29**

### 1 Derivation of the allelic equation

Consider an allele segregating at frequency $x_f$ and $x_m$ with phenotypic effect $a_f$ and $a_m$ in females and males, respectively, with an overall frequency $x \equiv (x_f + x_m)/2$ in the population. The expected change in the allele's overall frequency in a single generation is given by

$$E[\Delta x] = \frac{1}{2}\left(E[\Delta x_f] + E[\Delta x_m]\right), \tag{S.1}$$

with

$$E[\Delta x_j] = \frac{x_f x_m \bar{w}_0^j + \frac{1}{2}\left(x_f\left(1 - x_m\right) + x_m\left(1 - x_f\right)\right)\bar{w}_1^j}{\overline{w}^j} - x_j, \tag{S.2}$$

where

$$\overline{w}^j \equiv x_f x_m \bar{w}_0^j + \left(x_f\left(1 - x_m\right) + x_m\left(1 - x_f\right)\right)\bar{w}_1^j + \left(1 - x_f\right)\left(1 - x_m\right)\bar{w}_2^j \tag{S.3}$$

and $\bar{w}_c^j$ denotes the average fitness of females ($j = f$) and males ($j = m$) with $c = 0, 1$ or $2$ copies of focual the allele, where the averaging is over the distribution of contributions to the phenotype from other sites (Kidwell et al., 1977). It follows from an identical argument as that employed in Appendix 3, Section 1.1 of Hayward and Sella (2022) for the single sex case that, provided $V_{A,O} \ll V_S$, these average sex-specific fitnesses are well approximated by

$$\bar{w}_c^j \approx \mathrm{Exp}\left[-\frac{[D_j - a_j(c - 2x_j)]^2 \cdot \gamma_j^2}{V_S}\right], \tag{S.4}$$

where $D_f$ and $D_m$ are the distances of the male and female mean trait values from their respective optima.

Rewriting Equation S.2 in terms of the overall allele frequency, $x$, and the squared difference in allele frequency between males and females, $x_d^2 = (x_f - x_m)^2$, yields

$$\begin{aligned} E[\Delta x_j] =& \frac{x\left(1 - x\right)\left(x\left(\bar{w}_0^j - \bar{w}_1^j\right) + \left(1 - x\right)\left(\bar{w}_1^j - \bar{w}_2^j\right)\right)}{2\overline{w}^j} + \\ & \frac{x_d^2}{8\overline{w}^j}\left(\bar{w}_1^j - \bar{w}_0^j - x\left(2\bar{w}_1^j - \bar{w}_0^j - \bar{w}_2^j\right)\right) \text{ and} \end{aligned} \tag{S.5}$$

where

$$\overline{w}^j \equiv x^2\bar{w}_0^j + 2x\left(1-x\right)\bar{w}_1^j + \left(1-x\right)^2\bar{w}_2^j + \frac{x_d^2}{4}\left(2\bar{w}_1^j - \bar{w}_0^j - \bar{w}_2^j\right) \text{ and} \tag{S.6}$$

(Tim Connallon, personal correspondance).

We expect deviations from Hardy-Weinburg to be negligible and we neglect terms of $O\left(x_d^2\right)$, arriving at

$$E[\Delta x_j] \approx \frac{x\left(1-x\right)\left(x\left(\bar{w}_0^j - \bar{w}_1^j\right) + \left(1-x\right)\left(\bar{w}_1^j - \bar{w}_2^j\right)\right)}{2\overline{w}^j}, \tag{S.7}$$

with

$$\overline{w}^j \approx x^2\bar{w}_0^j + 2x\left(1-x\right)\bar{w}_1^j + \left(1-x\right)^2\bar{w}_2^j \tag{S.8}$$

for $j = f$ or $m$. Equation S.7 is identical in form to the expression for $E[\Delta x]$ for a single sex with a Gaussian fitness function. An identical derivation as that in Appendix 3, Section 1.1 of Hayward and Sella (2022), therefore shows that, provided $a \ll \sqrt{V_S}, V_{A,O} \ll V_S$ and $D_j \cdot \gamma_j \lessapprox \sqrt{V_S}$, Taylor expanding equation Equation S.7 and dropping higher order terms provides a good approximation, yielding

$$E[\Delta x_j] \approx \frac{a_j D_j (2\gamma_j^2)}{V_S}x(1-x) - \frac{a_j^2(2\gamma_j^2)}{V_S}x(1-x)\left(1/2 - x\right) \tag{S.9}$$

for $j = f$ or $m$. Consequently, the expected change in overall allele frequency in a single generation (Equation S.1) is well approximated by

$$E[\Delta x] \approx \left(\frac{a_f D_f \gamma_f^2}{V_S} + \frac{a_m D_m \gamma_m^2}{V_S}\right)x(1-x) - \left(\frac{a_f^2\gamma_f^2}{V_S} + \frac{a_m^2\gamma_m^2}{V_S}\right)x(1-x)\left(1/2 - x\right). \tag{S.10}$$

### 2 Genetic variances and covariance at equilibrium

#### 2.1 The number of segregating sites at equilibrium

At equilibrium the first two moments of change in allele frequency are given by

$$E_{eq}[\Delta x] = -\frac{a^2}{V_S} x(1-x)\left(1/2 - x\right) \tag{S.11}$$

$$V[\Delta x] \approx \frac{x(1-x)}{2N}, \tag{S.12}$$

where $a \equiv \sqrt{a_f^2\gamma_f^2 + a_m^2\gamma_m^2}$ is the overall phenotypic magnitude. Consequently, the density of sites segregating with overall phenotypic magnitude, $a$, and MAF $\tilde{x}$ per unit mutational input is

$$2\rho\left(a,\tilde{x}\right) \equiv \begin{cases} (2Nx)\cdot 4\cdot \mathrm{Exp}\left[-a^2\tilde{x}(1-\tilde{x})\right]/\left[\tilde{x}(1-\tilde{x})\right] & 0 \le \tilde{x} \le 1/(2N) \\ 4\cdot \mathrm{Exp}\left[-a^2\tilde{x}(1-\tilde{x})\right]/\left[\tilde{x}(1-\tilde{x})\right] & 1/(2N) < \tilde{x} \le 1/2 \end{cases} \tag{S.13}$$

(Hayward & Sella, 2022, Appendix 3, Section 3). The mutational input per generation of alleles with overall phenotypic magnitude, $a$, and angle, $\phi_a$ is $2NU\cdot g(a)\cdot h(\phi_a)$, where $g(a)$ is the distribution of incoming overall effect magnitudes and $h(\phi_a)$ is the distribution of incoming effect angles. It follows that the density of sites segregating with overall phenotypic magnitude, $a$, angle, $\phi_a$ , and MAF $\tilde{x}$ is

$$2NU\cdot 2\rho\left(a,\tilde{x}\right)\cdot h(\phi_a). \tag{S.14}$$

#### 2.2 Computing the variances and covariance

The genic variance in females ($j = f$), and males ($j = m$), and the covariance between the sexes (Equations 4 and 5 in the main text) can be rewritten as

$$V_{A,j} = \sum_i 2a_{i,j}^2\tilde{x}_i(1-\tilde{x}_i); \quad B = \sum_i 2a_{i,f}a_{i,m}\tilde{x}_i(1-\tilde{x}_i); \tag{S.15}$$

where, here, the $\tilde{x}_i$ are minor allele frequencies. Changing variables from sex-specific effects $a_{i,f}$ and $a_{i,m}$ to the overall phenotypic magnitude, $a_i$, and the angle, $\phi_{i,a}$

(Equations 14 and 15 in the main text), these expressions become

$$V_{A,f} = \frac{1}{\gamma_f^2} \sum_i 2a_i^2 \cos^2(\phi_{i,a}) \tilde{x}_i (1 - \tilde{x}_i), \tag{S.16}$$

$$V_{A,m} = \frac{1}{\gamma_m^2} \sum_i 2a_i^2 \sin^2(\phi_{i,a}) \tilde{x}_i (1 - \tilde{x}_i) \tag{S.17}$$

$$B = \frac{1}{\gamma_f \gamma_m} \sum_i 2a_i^2 \cos(\phi_{i,a}) \sin(\phi_{i,a}) \tilde{x}_i (1 - \tilde{x}_i) \tag{S.18}$$

Approximating the sums in Equations S.16, S.17 and S.18 by integrals over the density of segregating sites at equilibrium (Equation S.14), it follows that

$$V_{A,f} = \frac{1}{\gamma_f^2} \int_{\phi_a=0}^{2\pi} \int_{a=0}^{\infty} \int_{\tilde{x}=0}^{1/2} 2a^2 \cos^2(\phi_a) \tilde{x}(1 - \tilde{x}) [2NU \cdot 2\rho\,(a, \tilde{x})\, h(\phi_a)] d\tilde{x}\, da\, d\phi_a, \tag{S.19}$$

$$V_{A,m} = \frac{1}{\gamma_m^2} \int_{\phi_a=0}^{2\pi} \int_{a=0}^{\infty} \int_{\tilde{x}=0}^{1/2} 2a^2 \sin^2(\phi_a) \tilde{x}(1 - \tilde{x}) [2NU \cdot 2\rho\,(a, \tilde{x})\, h(\phi_a)] d\tilde{x}\, da\, d\phi_a \tag{S.20}$$

$$B = \frac{1}{\gamma_f \gamma_m} \int_{\phi_a=0}^{2\pi} \int_{a=0}^{\infty} \int_{\tilde{x}=0}^{1/2} 2a^2 \cos(\phi_a) \sin(\phi_a) \tilde{x}(1 - \tilde{x}) [2NU \cdot 2\rho\,(a, \tilde{x})\, h(\phi_a)] d\tilde{x}\, da\, d\phi_a \tag{S.21}$$

Rearranging integrals, these expressions can be rewritten as

$$V_{A,f} = \frac{1}{\gamma_f^2} \int_0^{2\pi} \cos^2(\phi_a) \cdot h(\phi_a) d\phi_a \cdot V_{A,O}, \tag{S.22}$$

$$V_{A,m} = \frac{1}{\gamma_m^2} \int_0^{2\pi} \sin^2(\phi_a) \cdot h(\phi_a) d\phi_a \cdot V_{A,O} \tag{S.23}$$

$$B = \frac{1}{\gamma_f \gamma_m} \int_0^{2\pi} \cos(\phi_a) \sin(\phi_a) \cdot h(\phi_a) d\phi_a \cdot V_{A,O} \tag{S.24}$$

where

$$V_{A,O} = 2NU \cdot \int_0^{\infty} \int_0^{1/2} 2a^2 \tilde{x}(1 - \tilde{x}) \cdot 2\rho\,(a, \tilde{x})\, d\tilde{x}\, da = 2NU \cdot \int_0^{\infty} v(a) g(a) da \tag{S.25}$$

with $v(a) = 4a \cdot D_+\,(a/2)$ and $D_+$ is the Dawson function (the second equality was shown in Hayward & Sella, 2022, Appendix 3, Section 3.2).

##### 2.2.1 The variances and covariance using our distribution of angles

Throughout the main text, we assume a very specific distribution of angles. Namely,

$$h_r(\phi_a) = \frac{(1-r)}{4}\left(\overbrace{\delta_0(\phi_a)+\delta_\pi(\phi_a)}^{\text{female specific}} + \overbrace{\delta_{\frac{\pi}{2}}(\phi_a)+\delta_{\frac{3\pi}{2}}(\phi_a)}^{\text{male specific}}\right) + \frac{r}{2}\overbrace{\left(\delta_{\frac{\pi}{4}}(\phi_a)+\delta_{\frac{5\pi}{4}}(\phi_a)\right)}^{\text{shared}}, \tag{S.26}$$

where $\delta_\theta$ is the delta distribution, i.e., $\int_\phi f(\phi)\delta_\theta(\phi)d\phi = f(\theta)$. Consequently,

$$\int_0^{2\pi} f(\phi_a)\cdot h_r(\phi_a)d\phi_a = \frac{(1-r)}{4}\left(f(0)+f(\pi)+f\left(\frac{\pi}{2}\right)+f\left(\frac{3\pi}{2}\right)\right) + \frac{r}{2}\left(f\left(\frac{\pi}{4}\right)+f\left(\frac{5\pi}{4}\right)\right) \tag{S.27}$$

so

$$\int_0^{2\pi} \cos^2(\phi_a)\cdot h_r(\phi_a)d\phi_a = (1-r)/4\cdot 2 + r/2\cdot 1 = 1/2 \tag{S.28}$$

$$\int_0^{2\pi} \sin^2(\phi_a)\cdot h_r(\phi_a)d\phi_a = (1-r)/4\cdot 2 + r/2\cdot 1 = 1/2 \tag{S.29}$$

$$\int_0^{2\pi} \cos(\phi_a)\sin(\phi_a)\cdot h_r(\phi_a)d\phi_a = (1-r)/4\cdot 0 + r/2\cdot 1 = r/2, \tag{S.30}$$

and it follows from Equations S.22 to S.24 that, with $\gamma_f = \gamma_m = 1/\sqrt{2}$ ,

$$V_{A,f} = V_{A,O}, \quad V_{A,m} = V_{A,O}, \quad \text{and} \quad B = r\cdot V_{A,O}. \tag{S.31}$$

### 3 Exploring the limits to shift size

We explore the range of magnitudes of shifts in sex-specific optima $\Lambda_f, \Lambda_m$ that fulfill the assumptions of the analytical framework. Concretely, we ensure that the shifts are substantially larger than the random fluctuations so that we can see a signal ($|\Lambda_f|, |\Lambda_m| > \delta$), but smaller than, or on the order of, the width of the fitness function ($|\Lambda_f|, |\Lambda_m| \lessapprox \sqrt{V_S}$). To test the limits to the shift size, we computed the average of sex-specific variances divided by the analytical variance for various sizes in shift size. As Figure S1 shows, with completely shared genetic architecture between sexes ($r = 1$ so $\phi_a = \pi/4$ or $5 \cdot \pi/4$), the realized variance remains as expected for shift sizes (scaled by the width of the fitness function) of $\Lambda_f/\sqrt{V_S}, \Lambda_m/\sqrt{V_S} \lessapprox 0.5$.

Importantly, this indicates that for shift sizes up to this magnitude the analytics predict simulation outcomes accurately. Indeed, it is likely that we could relax the restriction on shift size slightly when the genetic architecture includes some alleles with sex-specific effects, since in this case sex-specific trait means are will move closer to their respective optima, and thus to the center of the respective fitness functions.

Of particular note is the fact that we do not observe an increase in sex-specific variances for shift sizes within this range, providing some confirmation of the prediction derived from Equation 8 that—when effect sizes are the same across sexes ($\phi_a = \pi/4$ or $5 \cdot \pi/4$), selection acts exactly symmetrically across sexes ($V_{S,f} = V_{S,m} = V_S$) and both sexes undergo exactly opposite shifts in optima $\Lambda_f = -\Lambda_m$—the effect of sex-specific directional selection on allele frequencies exactly cancel out and individual alleles experience zero net directional selection.

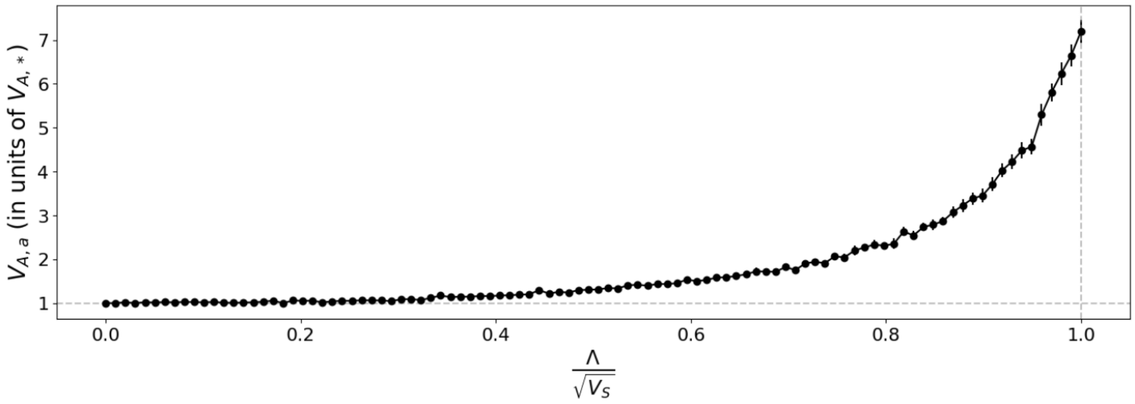

Figure S1: Exploring the limits to the shift size. The average of (asymptotic) sex-specific variances ($V_{A,a} = (V_{A,f} + V_{A,m})/2$ obtained from simulations) in units of the expected value of the average genetic variance ($V_{A,*}$ calculated using Equations 19 and 23) as a function of the shift size scaled by the width of the fitness function, $|\Lambda_f|/\sqrt{V_S}$ ($= |\Lambda_m|/\sqrt{V_S}$). When the normalized simulation results are close to 1 (the dashed grey line) it indicates that the analytic approximation for the average variance are accurate. The error bars represent 95% CI calculated as 1.96*SEM around the averages across 25 replicates.

### 4 Types of simulations

We wrote and compared three main types of simulations, varying in their assumptions and computational tractability to make sure that our analytical framework provides a reasonable approximation for the full model, in spite of its various simplifying assumptions. Documented code for these simulations can be found at https://github.com/gemmapuixeu/Puixeu_Hayward_2025.

First, we ran exact simulations, where we keep track of all individuals and the segregating alleles they carry. We defined two types of exact simulations realizing the full model described in the main text (Section 2.3): i) with fertility selection, where parents of the new generation are selected based on their fitness, and ii) with viability selection, where we accept or reject offspring generated from randomly-selected parents based on their fitness.

The second type of simulations, which we refer to as *Wright-Fisher* simulations, rely on the assumption of linkage equilibrium. In these simulations, we track sex-specific allele frequencies $(x_f, x_m)$ rather than individuals, and update them according to the Wright-Fisher process. To do this, we approximate the expected frequency any given allele, segregating at frequency $x_j$ with effect size $a_j$ in sex $j$, in the next generation $(x'_j)$ by

$$x'_j = x_j + E[\Delta x_j], \tag{S.32}$$

where $E[\Delta x_j]$ is calculated using Equation S.2 with the fitnesses, $\bar{w}^j_c$, of individuals with $c = 0, 1$ or 2 copies of the allele approximated by Equation S.4.

The third type of simulations, which we refer to as *Wright-Fisher Hardy-Weinberg* simulations, used for the results in the main text, additionally assumes Hardy-Weinberg equilibrium, meaning that the allele frequency differences between sexes after selection are negligible $(x_f = x_m = x)$. Thus, in these simulations, we need only track overall allele frequencies, $x$. In this case we use the fact that, manipulating terms in Equation S.2 and S.3, Equation S.1 can be expressed as

$$E[\Delta x] = \frac{x(1-x)}{2}\left(\left(1 - \frac{x_d^2}{4x(1-x)}\right)A + (1-2x)\left(1 + \frac{x_d^2}{4x(1-x)}\right)H\right), \tag{S.33}$$

with $x_d^2 \equiv (x_f - x_m)^2$ and $A$ (quantifying the strength of "additive selection") and

$H$ (quantifying the effect of dominance) defined as:

$$A = \frac{\bar{w}_2^f - \bar{w}_0^f}{2\overline{w}^f} + \frac{\bar{w}_2^m - \bar{w}_0^m}{2\overline{w}^m} \tag{S.34}$$

$$H = \frac{2\bar{w}_1^f - \bar{w}_2^f - \bar{w}_0^f}{2\overline{w}^f} + \frac{2\bar{w}_1^m - \bar{w}_2^m - \bar{w}_0^m}{2\overline{w}^m} \tag{S.35}$$

with

$$\overline{w}^j = \bar{w}_2^j + 2x\frac{\bar{w}_2^j - \bar{w}_0^j}{2} + 2x(1-x)\left(1 + \frac{x_d^2}{4x(1-x)}\right)\frac{2\bar{w}_1^f - \bar{w}_2^j - \bar{w}_0^j}{2}. \tag{S.36}$$

Consequently, taking $x_d = 0$ in Equations S.33 and S.36, we approximate the change in overall frequency by

$$E[\Delta x] = \frac{x(1-x)}{2}\left(A + (1-2x)H\right), \tag{S.37}$$

with

$$\overline{w}^j = \bar{w}_2^j + 2x\frac{\bar{w}_2^j - \bar{w}_0^j}{2} + 2x(1-x)\frac{2\bar{w}_1^f - \bar{w}_2^j - \bar{w}_0^j}{2}. \tag{S.38}$$

In the *Wright-Fisher Hardy-Weinberg* simulations, we use

$$x' = x + E[\Delta x], \tag{S.39}$$

and Equations S.37, S.34, S.35 and S.38 to update overall allele frequencies each generation according to a Wright-Fisher process. We approximate sex-specific fitnesses of individuals with $c = 0, 1$ or $2$ copies of the allele by setting $x_j = x$ in Equation S.4, yielding

$$\bar{w}_c^j \approx \mathrm{Exp}\left[-\frac{[D_j - a_j(c-2x)]^2 \cdot \gamma_j^2}{V_S}\right]. \tag{S.40}$$

As Figure S2 shows, all of our simulation types yield an average genetic variance at equilibrium under stabilizing selection that is close to the analytic prediction. In the multigenic case, $V_{A,a}$ obtained from exact simulations slightly exceeds the analytic prediction, likely owing to minor deviations from linkage equilibrium.

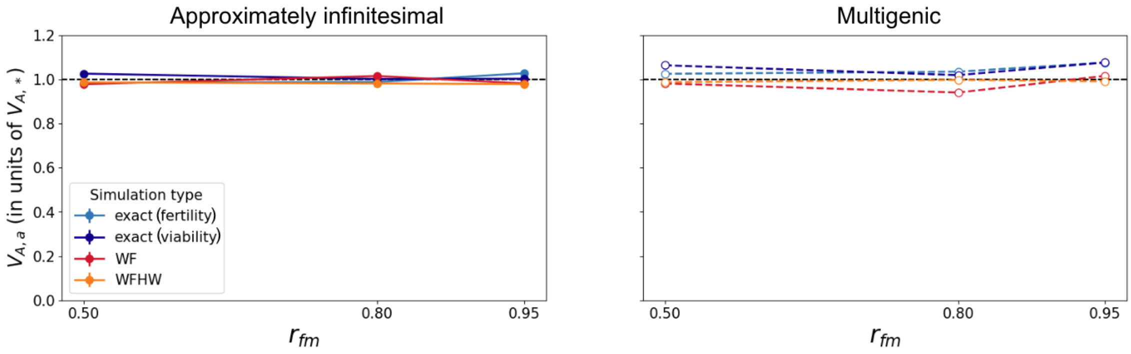

Figure S2: The various simulation types match our analytic predictions. The figure displays the average genetic variance ($V_{A,a} = (V_{A,f} + V_{A,m})/2$ in units of the expected value of the average genetic variance ($V_{A,*}$ calculated using Equations 19 and 23) for an approximately infinitesimal (filled solid, left) and multigenic (open dashed, right) genetic architectures, three values of $r_{fm}$ and for each of our simulation types: exact simulations with fertility (light blue) and viability (dark blue) selection, as well as *Wright-Fisher* (WF, red) and *Wright-Fisher Hardy-Weinberg* (WFHW, orange) simulations. Error bars represent 95% CI calculated as 1.96· SEM across 100 replicates.

### 5 Empirical calculation of sex-specific variances, intersex covariance and intersex correlation

When running *Wright-Fisher Hardy-Weinberg* simulations we can calculate sex-specific variances, intersex covariance and intersex correlation from allele frequencies, following equations 4, 5 and 3, respectively. However, when dealing with empirical data (or individual-based simulations, where we keep track of all individuals in the population and the mutations they carry), we need different, empirical, measures of these (co)variances.

Empirical sex-specific variances ($V^e_{A,f}$ and $V^e_{A,m}$) can be computed as the variance across all individuals of each sex in the population:

$$V^e_{A,f} = \frac{\sum_i^{N_f} (z_{i,f} - \bar{z}_f)^2}{N_f} \tag{S.41}$$

$$V^e_{A,m} = \frac{\sum_i^{N_m} (z_{i,m} - \bar{z}_m)^2}{N_m}, \tag{S.42}$$

which, under our assumptions of linkage equilibrium and an additive trait with no environmental contribution, should correspond to the sex-specific genic variances in equation 4.

The intersex covariance, $B$, and correlation, $r_{fm}$, are defined as the covariance and correlation between allelic effects if they were to be expressed in both sexes at the same time (Equations 5 and 3). Since genotypes are never expressed simultaneously in both a female and male (except in the case of heavily inbred populations), the empirical calculation of intersex covariance and correlation relies on phenotypic similarity between relatives. For example, one can use a parent-to-offspring regression to calculate empirical intersex correlation (Lynch & Walsh, 1998; as in e.g. Bonduriansky & Rowe, 2005) as:

$$r^e_{fm} = \sqrt{\frac{h^2_{MS} h^2_{FD}}{h^2_{MD} h^2_{FS}}}, \tag{S.43}$$

where $h^2$ represents heritability, calculated as twice the offspring-to-parent phenotypic regression coefficient, for mother-son (MS), father-daughter (FD), mother-daughter (MD) and father-son (FS). We compute empirical between-sex covariance from empirical sex-specific variances and empirical intersex correlation as

$$B^e = \frac{\sqrt{V^e_{A,f} V^e_{A,m}}}{r^e_{fm}}. \tag{S.44}$$

#### 6 Calculation of the total genetic variance, $V_{A,t}$

Empirically, besides the sex-specific variances, which can be computed following equations S.41 and S.42, we can compute the total variance across all individuals in the population as

$$V^e_{A,t} = \frac{\sum_i^N (z_i - \bar{z})^2}{N}. \tag{S.45}$$

However, to compute total variance from *Wright-Fisher* and *Wright-Fisher Hardy-Weinberg* simulations we need to compute the total variance as the sum of the within-sex and between-sex variance: $V_{A,t} = V_{A,w} + V_{A,B}$, where $V_{A,w} = \frac{SSW}{N}$ , $V_{A,b} = \frac{SSB}{N}$.

The sum of squares within (SSW) is the sum of both sex-specific sums of squares:

$$SSW = \sum_{j\in\{m,f\}} \sum_{i=1}^{N_j} (z_{i,j} - \bar{z}_j)^2 \tag{S.46}$$

$$= \sum_{j\in\{m,f\}} \sum_{i=1}^{N/2} (z_{i,j} - \bar{z}_j)^2 \tag{S.47}$$

$$= N/2 \cdot \frac{\sum_{i=1}^{N/2} (z_{i,f} - \bar{z}_f)^2}{N/2} + N/2 \cdot \frac{\sum_{i=1}^{N/2} (z_{i,m} - \bar{z}_m)^2}{N/2} \tag{S.48}$$

$$= N \cdot \frac{(V_{A,f} + V_{A,m})}{2}, \tag{S.49}$$

where $z_{i,j}$ is the trait value of individual $i$ with sex $j = f$ or $m$, and $\bar{z}_f$ and $\bar{z}_m$ are sex-specific trait means. From Equations S.46 to S.49 it is clear that, since we assumed an equal number of males and females, the SSW corresponds to $N$ times the average of sex-specific variances (equivalent to $V_{A,a}$, following Equation 39), for which we have genic formulae (Equation 4). The sum of squares between (SSB) can also be calculated using genic formulae, since

$$SSB = \sum_{j\in\{m,f\}} \frac{N}{2} \cdot (\bar{z}_j - \bar{z})^2 = \frac{N}{2} \cdot [(\bar{z}_f - \bar{z})^2 + (\bar{z}_m - \bar{z})^2] \tag{S.50}$$

and sex-specific trait means, $\bar{z}_f$ and $\bar{z}_m$, can be calculated from the contributions across mutations (as can $\bar{z}$, their average).

To summarize, it follows from Equations S.49 and S.50, that

$$V_{A,w} = \frac{1}{2}(V_{A,f} + V_{A,m}) = V_{A,a} \text{ and} \tag{S.51}$$

$$V_{A,b} = \frac{1}{2} \cdot [(\bar{z}_f - \bar{z})^2 + (\bar{z}_m - \bar{z})^2], \tag{S.52}$$

allowing us to calculate

$$V_{A,t} = V_{A,w} + V_{A,b} \tag{S.53}$$

even in *Wright-Fisher* and *Wright-Fisher Hardy-Weinberg* simulations when we do not keep track of individuals.

### 7 The relationship between various measures of sexual dimorphism and $r_{fm}$ at equilibrium

#### 7.1 The relationship between $SD$ (signed and absolute) and expected intersex correlation

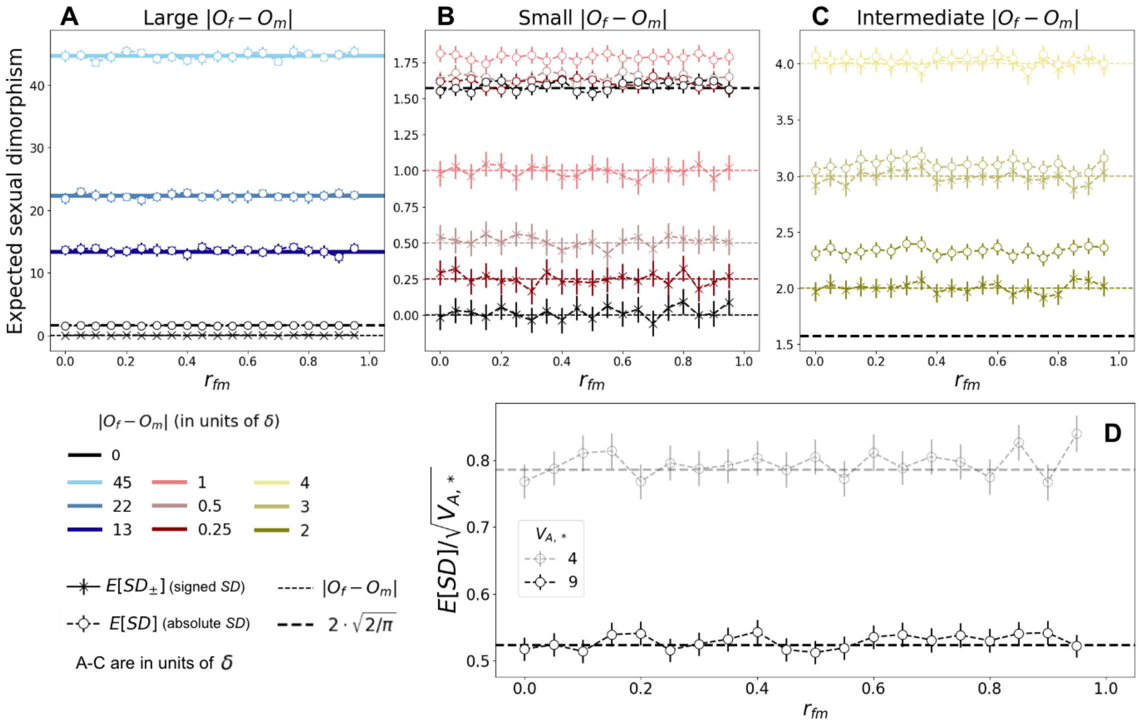

Figure S3: Relationship between expected intersex correlation ($r_{fm}$) and sexual dimorphism at equilibrium with a multigenic genetic architecture. A-C: Expected sexual dimorphism, signed (as the difference between sex-specific trait means, Equation 7; crosses) and absolute (as the absolute difference between sex-specific trait means, Equation 6; circles) across $r_{fm} \in [0, 1)$, with $V_{A,*} = 9$ and for various $|O_f - O_m|$ ranges: large, with $|O_f - O_m| > 10$ (A); small, with $|O_f - O_m| \in [0, 1]$ (B); intermediate, $|O_f - O_m| \in [2, 4]$ (C). The thick black dashed line corresponds to $E[SD]$ predicted by Equation 33. D: Expected (absolute) sexual dimorphism, scaled by the standard deviation in sex-specific trait distributions, $E[SD]/\sqrt{V_{A,*}}$, for $O_f = O_m = 0$ and genetic variances $V_{A,*} = 4$ (semi-transparent) and 9 (opaque). Simulations were run for $100N$ generations, except for $r_{fm} \geq 0.8$, which were run for $500N$ generations. Markers and error bars indicate estimates and 95% CIs calculated as 1.96·SEM across 2,000 replicates.

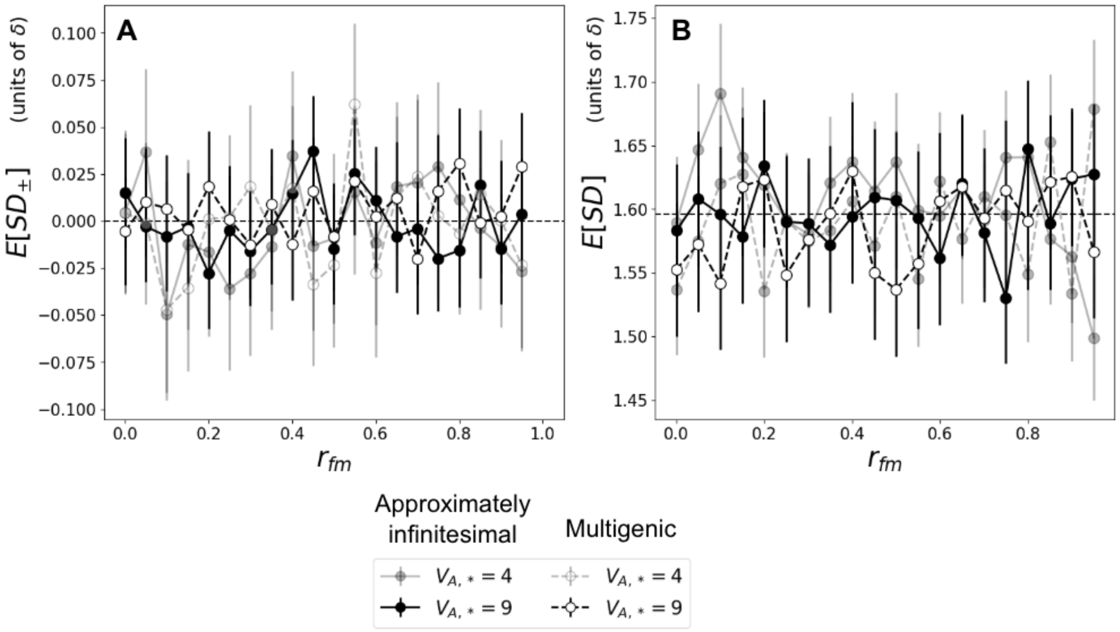

Figure S4: Relationship between expected intersex correlation ($r_{fm}$) and signed sexual dimorphism (Equation 7, in A) and absolute sexual dimorphism (Equation 6, in B) at equilibrium, for an approximately infinitesimal (filled solid) and multigenic (open dashed) genetic architectures, and for $V_{A,*} = 4$ (semi-transparent) and 9 (opaque). The horizontal dashed line in B corresponds to the prediction $E[SD] = 2 \cdot \sqrt{2/\pi}$ (Equation 33). Simulations were run Markers and error bars indicate averages and 95% CIs calculated as 1.96·SEM across 2,000 replicates.

#### 7.2 Sexual dimorphism with an $r_{fm} = 1$

Figure S5 shows the signed ($SD_{\pm}$, A) and absolute ($SD$, B) sexual dimorphism at equilibrium. As we predict, $SD_{\pm} = 1$ for all differences between sex-specific optima. $SD$, however, is slightly larger than 0, even when all mutations have identical effects between the sexes. This is due to the fact that new incoming mutations arise in a sex-specific manner, so they generate a slight sexual dimorphism even for $r_{fm} = 1$. This is effect is stronger with a multigenic genetic architecture, since there are more large-effect mutations.

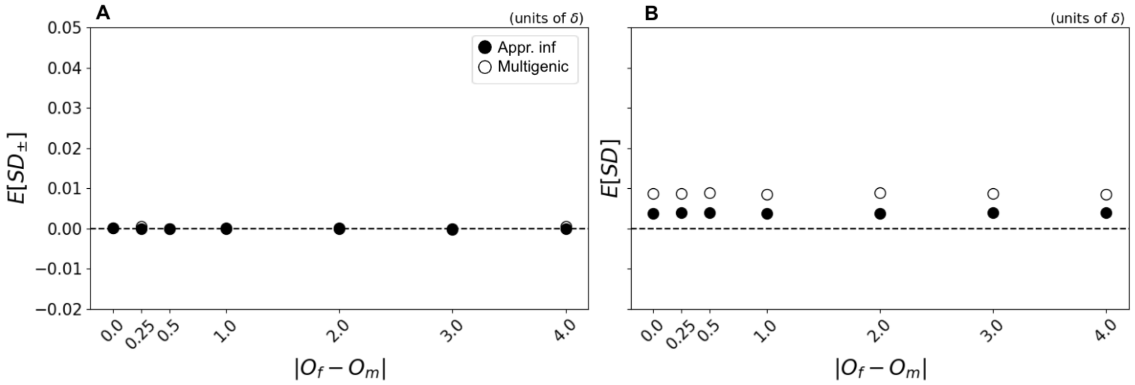

Figure S5: Sexual dimorphism for $r_{fm} = 1$. Signed sexual dimorphism (Equation 7, in A) and absolute sexual dimorphism (Equation 6, in B) at equilibrium, for an approximately infinitesimal (solid circles) and multigenic (open circles) genetic architectures for various $|O_f - O_m|$ (representing the various scenarios in Figure 1B,C), and for $V_{A,*} = 9$ and $r_{fm} = 1$. Simulations were burned-in for $100N$ generations. Markers and error bars indicate averages and 95% CIs calculated as 1.96·SEM across 2,000 replicates.

#### 7.3 Fluctuations in sex-specific trait means and $SD$ at equilibrium

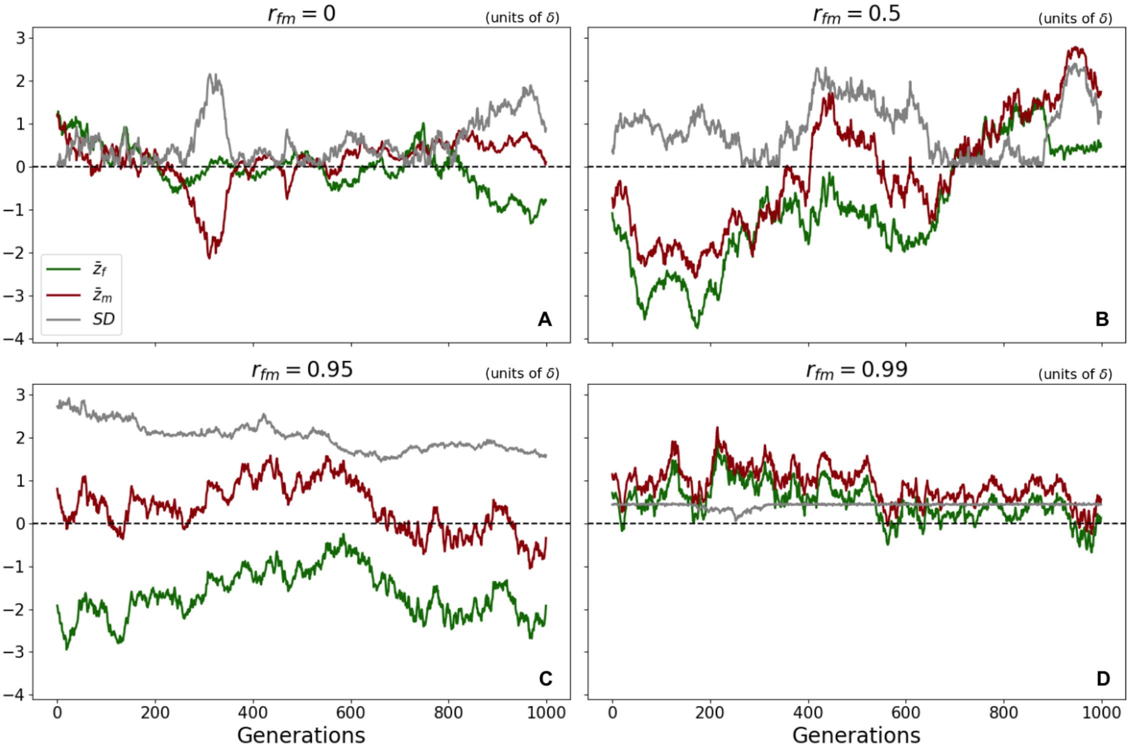

Figure S6: Sex-specific trait means and (absolute) $SD$ across time for various levels of $r_{fm}$: 0 (A), 0.5 (B), 0.95 (C) and 0.99 (D), with a multigenic genetic architecture and $V_{A,*} = 9$. Simulations have been burnt-in for $500N$ (except for D, which was burnt-in for $1000N$ generations. Results for a single replicate are displayed.

#### 7.4 Comparison of the distribution of $SD_{\pm}$ with that of a Gaussian

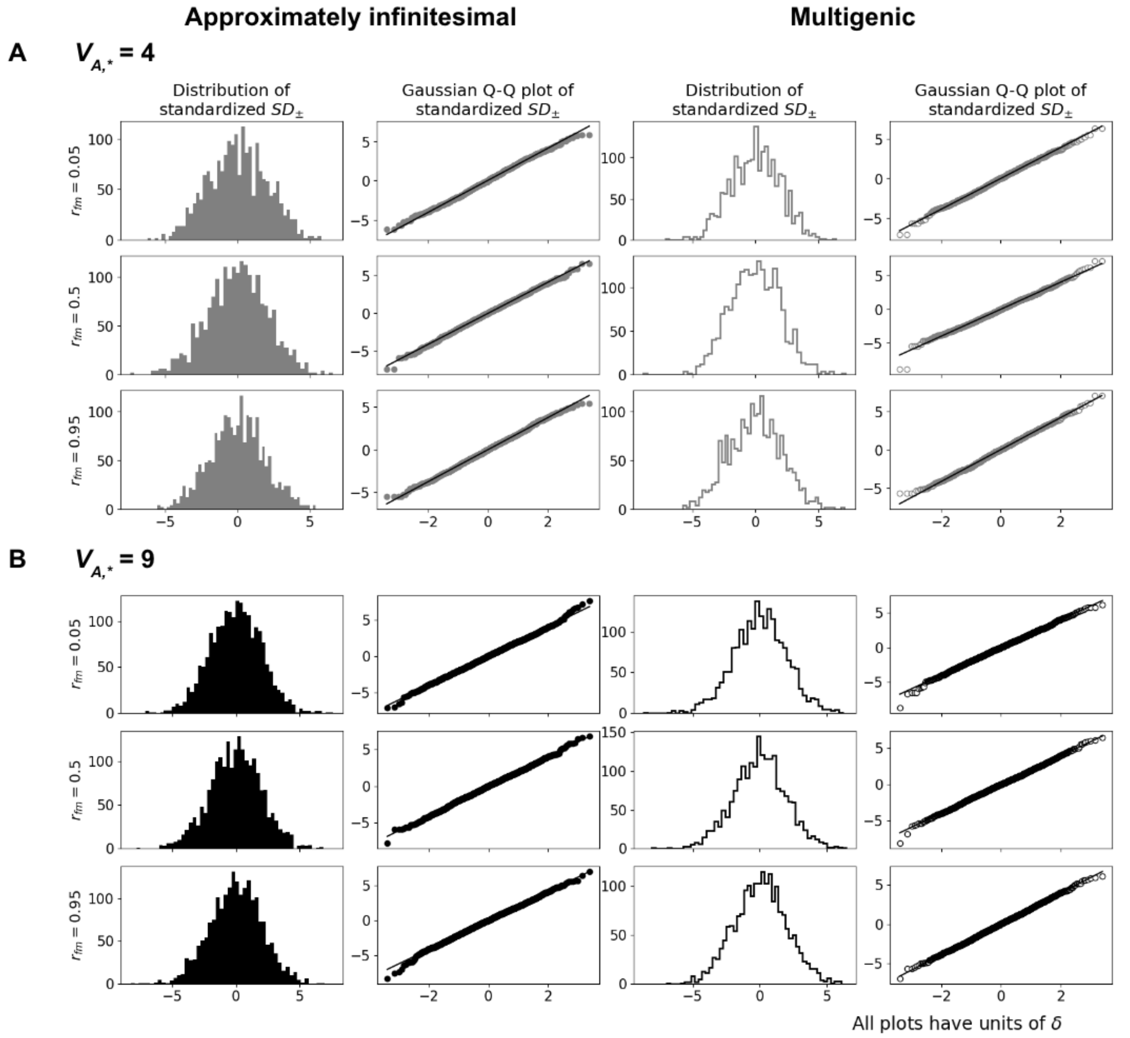

Figure S7: The distribution of $SD_{\pm}$ is approximately Gaussian for a wide range of $r_{fm}$ and both an approximately infinitesimal and multigenic genetic architecture. The panels display the distribution (first and third columns) and Q-Q plots against a Gaussian distribution (second and fourth columns) of standardized $SD_{\pm} = (SD_{\pm} - E[SD_{\pm}])/\sqrt{Var[SD_{\pm}]}$ for genetic variances $V_{A,*} = 4$ (A, grey) and 9 (B, black) and approximately infinitesimal ($E(a^2) = 1$; filled) and multigenic ($E(a^2) = 16$; open) genetic architectures.

#### 7.5 Characterization of the relationship between $V[SD]$ and $r_{fm}$ due to drift

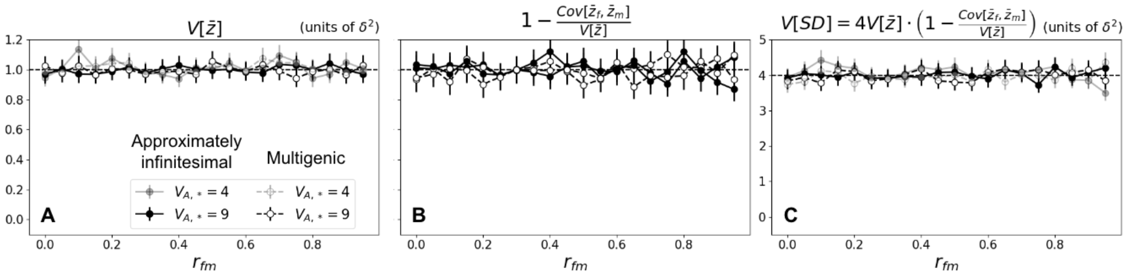

Figure S8: Characterization of the relationship between $V[SD]$ and $r_{fm}$. Contributions of $V[\bar{z}]$ (B) and $1 - Cov[\bar{z}_f, \bar{z}_m]/V[\bar{z}]$ (C) to $V[SD]$ (D), according to Equation 28, for an approximately infinitesimal (filled solid) and multigenic (open dashed) genetic architectures, and for $V_{A,*} = 4$ (semi-transparent) and 9 (opaque). In B we see that $1 - Cov[\bar{z}_f, \bar{z}_m]/V[\bar{z}] \approx 1$ for all values of $r_{fm}$ (implying that $Cov[\bar{z}_f, \bar{z}_m] \approx 0$) and $V[SD]$ is completely determined by $V[\bar{z}]$. Markers and error bars indicate estimates and 95% CIs across 2,000 replicates calculated as follows: for variances (A,C), using Chi-squared distribution. For the between-sex covariance, $Cov[\bar{z}_f, \bar{z}_m] = E[\bar{z}_f \cdot \bar{z}_m] - E[\bar{z}_f]E[\bar{z}_m]$. Given that, in our case, $E[\bar{z}_f] = E[\bar{z}_m] = 0$, we can estimate the covariance as $E[\bar{z}_f \cdot \bar{z}_m]$ and obtain 95% CI as 1.96·standard error of the mean (SEM). For $Cov[\bar{z}_f, \bar{z}_m]/V[\bar{z}]$ (B), our estimates are of form $E[B]/E[C]$, for which we were able to approximate CIs ($[l, u]$ where $l = E[B]/E[C] - s_l$ and $u = E[B]/E[C] + s_u$ are the lower and upper bounds of the CI) by first computing CIs for the numerator and denominator, and then combining them to obtain $s_l = E[B]/E[C] \cdot \sqrt{(s_{l_B}/E[B])^2 + (s_{l_C}/E[C])^2}$ (and similarly to get the upper bound).

### 8 Deviations in phenotypic evolution with a multigenic genetic architecture

With an approximately infinitesimal genetic architecture (light lines in Figure S9) 2$^{nd}$ and 3$^{rd}$ central moments of the phenoytpic distribution remain at the equilibrium values during directional selection after a shift in optima. In contrast, with a multigenic genetic architecture, selection in the rapid phase drives an increase in 2$^{nd}$ and 3$^{rd}$ central moments of the phenoytpic distribution (dark lines Figure S9). Concretely, after a shift in optima of the same magnitude and direction between sexes (concordant adaptation) we see an increase in overall variance (Figure S9A), which is driven by an increase in frequency of shared (Figure S9B) rather than sex-specific (Figure S9C) mutations, and this generates an increase of the intersex covariance (Figure S9E). After a shift in optima of same magnitude and opposite direction between the sexes (dimorphic adaptation) we also see an increase in overall variance (Figure S9G), which in this case is driven by an increase in frequency of sex-specific (Figure S9I) rather than shared (Figure S9H) mutations. Consequently, it does not generate an increase of intersex covariance (Figure S9K). We also see that with both concordant and dimorphic adaptation there is an increase in the skewness of the phenotypic distribution (3$^{rd}$ central moments, Figure S9F,L) with a multigenic architecture. However, even with a multigenic genetic architecture, the difference in sex-specific variances ($V_{A,d}$, Figure S9D,J) remains low in both scenarios.

The changes described above in the 2$^{nd}$ and 3$^{rd}$ central moments in the multigenic case lead to deviations in the phenotypic dynamics with respect to an infinitesimal architecture. In particular, concordant adaptation tends to be accelerated by changes in the 2$^{nd}$ central moments during the rapid phase (reflected by the slightly more rapid decay in $D_a$ in Figure S10, top), and to be delayed by changes in the 3$^{rd}$ central moments during equilibration (reflected in the quasi-static approximation for $D_a$ in Figure S11, top, which we describe in Section 8.1). Dimorphic adaptation is similarly accelerated by changes in the 2$^{nd}$ central moments during the rapid phase with a multigenic architecture (reflected by the faster decay in $D_d$ during dimorphic adaptation in Figure S10, bottom), and delayed by changes in the 3$^{rd}$ central moments during equilibration (reflected in the quasi-static approximation for $D_d$ which we describe in Section 8.1). The accelerated decay due to changes in variance have a particularly strong effect with high $r_{fm}$ (as we illustrate for $r_{fm} = 0.95$)

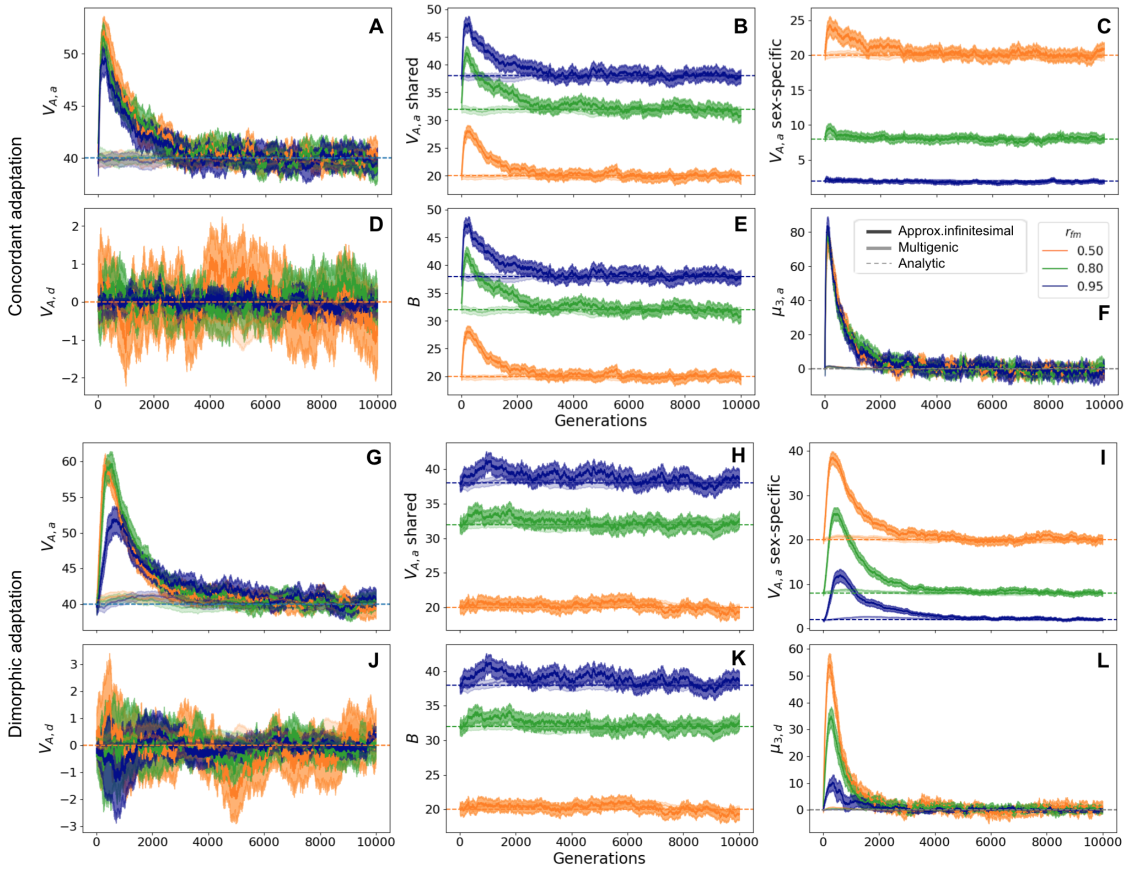

Figure S9: 2$^{\text{nd}}$ and 3$^{\text{rd}}$ central moments of the phenotypic distribution along time. Top and bottom panels correspond to evolution after sexually-concordant and sexually-dimoprhic shifts in optima, respectively corresponding to scenarios illustrated in main Figure 2A ($\Lambda_a = 0.25\sqrt{V_S}$ and $\Lambda_d = 0$) and 2B ($\Lambda_a = 0$ and $\Lambda_d = 0.25\sqrt{V_S}$). A, G: Average genetic variance, $V_{A,a} \equiv 0.5\cdot(V_{A,f}+V_{A,m})$. B, H: Variance contributed by shared mutations, which exactly corresponds to the intersex covariance (in E, K). C, I: Variance contributed by sex-specific mutations. D,J: Average difference genetic variance $V_{A,d} \equiv 0.5\cdot(V_{A,f}-V_{A,m})$. E, K: Between-sex covariance, which exactly corresponds to the variance contributed by shared mutations. F and L: Average and average difference third central moments of the phenotypic distribution, respectively. Only the relevant 3$^{\text{rd}}$ central moment is shown, as $\mu_{3,d}$ ($\mu_{3,a}$) remains zero for sexually-concordant, top (sexually-dimorphic, bottom) adaptation. Orange, green and blue indicate simulations with different intersex correlations ($r_{fm} = 0.5, 0.8, 0.95$, respectively). Simulations with multigenic ($E(a^2) = 16$) and approximately infinitesimal ($E(a^2) = 1$) genetic architecture are respectively plotted in bright and dim colors. Dashed horizontal lines correspond to the equilibrium values. The shaded ares correspond to 95% CIs around averages across 200 replicates.

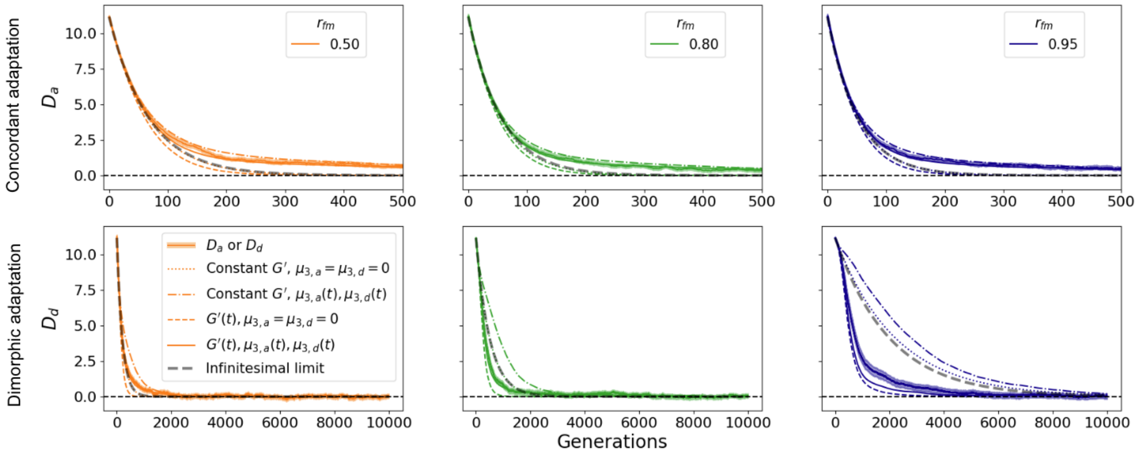

Figure S10: Phenotypic evolution in the multigenic case. Top row corresponds to $D_a$ along time, for optima shift $\Lambda_a = 0.25\sqrt{V_S}$ and $\Lambda_d = 0$ leading to sexually-concordant adaptation (the scenario depicted in Figure 2A). Bottom row corresponds to $D_d$ along time, for optima shift $\Lambda_a = 0$ and $\Lambda_d = 0.25\sqrt{V_S}$, leading to sexually-dimoprhic adaptation (the scenario depicted in Figure 2B). Thick lines with shaded areas correspond to simulations with multigenic genetic architecture (with shaded areas representing the 95% CIs around averages across 200 replicates), dashed grey lines to the prediction in the infinitesimal limit (Equation 42), and various types of thin lines to predictions using various versions of Equation 47: dotted, with constant G' matrix and with $\mu_{3,a} = \mu_{3,d} = 0$, effectively coinciding with the prediction assuming infinitesimal architecture; dash-dotted, with constant G' matrix and with $\mu_{3,a}, \mu_{3,d}$ updated generation-wise according to their values in the simulations; dashed, with elements in the G' matrix updated generation-wise according to their values in the simulations and with $\mu_{3,a} = \mu_{3,d} = 0$; solid, with both $2^{\text{nd}}$ and $3^{\text{rd}}$ order moments updated generation-wise according to their values in the simulations (which most closely match the simulations). First, second and third columns (orange, green and blue) correspond to simulations with $r_{fm} = 0.5, 0.8$ and $0.95$. We have zoomed in the results for $D_a$ by depicting a shorter timescale, so that the relevant dynamics can be appreciated for all cases.

#### 8.1 The quasi-static approximation

With an approximately infinitesimal genetic architecture, $D_a$ and $D_d$ are essentially zero after the rapid phase. In contrast, with a multigenic genetic architecture, $D_a$ and $D_d$ can remain some small distance away from zero for many generations after the rapid phase (Figure S10, top). In this case, to describe the dynamics of $D_a$ and $D_d$ during equilibration, we derive a quasi-static approximation similar to that derived for a single sex in Hayward and Sella (2022). This approximation expresses the distances from the optimum as functions of the second and third moments. It can be obtained by setting $E[\Delta D_a] \approx 0$ and $E[\Delta D_d] \approx 0$ in Equation 47 which, after working through the algebra, yields:

$$\begin{aligned} D_a^*(t) \approx & \frac{\mu_{3,a}(t)}{V_{A,a}(t) + B(t)} \cdot (1 - \xi(t)) - \frac{\mu_{3,d}(t)}{V_{A,d}(t)} \cdot \xi(t) \\ D_d^*(t) \approx & \frac{\mu_{3,d}(t)}{V_{A,a}(t) - B(t)} \cdot (1 - \xi(t)) - \frac{\mu_{3,a}(t)}{V_{A,d}(t)} \cdot \xi(t) \end{aligned} \quad \text{(S.54)}$$

where

$$\xi(t) \equiv \frac{V_{A,d}^2(t)}{V_{A,a}^2(t) - B^2(t) - V_{A,d}^2(t)}$$

When things are perfectly symmetric between the sexes, the above simplifies further since $V_{A,d}(t)$ should remain zero after the shift. In that case we get

$$D_a^{**}(t) \approx \frac{\mu_{3,a}(t)}{V_{A,a}(t) + B(t)}; \quad D_d^{**}(t) \approx \frac{\mu_{3,d}(t)}{V_{A,a}(t) - B(t)} \quad \text{(S.55)}$$

As Figure S11 shows, in the infinitesimal case at $t_a$ and $t_d$ (the end of rapid phase, determined using Equation 43) sex-specific means have matched their optima $D_a, D_d \approx 0$. However, with multigenic genetic architecture there is a slower equilibration phase for $D_a$, while for $D_d$ equilibration (like overall adaptation) is similar to (faster than) in the infinitesimal case for low (high) $r_{fm}$. Both these dynamics are well predicted by the quasi-static approximation (Equations S.55).

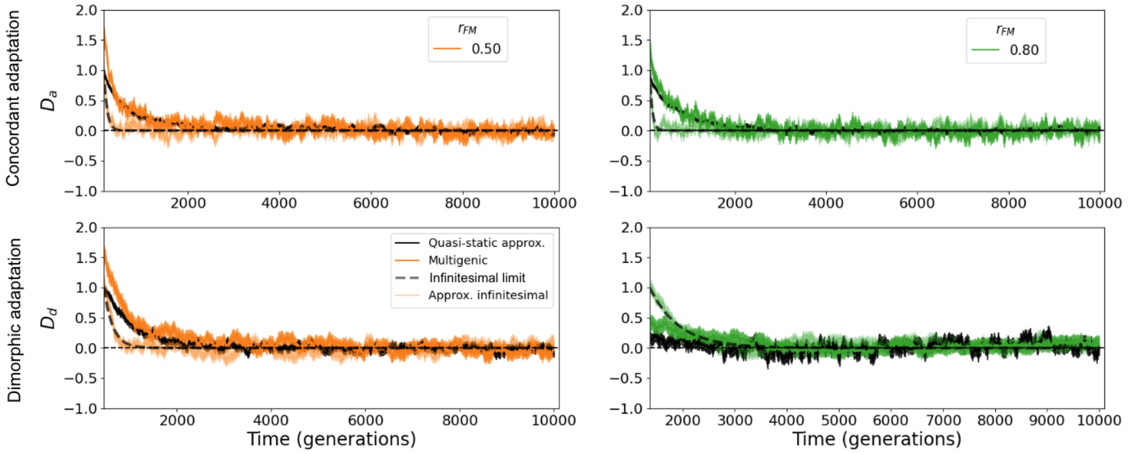

Figure S11: Quasi-static approximation during equilibration. Evolution of $D_a$ and $D_d$ (first and second rows), for sexually-concordant and sexually-dimorphic adaptation after shifts in optima corresponding to $\Lambda_a = 0.25\sqrt{V_S}$, $\Lambda_d = 0$ and $\Lambda_a = 0$, $\Lambda_d = 0.25\sqrt{V_S}$ (scenarios depicted in Figure 2A and 2B, respectively) for $r_{fm} = 0.5, 0.8$ (first and second columns, in orange and green) with multigenic ($E(a^2) = 16$, dark) and approximately infinitesimal ($E(a^2) = 1$, light) genetic architecture following equation 47. Black solid lines correspond to the quasi-static approximation. Black dashed lines correspond to the evolution in $D_a, D_d$ in the infinitesimal case (Equation 42). The starting point of the current plots corresponds to the time where $D_a$ and $D_d$ reach the typical deviation of the population mean from the optima at equilibrium, $\delta = \sqrt{V_S/2N} = 1$, which determines the end of the rapid phase and beginning of equilibration ($t_a, t_d$ in Equation 43). The error bars represent 95% CIs across 200 replicates.

#### 8.2 Transient increase in variances under directional selection

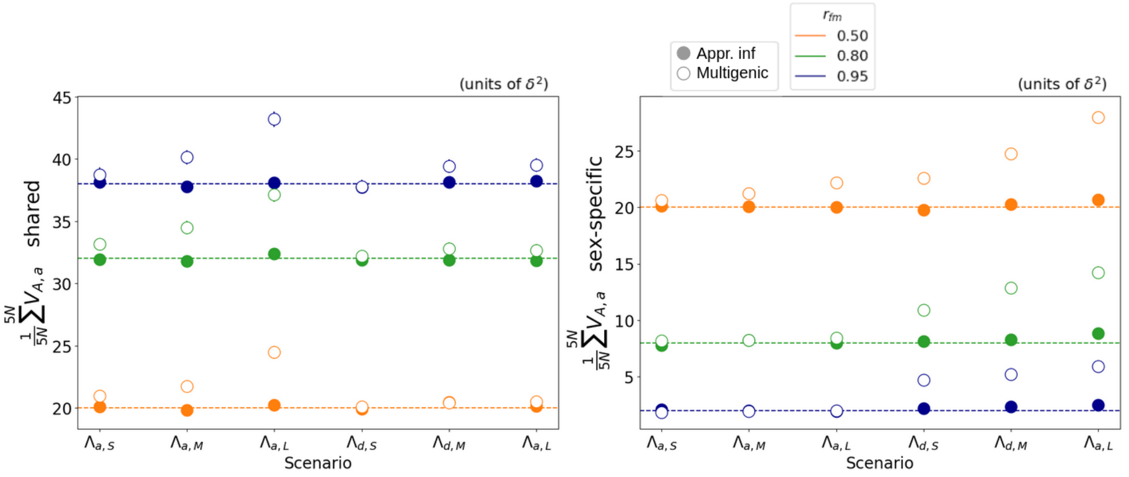

Figure S12: Transient increase in shared (left) and sex-specific (right) variances during adaptation under directional selection. Means and 95% CIs of average genetic variance ($V_{A,a}$) contributed by shared (left) and sex-specific (right) mutations across $5N$ generations after the shift in optima ($\equiv$ empirical integrals during rapid phase of adaptation). Results are shown for approximately infinitesimal (solid circles) and multigenic (open circles) genetic architectures and across different scenarios indicating different types of shifts: $\Lambda_{a,\_}$ are shifts of same magnitude and direction in both sexes, leading to sexually-concordant adaptation (similar to the scenario depicted in Figure 2A, in which $\Lambda_{d,\_} = 0$); $\Lambda_{d,\_}$ are shifts of same magnitude and different direction in both sexes, leading to sexually-dimorphic adaptation (similar to the scenario in Figure 2B, in which $\Lambda_{a,\_} = 0$). $\Lambda_{\_,S}$, $\Lambda_{\_,M}$ and $\Lambda_{\_,L}$ indicate small, medium and large shifts, with magnitudes $0.15\sqrt{V_S}$, $0.25\sqrt{V_S}$ and $0.5\sqrt{V_S}$, respectively. The colors indicate results for different $r_{fm}$ values: 0.5, 0.8 and 0.95 in orange, green and blue.

### References

Bonduriansky, R., & Rowe, L. (2005, September). Intralocus sexual conflict and the genetic architecture of sexually dimorphic traits in Prochyliza xanthostoma (Diptera: Piophilidae). *Evolution; International Journal of Organic Evolution*, *59*(9), 1965–1975.

Hayward, L. K., & Sella, G. (2022, September). Polygenic adaptation after a sudden change in environment. *eLife*, *11*, e66697. doi: 10.7554/eLife.66697

Kidwell, J. F., Clegg, M. T., Stewart, F. M., & Prout, T. (1977, January). Regions of stable equilibria for models of differential selection in the two sexes under random mating. *Genetics*, *85*(1), 171–183. doi: 10.1093/genetics/85.1.171

Lynch, M., & Walsh, B. (1998). *Genetics and Analysis of Quantitative Traits* (1st edition ed.). Sunderland, Mass: Sinauer Associates is an imprint of Oxford University Press.
